# Intracellular *C. neoformans* infection stimulates increased glycolytic activity in fetal liver-derived alveolar-like macrophages

**DOI:** 10.1101/2025.11.28.690224

**Authors:** Derek A. Wiggins, Josh B. Griggs, Andrew M. England, Emily N. Callison, Cedra H. Kamel, Cathrine A. Hasan, Anna E. Davis, Morgan E. Chappell, Kayla N. Conner-Halim, Andrew J. Olive, Erin E. McClelland, Rachel N. Leander, Rebecca L. Seipelt-Thiemann, David E. Nelson

**Affiliations:** Department of Biology, Middle Tennessee State University, Murfreesboro, TN 37132; Department of Mathematical Sciences, Middle Tennessee State University, Murfreesboro, TN 37132; Veterinary Diagnostic Laboratory, Michigan State University, Lansing, MI 48910; Department of Microbiology and Molecular Genetics, Michigan State University, East Lansing, MI 48824; Wood College of Osteopathic Medicine, Marian University, Indianapolis, IN 46222

**Keywords:** Macrophage, alveolar macrophage, macrophage polarization, *Cryptococcus neoformans*, interferon-gamma, HIF-1

## Abstract

Alveolar macrophages (AMs) serve as a first line of defense against respiratory pathogens, including *Cryptococcus neoformans*, the primary causative agent of cryptococcosis, a deadly pulmonary mycosis which commonly afflicts immunocompromised individuals. While these innate immune cells are thought to play a pivotal role in controlling the outcome of *C. neoformans* infections, this critical host-pathogen interaction is more commonly studied *in vitro* using bone marrow-derived macrophages (BMDM) or immortalized macrophage cell lines that differ in ontogeny and phenotype from AMs. In this work, we characterized fetal liver-derived alveolar-like macrophages (FLAMs) as an alternate model to study the earliest stages of *C. neoformans* infection. Here, we show that the FLAM steady state transcriptome is more similar to primary AMs than peritoneal macrophages and the macrophage cell lines, RAW264.7 and J774, and that FLAMs exhibit distinct transcriptional responses to IFNγ stimulation and *C. neoformans* infection compared to J774 cells. Specifically, transcriptome profiling and gene ontology analysis indicate that *C. neoformans* infection of FLAMs, but not J774 cells, increases the expression of canonical glycolytic genes, including *Slc2a1, Pgk1, and Ldha*, which is accompanied by a metabolic shift favoring glycolysis. Furthermore, activation or inhibition of hypoxia inducible factor 1 (HIF1) activity utilizing dimethyloxalylglycine (DMOG) and echinomycin, respectively, indicates that the expression of select glycolytic genes in *C. neoformans*-infected FLAMs is HIF1-dependent. Collectively, our results suggest that FLAMs serve as an appropriate tool for modeling AM:*C. neoformans* interactions and investigating the effects of this pathogen on host AM immunometabolism.

## Introduction

The facultative intracellular pathogen, *Cryptococcus neoformans*, is the main causative agent of cryptococcosis. This fatal fungal infection that typically starts in the lungs before disseminating to the central nervous system, and is responsible for an estimated 152,000 cases of cryptococcal meningitis (CM) and ∼112,000 deaths per year (1). As an opportunistic mycosis, cryptococcosis disproportionately affects individuals who are severely immunocompromised, with the majority of CM cases occurring in patients afflicted with late-stage or advanced acquired immunodeficiency syndrome (AIDS), permitting extrapulmonary disease progression (2). However, in immunocompetent individuals, *C. neoformans* rarely disseminates from the lung or causes serious disease. Rather, when *C. neoformans* propagules are inhaled into the alveoli of the lung, they encounter alveolar macrophages (AMs), which eliminate fungal cells and restrict their spread throughout the body. In this way, these innate immune cells operate as a first line of defense against the pathogen and are essential for the successful control of the infection (3). As many of the current drug treatments for cryptococcosis are fungistatic and often carry serious side effects, and with cases of drug-resistant *C. neoformans* infections on the rise (4–7), improving our understanding of the *C. neoformans*:AM interaction may provide vital information for the development of new host-directed therapies. However, progress in this area has been hampered by a lack of suitable *in vitro* models.

While *C. neoformans*:macrophage interactions have been investigated extensively over the last two decades, these studies have typically employed cell models that represent myeloid-derived macrophages, including bone marrow-derived macrophages (BMDMs), the cancer-derived macrophage-like cell lines, J774 and RAW264.7, or monocyte cell lines, like THP-1, that can be differentiated into macrophages (8–13). AMs, however, have a distinct ontology and do not originate from the myeloid lineage (14–17). Instead, AMs are derived from yolk sac macrophages and migrating hepatic monocytes, which seed the alveoli during embryonic development and differentiate into mature AMs following exposure to the cytokines granulocyte-macrophage colony-stimulating factor (GM-CSF) and transforming growth factor beta (TGF-β) (18,19).

These self-sustaining, multifunctional cells are distinct from other macrophage types in several ways. They are uniquely adapted to the high-oxygen and low-glucose environment of the lung, favoring mitochondrial respiration over glycolysis (20,21). They catabolize surfactant, which helps to maintain surfactant homeostasis in the alveoli (22,23). They are also characterized as immunosuppressive or hypoinflammatory; they do not conform to the M1/M2 paradigm of macrophage polarization, expressing M1 markers at low levels or adopting intermediate polarization states, possibly to avoid potentially harmful inflammation that could compromise lung function (24–27).

Generally obtained from mice, primary AMs are difficult to isolate in high numbers, fail to replicate efficiently, and lose AM characteristics after prolonged maintenance in cell culture (28), complicating their use for modeling AM:*C. neoformans* interactions. To circumvent these issues, *Thomas et al* recently developed a new AM model by differentiating cultured murine fetal liver monocytes with GM-CSF and TGF-β (29). These cells, which Thomas *et al* named ‘fetal liver-derived alveolar-like macrophages’ (FLAMs), exhibit a stable AM-phenotype for at least 40 days in culture, expressing the lineage-specific AM surface markers, Siglec-F and CD11c, and various AM-identifying genes, including *Marco*, *Pparg*, and *Tgfbr1*.

In this study, we evaluated FLAMs as a potential new model for the investigation of AM:*C. neoformans* interactions. We provide substantial evidence that the steady state transcriptome of FLAMs is more similar to primary AMs than peritoneal macrophages (PMs) or macrophage cell lines, supporting their use as a surrogate AM model, and show that these cells exhibit a contrasting transcriptional and metabolic response to IFNγ exposure and intracellular *C. neoformans* infection when compared to J774 macrophage-like cells. Specifically, we show that FLAMs, but not J774 cells, undergo a profound metabolic shift following *C. neoformans* infection, and switch from utilizing mitochondrial respiration to glycolysis as the primary mechanism of ATP production. This shift was accompanied by the heightened expression of glycolytic genes, including *Slc2a1, Pgk1, and Ldha*, which was found to be dependent on hypoxia inducible factor 1 (HIF1) activity.

## Results

### The transcriptome profile of FLAMs is highly similar to primary alveolar macrophages

As a first step towards determining whether FLAMs are phenotypically more like AMs than other cell models that are frequently used for *C. neoformans* host-pathogen interaction studies, RNA-sequencing (RNAseq) analysis was performed on resting FLAMs and J774.7 cells. These transcriptome profiles were compared to previously published datasets from RAW264.7 cells (9), primary AMs and peritoneal macrophages (PMs) from the Immunological Genome Project (ImmGen) database (30), as well as primary and *ex vivo* AMs from Subramanian *et al* 2022 (31). These data were clustered, and differences visualized as a multidimensional scale plot (MDS; Fig. 1A). Here, three distinct clusters were apparent when visualized using the first two dimensions, PC1, which represents 44.6% of the variability in the samples and PC2, which represents the next 33.2% of the variability in the samples. One cluster had all FLAM and AM samples, while the PM samples formed a second cluster. Finally, a third cluster contained RAW264.7 (RAWM0) and J774 (J77M0) samples. Within the first cluster, both sets of primary AM samples clustered together tightly with the FLAM samples clustering more closely with ‘AMX2’ samples, which represent alveolar macrophages cultured *ex vivo* for two months.

**Figure 1:**
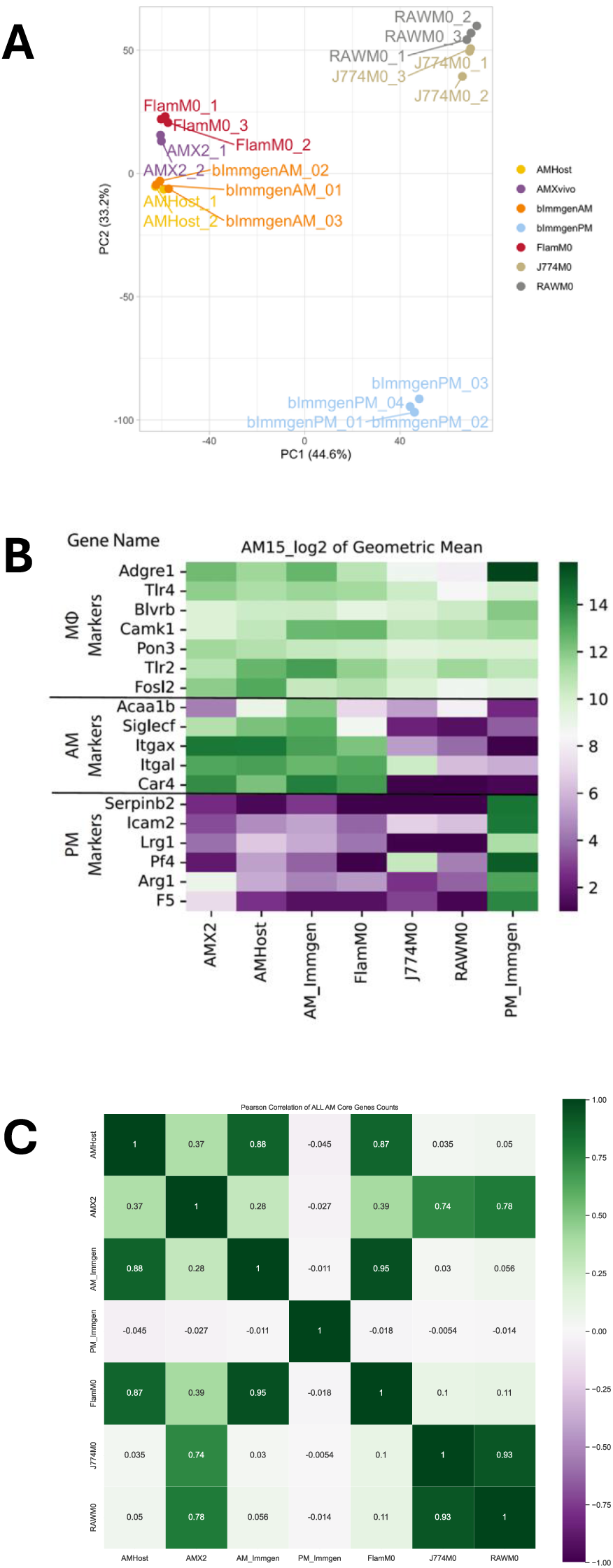
Steady-state transcript levels in FLAMs and AMs are highly similar. (A) Steady-state transcript levels in unstimulated FLAMs (‘FlamM0’) and J774 macrophage-like cells (J774M0’) were measured using RNAseq and compared to previously published transcriptome data from primary alveolar macrophages (AMs; ‘bImmgenAM’ (30) and ‘AMHost’ (31)), alveolar macrophages grown *ex vivo* for two months (‘AMX2’ (31)), peritoneal macrophages, (PMs; ‘bImmgenPM’ (30)), and RAW264.7 cells (‘RAWM0’ (9)) in a multidimensional scale (MDS) plot. Data from between two and four biological repeats were included for each cell type (indicated by the number following each cell type label on the plot). (B) The expression (log2 geometric mean) of common macrophage, AM, and PM markers in each transcriptome dataset are presented as a heat map. Markers were selected based on data from Gautier *et al* 2012, Thomas *et al* 2022, and Ankley *et al* 2024 (29,101,104). (C) The expression of 155 known AM markers in these transcriptome datasets was also compared using Pearson’s correlation (101). A value of 1 (green) indicates a perfect correlation, whereas negative values (purple) indicate anti-correlation.

As an alternative approach to appraising the similarity of these transcriptome datasets to that expected for AMs, the expression of known macrophage, AM, and PM markers was compared (Fig. 1B). As expected, expression of the general macrophage markers, which included *Adgre1*, *Tlr2*, and *Tlr4*, was observed in all seven transcriptome datasets, and the expression of the PM markers was largely restricted to the PM samples. While expression of the core AM markers was mostly low or absent in J774, RAW264.7, and PM samples, these were expressed at high levels in FLAMs except for *Acaa1b*, which was present at a moderate to low level. This core AM marker is involved in lipid metabolism and is known to diminish in expression when AMs are cultured *ex vivo* (31). Consistent with this notion, amongst all AM transcriptome data, *Acaa1b* transcript levels were lowest in the AMs cultured *ex vivo* for two months (AMX2). Finally, the expression of 155 AM markers was used to compare the transcriptome data sets by Pearson correlation (Fig. 1C). This analysis indicated that FLAMs were most similar to primary AMs in the expression of these signature AM genes than AMs cultured *ex vivo* and were highly dissimilar in the expression of the AM genes in PMs and the two macrophage cell lines tested (J774 and RAW264.7).

### Naïve FLAMs favor mitochondrial respiration

Possibly due to the high-oxygen and low-glucose environment of the lung, AMs favor mitochondrial respiration over glycolysis, with primary murine AMs generating ∼92% of all ATP by this mechanism (20). As our transcriptome analyses indicated that FLAMs were highly similar to primary AMs, we hypothesized that these cells would also phenocopy this aspect of AM biology and generate a larger proportion of their ATP by mitochondrial respiration than glycolysis, as compared to tumor-derived macrophages, such as J774 cells. To test this, the glycolytic and mitochondrial ATP production rates of FLAMs and J774 cells were compared. Consistent with our hypothesis, FLAMs were found to generate 85.74% of all ATP by mitochondrial respiration, similar to the levels reported for primary AMs (20), with only 14.26% produced by glycolysis (6:1 ratio; Fig. 2A). In contrast, J774 cells produced 60.19% of their ATP from mitochondrial respiration and 39.81% from glycolysis (1.5:1 ratio).

**Figure 2:**
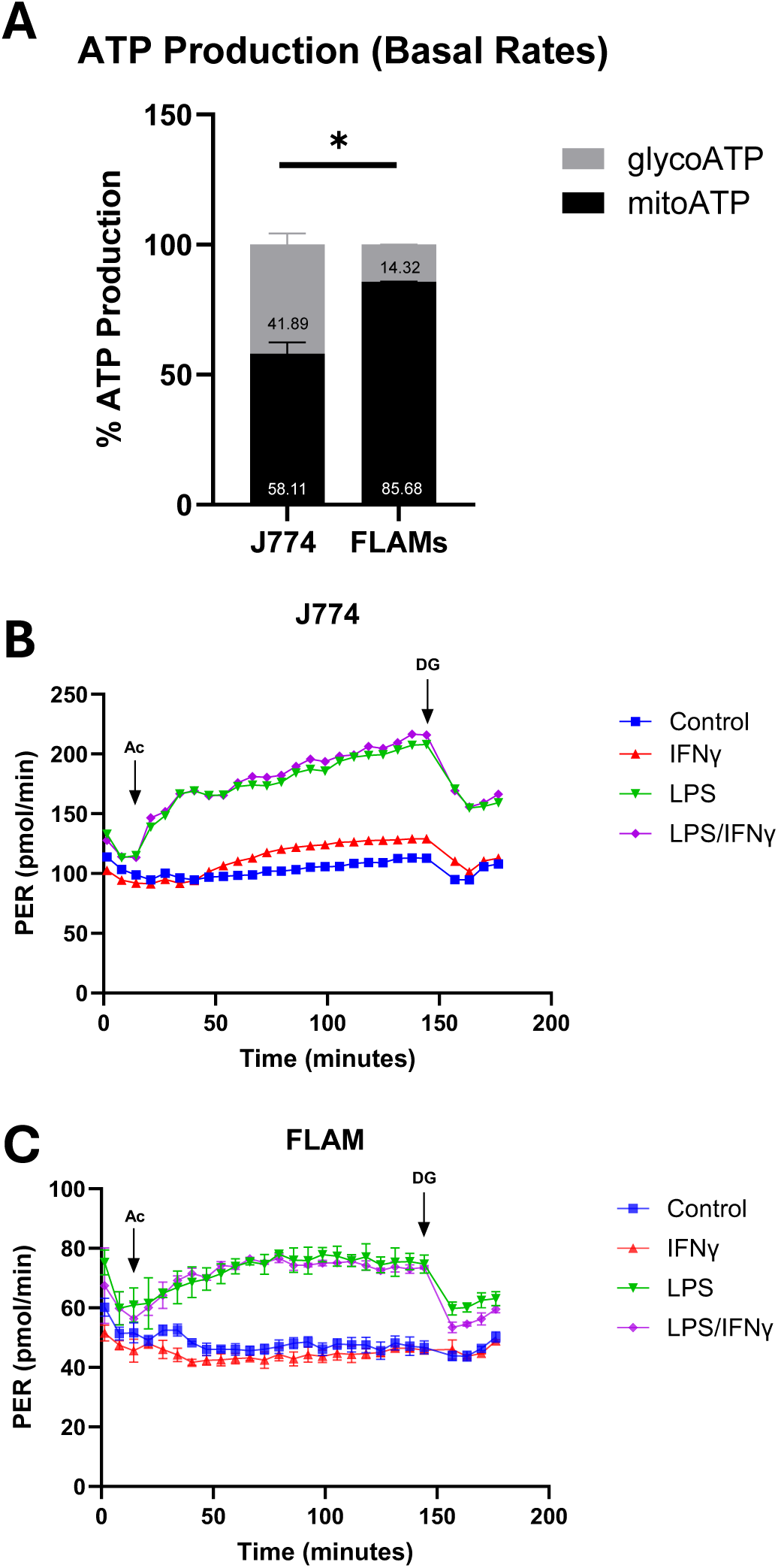
Unstimulated FLAMs use mitochondrial respiration as the primary mechanism of ATP production. (A) ATP production in unstimulated FLAM and J774 cells was measured using an XF Real-Time ATP Rate Assay (Agilent). (B+C) The change in proton efflux rate (PER) as a measure of altered glycolytic activity was measured in (B) J774 and (C) FLAM cells using a modified version of the Agilent Seahorse XF-Mini Macrophage Activation Assay. Arrows ‘Ac’ and ‘DG’ demarcate the timing of ‘activator’ and 2-deoxyglucose injection, respectively. Activators were IFNγ (200 units/mL) and LPS (100 ng/mL), delivered individually and in combination. All experiments were performed as a minimum of three biological repeats. Error is represented as S.E.M. Statistical differences between the samples in A were appraised using an unpaired *t*-test. Statistical significance is indicated as follows: * *p*<0.05.

A rapid increase in macrophage glycolytic activity is known to occur following exposure to select pro-inflammatory stimuli, including LPS, and is associated with polarization to the M1 state. To determine whether FLAMs perform a similar metabolic shift, changes in proton efflux rate (PER; a proxy for glycolytic activity) were measured in FLAM and J774 cultures following exposure to IFNγ and/or LPS. For both FLAMs and J774 cells, LPS treatment stimulated a rapid increase in PER, which was not enhanced by IFNγ co-treatment, and was sustained until glycolysis was inhibited by treatment with 2-deoxyglucose (Fig. 2B+C). Exposure to IFNγ alone had no significant impact on glycolysis over the period of observation in either cell type. Furthermore, incubation with IFNγ for 24 h prior to measurement had no significant impact on overall ATP production or the relative amounts produced from mitochondrial respiration and glycolysis (Fig. S1).

### FLAMs and J774 cells exhibit distinct responses to IFNγ stimulation

As activation of the IFNγ:JAK:STAT1 signaling axis is essential for a robust innate immune response to pulmonary *C. neoformans* infection in rodent models (32–34), we examined the gene expression responses of FLAMs to IFNγ by RNA sequencing and compared these data to equivalent J774 transcriptome data sets. In our hands, both cell types demonstrated a robust response to IFNγ with similar numbers of differentially expressed genes (DEGs; Fold Change (FC) ≥ 2.0 and q ≤ 0.05) identified in J774 (932 genes) and FLAMs (961 genes) (Fig. 3A-D). Direct pairwise comparisons of these transcriptome datasets, separating up– and down-regulated DEGs, revealed substantial differences in the response between these two macrophage types (Fig. 3E+F). Firstly, there was an almost even split in the number of up-(466; 48.5%) and down-regulated genes (495; 51.5%) in FLAMs, whereas a greater proportion of J744 genes were up-regulated (585; 62.8%) than down-regulated (347; 37.2%). Amongst up-regulated DEGs, 207 were common to FLAMs and J774 cells, representing less than half for each cell type (FLAMs=44.4%; J774=35.4%). These shared up-regulated DEGs included *Stat1* and *Irf1*, indicating that the STAT1-IRF1 reciprocal positive feedback loop that regulates the transcriptional response to IFNγ was activated in both cell types (35). This was accompanied by increases in steady state RNA levels of other *Irf* genes (*Irf2, Irf8*) and *Socs3*, a JAK/STAT signaling regulator (36), and a range of known interferon-inducible genes (ISGs). These included various members of the interferon-induced protein with tetracopeptide repeats (*Ifit*) family of genes (*Ifit1*, *2*, and *3*), Guanylate binding protein (*Gbp*) family of genes (*Gbp2*, *3*, *6*, *7*, and *11*), and *Isg15*. As a notable difference, the M1 marker *Nos2* was upregulated in J774 cells (7.33-fold; q=0.0006), but not in FLAMs, although a decrease in the M2 marker, *Arg1* was detected (4.66-fold, q=0.0006).

**Figure 3:**
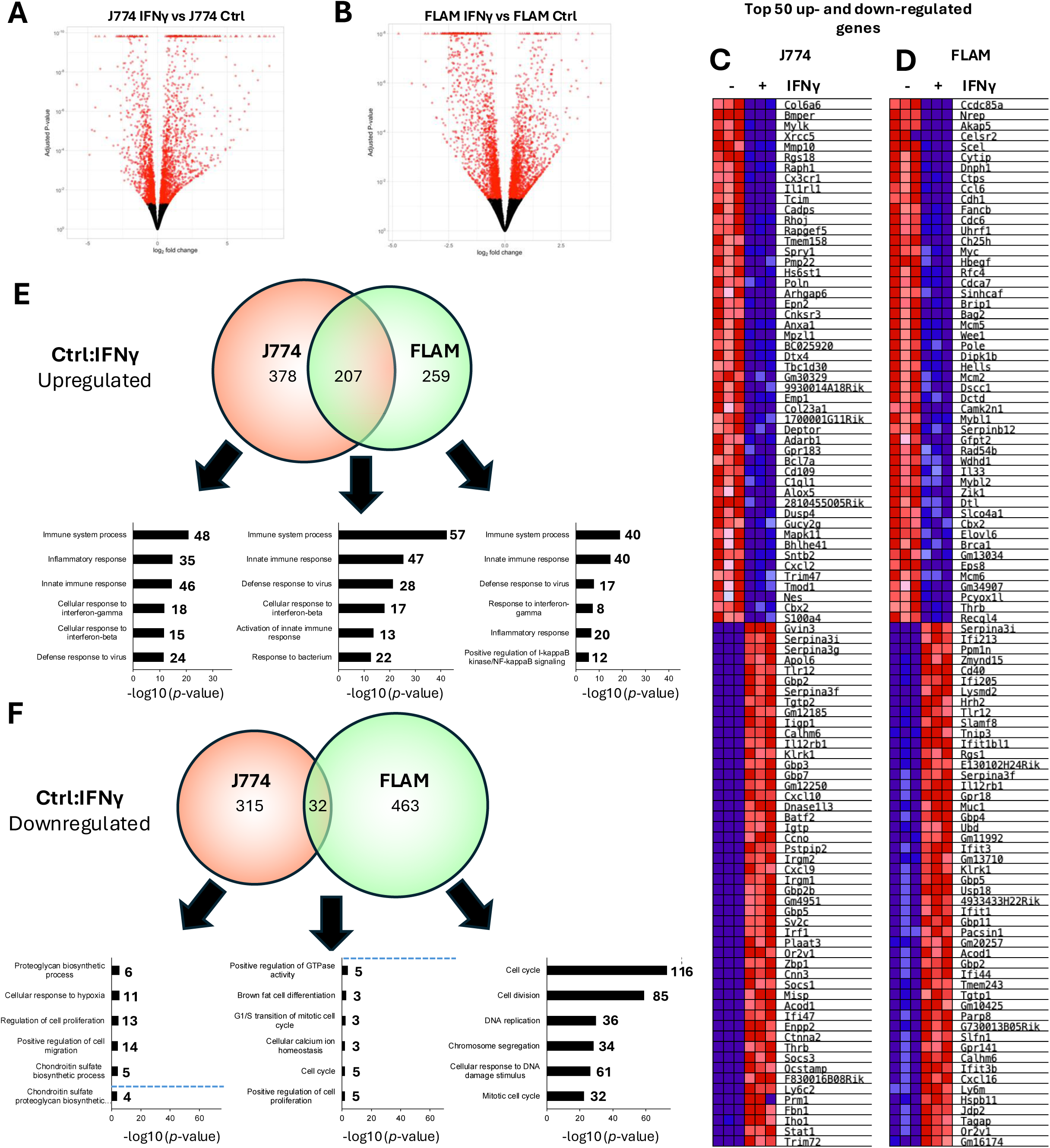
FLAM and J774 cells exhibit distinct responses to IFNγ stimulation. (A+B) J774 and FLAM cells were left untreated (Ctrl) or were incubated with IFNγ (200 units/mL) for 48 h and then harvested for RNA sequencing analysis. Differentially expressed genes (DEGs) were identified between Ctrl and IFNγ-stimulated (A) J774 and (B) FLAM cells. Genes with an adjusted *p-value* <0.05 are displayed in red. (C+D) The same transcriptome data set was subjected to GSEA analysis and the top 50 up– and down-regulated genes from (C) J774 and (D) FLAM cells between control and IFNγ-stimulated cells are displayed as heatmaps. (E+F) Common and differentially expressed genes in J774 Ctrl:IFNγ and FLAM Ctrl:IFNγ pairwise comparisons are represented as Venn diagrams for (E) upregulated and (F) downregulated genes. Gene ontology (GO) analysis was performed for each gene set in DAVID and the top seven biological process (BP) terms are displayed ranked by –log(*p*-value). The number of genes associated with each term is reported and all GO terms are significant (False discovery rate <0.5%) except those below the dashed blue line where this is present.

When the function of these regulated genes was examined using gene ontology (GO) tools, we found that biological process (BP) terms “immune system process” and “innate immune process” were associated with the common, J774 exclusive, and FLAM exclusive up-regulated DEG sets, with the highly similar terms “cellular response to interferon-gamma” or “response to interferon-gamma,” appearing for J774 and FLAM exclusive DEGs, respectively. While these terms are broad, these data indicate that common processes are affected in both FLAMs and J774 cells following IFNγ stimulation, although associated gene expression changes show some differences between the two cell types.

Amongst the down-regulated DEGs, only 32 genes were common between FLAMs (6.5%) and J774 cells (9.2%), with no GO terms reaching statistical significance for this set. For the J774 exclusive genes, top GO terms included diverse biological processes, such as “proteoglycan biosynthetic process,” “cellular response to hypoxia,” “regulation of cell proliferation,” and “positive regulation of cell migration.” However, down-regulated DEGs in FLAMs were highly enriched in genes associated with cell proliferation, with top GO terms including “cell cycle,” “cell division,” “DNA replication,” and “chromosome segregation”. Collectively, these data indicate that while the gene expression responses of J774 and FLAMs to IFNγ share some commonalities, particularly amongst up-regulated genes, they are largely distinct, with exposure to IFNγ resulting in decreased expression of genes associated with cell proliferation in FLAMs but not in J774 cells.

### C. neoformans replicates within M1-polarized FLAMs

Previous studies in mouse pulmonary infection models have shown that AMs will readily ingest *C. neoformans* following intratracheal infection (37). This can progress to a long-lasting intracellular infection where *C. neoformans* utilizes these cells as a niche for growth and replication (38). As mice deficient in AMs show decreased *C. neoformans* dissemination to other tissues, including the CNS (39), it has also been hypothesized that these cells function as a vehicle or ‘Trojan horse,’ contributing to this process (40,41). The spread of *C. neoformans* infection *in vivo* is likely aided by two behaviors observed in infected phagocytes, including AMs: vomocytosis, the non-lytic escape of *C. neoformans* from host cells without compromising their viability (42,43), and dragotcytosis, a macrophage-to-macrophage *C. neoformans* transference event (44,45). These behaviors and intracellular replication of *C. neoformans* have been observed in a variety of macrophage cell line models, including BMDMs (46), J774 (11), and RAW264.7 cells (10), but have yet to be explored in FLAMs, creating a barrier to their application as a model for AM:*C. neoformans* interactions. To address this, FLAMs and J774 cells were infected with EGFP-expressing *C. neoformans* and imaged by confocal microscopy in the presence of calcofluor white (CFW) and propidium iodide (PI) to label extracellular *C. neoformans* and dead cells, respectively. Here, 40.1% of FLAMs and 45.4% of J774 cells contained at least one live yeast cell, demonstrating these cells ingest *C. neoformans* with similar efficiency (Fig. 4A).

**Figure 4:**
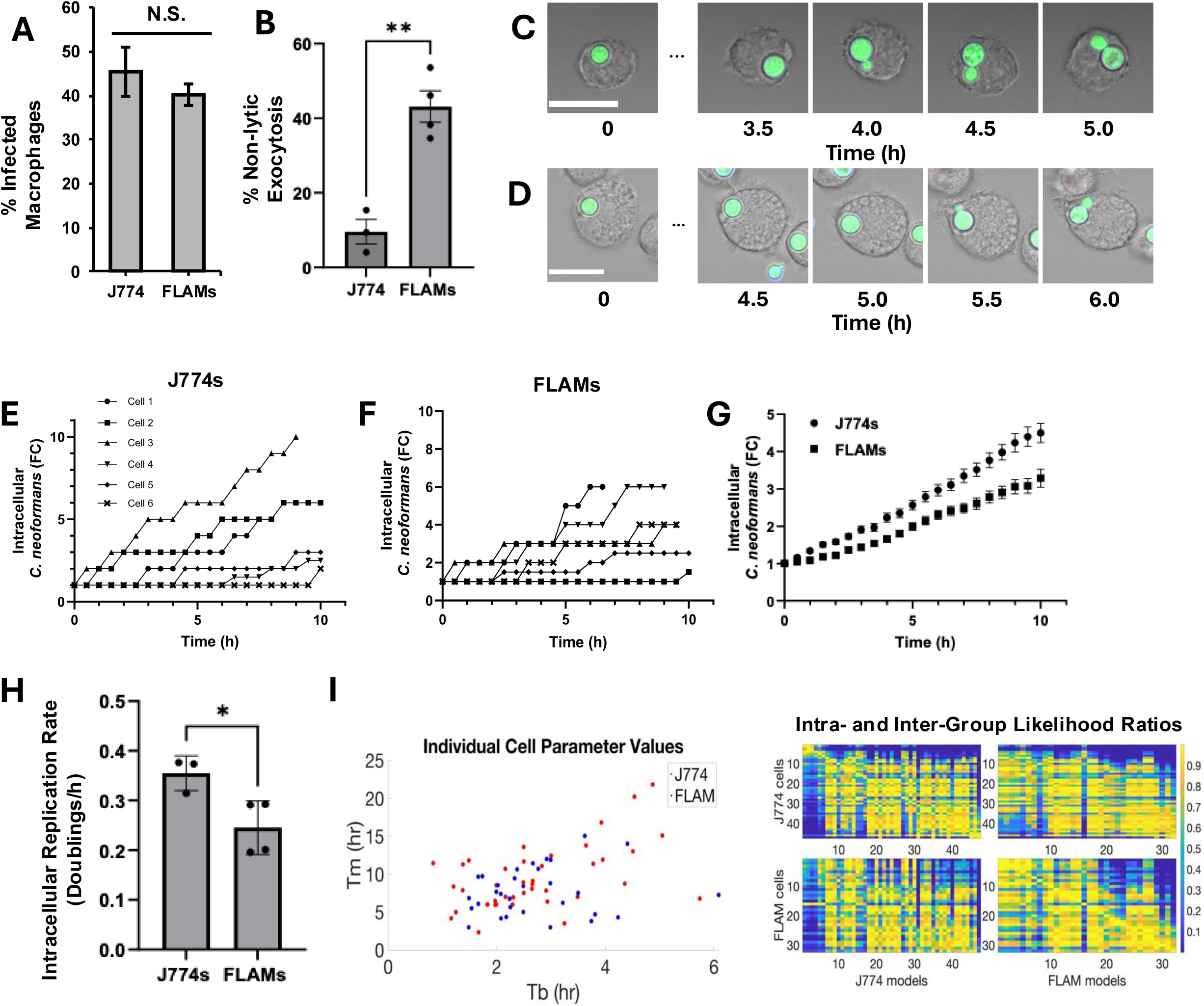
Intracellular replication of *C. neoformans* in J777 and FLAM cells. (A) J774 and FLAM cells were infected with opsonized H99S-EGFP *C. neoformans* at an MOI of 3:1 (*Cn*:Macrophage). Extracellular yeast was removed at 2 h post-infection, the cells were imaged by confocal microscopy, and the number of macrophages containing at least one intracellular yeast was quantified with >300 cells across three biological repeats counted for both J774 and FLAMs. (B-H) J774 and FLAM cells were infected at an MOI of 0.5:1 (*Cn*:Macrophage) with opsonized H99S-EGFP *C. neoformans*. Extracellular yeast was removed at 2 h post-infection, calcofluor white and PI were added to the cultures to label extracellular *C. neoformans* and dead cells, respectively, and the cells were subjected to time-lapse confocal microscopy. (B) The number of non-lytic exocytosis (NLE) events during the imaging period was counted for 114 and 84 infected FLAMs and J774s cells across four and three independent biological repeats, respectively, and expressed as the percentage. Representative images for *C. neoformans*-infected (C) J774 and (D) FLAMs are shown. Scale bar represents 20 μm. The number of intracellular *C. neoformans* in six representative macrophages at each time is expressed as the fold change (FC) and is shown for (E) J774 and (F) FLAM host cells. (G) The average number of intracellular *C. neoformans* per macrophage at each time is expressed as the fold change (FC) for J774 and FLAM host cells. (H) Intracellular replication rate is expressed as doublings per hour and was calculated by using the line of best fit for the FC in intracellular *C. neoformans* over time for each biological repeat. Here the first 8 hours of observation for 84 J774 host cells across 3 biological repeats and 114 FLAM host cells across 4 was used. For A, B, and H, error is represented as S.E.M., and significance was tested using an unpaired *t*-test with a Welch’s correction and is indicated as follows: N.S., not significant; * *p*<0.05; \*\**p*<0.01. (I) Left: Scatter plot of best-fit individual-cell parameter values. Goodness of fit was determined via maximum likelihood parameter estimation as described in the mathematical appendix. Right: Matrix of likelihood ratios for individual cells. Likelihood ratios serve as a relative measure of the similarity of the growth processes in individual cells by quantifying the ability of the best-fit model for one cell to describe the data of another relative to its best-fit model. Cells are arranged by the length of their count data in descending order.

Having established that FLAMs were able to ingest *C. neoformans*, we compared the cellular behavior and intracellular growth rate of the pathogen in FLAMs and J774 cells using live cell microscopy. In brief, macrophages were infected with fluorescent *C. neoformans* at a low MOI (0.5:1) to increase the spatial separation of infected cells and reduce the initial fungal burden of individual macrophages. The infected cells were then imaged every 30 minutes for up to 24 h. We observed non-lytic exocytosis (NLE), as indicated by the non-lytic release of *C. neoformans*, in both FLAM and J774 cells. However, the rate of NLE was higher in FLAMs than J774 cells (*p*<0.01), occurring in 43.1% of FLAMs versus 9.59% J774 host cells over the first 10 h of the observation period (Fig. 4B). Our measurement of *C. neoformans* NLE in J774 cells was similar to that reported in an earlier study, where this was observed in 9.7% of infected cells over the same period (43). At the level of individual host cells, intracellular *C. neoformans* replication rate was highly variable in both FLAM and J774 cells (Fig. 4C-F). However, when viewed at the population level, the intracellular replication rate appeared slower in host FLAMs with an average of 0.25 doublings occurring per hour versus and 0.35 in J774 cells (*p*<0.05; Fig. 4G+H).

Given the high potential for individual-cell variability and apparent differences in the rate of censoring events (e.g., vomocytosis), to more rigorously appraise *C. neoformans* intracellular replication in FLAMs and J774s, we constructed a mathematical model of *C. neoformans* population growth via asymmetric budding. In the model, a population’s growth potential is determined by two parameters, the inter-budding time, or time between the successive emergence of buds in a mature mother cell, and the maturation time, or time between a daughter cell’s emergence as a bud and its first budding event. The population growth model was coupled to a probabilistic model of counting errors which result from small cells being caught between z-slices, mother cells obscuring daughter cells, miscounts, and artifacts. This enabled us to determine best-fit growth parameters via maximum likelihood parameter estimation (See Mathematical Modeling Appendix and Materials and Methods for details related to mathematical modeling, parameter estimation, and numerical implementation).

The intracellular population growth model was fit to individual cell data from each host. Since this method fits the overall growth pattern for each cell, as opposed to the population size at each time, the time interval for the model-based analysis was extended until the end of the observation period for each cell. However, cells with limited data, i.e., those that experienced very few complete budding events (n<2) were excluded since there is limited information to parameterize the budding and maturation process in these cells. After data processing, 32 FLAMs and 47 J774s met our criteria for individual-cell parameter estimation. The ability of the model to describe the intracellular count data was confirmed visually and by the low value of the normalized negative log likelihoods of the fits (Mathematical Appendix Fig. 12), where normalization was with respect to the number of observations in an individual cell count. Best-fit parameter distributions for J774s and FLAMs were visually similar. However, the J774 parameter distribution was more variable and included a few very slow-growing outlier cells (Fig. 4I). The similarity of *C. neoformans* proliferation in the two host cells was further quantified through inter– and intra-group likelihood ratios for the best-fit models to individual cells. The mean likelihood ratio of J774 models for FLAM data was 0.5749. The mean likelihood ratio of FLAM models for J774 data was 0.5952. The mean inter-group likelihood ratio for J774 cells (0.5952) was actually higher than the mean intra-group likelihood ratio for this host cell type (0.5541). While this can be explained in part by the presence of outlier cells in the J774 data, the high values of the mean inter-group likelihood ratios do not support the idea that rates of intracellular proliferation differ between the two host cell types. Since likelihood ratios represent probabilities, we also considered using the geometric mean of the inter– and intra-group likelihood ratios to quantify inter– and intra-group variability. However, due to the presence of outlier cells in the J774 population, the geometric mean of the inter– and intra-group likelihood ratios for J774s was zero.

Next, we computed population-level best-fit parameter estimates for each host cell type by pooling data for each host type and weighting individual observations (as opposed to individual cell counts) equally. The descriptive ability of population-level parameters for individual cells can be seen in Mathematical Appendix Fig. 10. Population-level best-fit parameters also indicate similar intracellular proliferation dynamics in the two host cell types. Best-fit inter-budding times were about 158 minutes in J774 cells, compared to 148 minutes in FLAMs. Best-fit maturation times were very long in both host cells, over 20 hours in J774 cells, and over 15 hours in FLAMs. While long maturation times were observed in the microscopy data, these very long times may reflect the challenge of fitting this parameter to an entire population. For example, in individual J774 cells, the largest best-fit maturation time was almost ten times longer than the smallest. As additional points of comparison, we compared the model-predicted, population stable stage distributions and doubling times for the two host cell types. At the stable stage– and age-distribution, about 78% of FLAMS and 81% of J774 cells are immature daughter cells. Meanwhile, the population doubling time at the stable stage– and age-distribution is about 7 hours in FLAMs and 8.6 hours in J774 cells. We note these doubling times are much longer than those corresponding to the doubling rates found previously, namely about 4 hours and 2.8 hours in FLAMs and J774 cells, respectively. However, this difference is not necessarily contradictory since the population doubling time is time-dependent for an age-and stage-structured population. For example, the doubling times determined previously may be heavily weighted toward the inter-budding time due to the initial state of the population, whereas the doubling time at the stable age– and stage-distribution provides a time-invariant measure of the population’s growth potential. In summary, our model-based analysis returns slightly slower rates of intracellular proliferation in J774 cells compared to FLAMs. However, large inter-group likelihood ratios and very similar inter-budding times suggest that *C. neoformans* intracellular proliferation proceeds at similar rates in these host cell types.

We next conducted a model-based analysis of NLE and other cell fates in J774 and FLAM cells. Here NLE and other fates are events that trigger then end of the observation for an individual macrophage. Examples of other cell fates include cell death, cell division, detachment, and migration out of the frame of view. It should be noted that since NLE and other cell-fate events trigger the end of an observation, these events censor each other’s event times. For the analysis, NLE and other cell-fate events were assumed to be controlled by a sequence of competing Poisson processes. This assumption provides a model template so that the number of stages required to reach an NLE fate and the rates of progression and exit from cell stages can be fit via maximum likelihood parameter estimation. The most appropriate cell-fate model for each host cell type was then selected via the Akaike information criterion. See Materials and Methods for details.

In FLAMS, 107 of 217 (49.3%) cells experienced NLE events before the end of the experiment, while only 3 FLAM cells (1.4%) failed to experience a cell-fate event before the end of the experiment. The selected cell-fate model for FLAMs included just one stage (two parameters) and predicts about 50.2% of cells will experience an NLE event (before a censoring event occurs). Neglecting the rate of transition into other events in the selected cell-fate model for FLAMs, we find the mean time to experience an NLE event is about 2.6 hours in this host cell type.

In J774 cells, 8 of 93 (8.6%) cells experienced NLE events before the end of the experiment, while 5 J774 cells (5.4%) failed to experience a cell-fate event before the end of the experiment. The selected cell-fate model for J774 cells included two stages (four parameters) and predicts about 8.6% of cells will experience an NLE event (before a censoring event occurs). Neglecting the rate of transition into other events in the selected cell-fate model for J774 cells, we find the mean time to experience an NLE event is about 10.8 hours in this host cell type.

To quantify the significance of differences in the dynamics of NLE events between host cell types, we tested the ability of the NLE model framework selected for one host type to describe the cell-fate data from the other host cell type by holding rates governing NLE events fixed and optimizing over rates of exit to other events. Here too the Akaike information criterion is used to compare models, penalizing each model by the number of free parameters which were fit to the data. When assessing the descriptive ability of the FLAM NLE model for the J774 data, the best fit of the FLAM NLE model to the J774 data is penalized for a single free parameter while the selected model for describing the J774 data is penalized for each of its four parameters. Similarly, when assessing the descriptive ability of the J774 NLE model for the FLAM data, the best fit of the J774 NLE model to the FLAM data is penalized for two free parameters while the selected model for describing the FLAM data is penalized for each of its two parameters. The relative likelihoods of the FLAM NLE model for the J774 data and the J774 model for the FLAM data were both less than 0.01. In summary, model-based analysis suggests FLAMs experience NLE events at about 4 times the rate of J774s and that this difference is significant.

### FLAMs and J774 cells exhibit distinct responses to C. neoformans infection

Having established that FLAMs and J774s could ingest *C. neoformans* with similar efficiency and that the pathogen could utilize both cell types as a growth niche, we compared the gene expression responses of these cells to infection with the pathogen. This was achieved by infecting macrophages with *C. neoformans* and carefully replacing the culture medium at regular intervals to remove un-ingested and extruded yeast, preventing excessive nutrient depletion by extracellular *C. neoformans*, as has previously been reported (9). The macrophages were harvested, and differences in gene expression between mock and *C. neoformans*-infected FLAMs and J774 cells were appraised by RNAseq.

Our transcriptome profiling data show that FLAMs undergo a far more extensive and dramatic response to *C. neoformans* infection than J774 cells (Fig. 5A-F), as indicated by both the number of DEGs associated with IFNγ:IFNγCn pairwise comparison, which was 1393 DEGs in FLAMs versus 685 in J774 cells, and the broader distribution in the fold changes of these DEGs observed in FLAMs (Fig. 5A+B). Additionally, the number of upregulated genes associated with the FLAM IFNγ:IFNγCn pairwise comparison (819 DEGs) was greater than in J774 cells (302 DEGs), with only a modest overlap of 116 genes observed between the two, which represented only 14.2% of upregulated DEGs in FLAMs and 38.4% in J774 cells (Fig. 5E). As might be expected, these genes were associated with the GO BP terms “Inflammatory response,” and “Cellular response to interferon-gamma,” but ten of these common genes were also associated with the term “cellular response to hypoxia.” A similar term, “Response to hypoxia,” appeared together with “canonical glycolysis” among the top GO terms associated with FLAM-exclusive DEGs. This was notable as the HIF1 signaling pathway, which regulates the transcriptional response to hypoxia, also promotes an increase in macrophage glycolytic activity following exposure to various pathogens or pathogen-associated molecular patterns (PAMPs), such as LPS (47).

**Figure 5:**
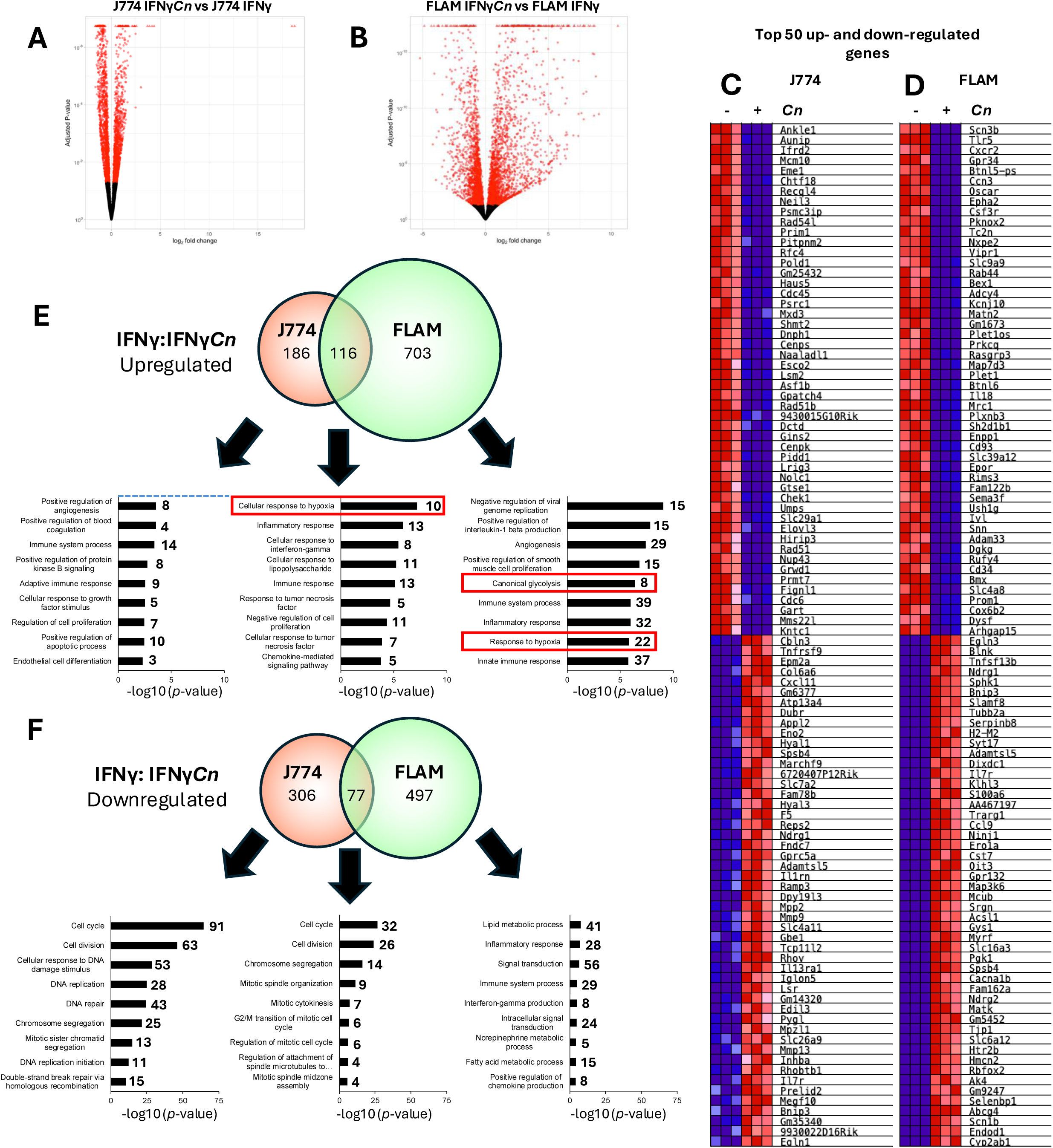
FLAM and J774 cells exhibit distinct responses to C. neoformans infection. (A+B) J774 and FLAM cells were incubated with IFNγ (200 units/mL) for 24 h, mock-infected (IFNγ) or infected with opsonized H99S *C. neoformans* (IFNγ*Cn*) and then harvested for RNA sequencing analysis. Differentially expressed genes (DEGs) were identified between mock and *C. neoformans*-infected (A) J774 or (B) FLAMs. Genes with an adjusted *p-value* <0.05 are displayed in red. (C+D) The same transcriptome data set was subjected to GSEA analysis and the top 50 up– and down-regulated genes from (C) J774 and (D) FLAM cells between mock and *C. neoformans*-infected cells are displayed as heatmaps. (E+F) Common and differentially expressed genes in J774 IFNγ:IFNγ*Cn* and FLAM IFNγ:IFNγ*Cn* pairwise comparisons are represented as Venn diagrams for (E) upregulated and (F) downregulated genes. Gene ontology (GO) analysis was performed for each gene set in DAVID and the top seven biological process (BP) terms are displayed ranked by –log(*p*-value). The number of genes associated with each term is reported, and all GO terms are significant (False discovery rate <0.5%) except those below the dashed blue line where this is present. GO terms of special interest are marked in red.

The overlap in downregulated genes was also modest (Fig. 5F), with only 77 genes common between FLAMs (13.4% of 574 DEGs) and J774 cells (20.1% of 383 DEGs). While GO terms for J774-exclusive DEGs were largely associated with cell proliferation and DNA integrity, and included “cell cycle,” “cell division,” and “cellular response to DNA damage stimulus,” the top GO term for FLAM-exclusive downregulated DEGs, “Lipid metabolic process,” was also related to metabolism. Collectively, these data indicated that *C. neoformans* infection may alter FLAM metabolic activity.

To better understand how *C. neoformans* infection altered the transcriptome of IFNγ-stimulated FLAMs, we also performed a pairwise comparison of Ctrl:IFNγ and Ctrl:IFNγ*Cn* datasets. In both FLAM and J774 cells, the number of DEGs for Ctrl:IFNγ*Cn* was far larger than Ctrl:IFNγ (Fig. S2A+B), and the degree of overlap was high with 70.7% and 75.2% of Ctrl:IFNγ DEGs also appearing in Ctrl:IFNγ*Cn* for FLAM and J774 cells, respectively. This indicated that *C. neoformans* infection resulted in broader transcriptional changes in both cell types than exposure to IFNγ alone. Notably, the GO terms “Lipid metabolic process,” “Response to hypoxia,*”* and “Canonical glycolysis” were associated with DEGs exclusive to Ctrl:IFNγ*Cn* in FLAMs but not J774 cells, suggesting that metabolic reprogramming was a major aspect of the broader gene expression changes occurring following *C. neoformans* infection in this alveolar macrophage model.

To investigate this more thoroughly, the same pairwise comparisons were performed for upregulated and downregulated genes. Here, we observed a high degree of overlap with 336 of the 466 upregulated DEGs (72.1%) appearing in the Ctrl:IFNγ pairwise comparison also present in Ctrl:IFNγ*Cn* (Fig. S3A). GO terms associated with this common set of DEGs were broad and included “innate immune system process” and “activation of innate immune response.” However, 714 DEGs were exclusive to Ctrl:IFNγ*Cn*, representing 68% of all genes associated with the Ctrl:IFNγ*Cn*. A similar pattern was observed for downregulated genes, with the majority of Ctrl:IFNγ DEGs (319 of the 495 DEGs; 64.4%) also appearing in the Ctrl:IFNγ*Cn* dataset (Fig. S3B), and 535 of the 854 DEGs in Ctrl:IFNγ*Cn*. These common DEGs were associated GO terms concerning cell replication and the DNA damage response, including “cell cycle,” “chromosome segregation,” and “DNA repair.” Interestingly, the metabolism and hypoxia-associated GO terms observed in our analysis of the complete Ctrl:IFNγ*Cn* DEG set were split between up– and downregulated genes with “response to hypoxia,” “cellular response to hypoxia,” “glycolytic process,” and “canonical glycolysis” appearing significant for Ctrl:IFNγ*Cn* exclusive up-regulated genes and “lipid metabolic process” and “fatty acid metabolic process” significant for the corresponding set of down-regulated genes. As lipid metabolism is typically associated with mitochondrial respiration, these data were consistent with a transition from mitochondrial activity towards glycolysis in FLAMs.

Analysis of the corresponding J774 data set revealed a higher degree of overlap between upregulated Ctrl:IFNγ and Ctrl:IFNγ*Cn* DEGs with 549 of 585 (93.9%) from the Ctrl:IFNγ pairwise comparison also appearing in the Ctrl:IFNγ*Cn* dataset (Fig. S4A). Overlap between the downregulated genes was more modest, with only 148 of 347 DEGs (42.7%) from the Ctrl:IFNγ pairwise comparison appearing in the Ctrl:IFNγ*Cn* dataset (Fig. S4B). Consistent with our prior analyses, no GO terms associated with hypoxia or metabolism were significant for either up-or downregulated DEGs exclusive to the Ctrl:IFNγ*Cn* pairwise comparison. Additionally, while “immune system process*”* was the only significant GO term for upregulated DEGs exclusive to the Ctrl:IFNγ*Cn* pairwise comparison, GO terms from for corresponding downregulated DEGs included *“*cell cycle,” “chromosome segregation,” “DNA repair,” and “cellular response to DNA damage.”

To further compare the response of FLAM and J774 cells to IFNγ stimulation and *C. neoformans* infection, the expression of known polarization markers, cytokines, stress response genes, and regulators of proinflammatory gene expression were interrogated by quantitative reverse transcription-PCR (qRT-PCR). In FLAMs, IFNγ stimulation alone did not induce a change in *Nos2* RNA levels (Fig. 6A); *Nos2* encodes the murine M1-marker, inducible nitric oxide synthase (iNos). However, *C. neoformans* infection of these cells stimulated an ∼8,600-fold increase in *Nos2* RNA levels. Interestingly, this was accompanied by increased steady state RNA levels of the M2 marker *Arg1* (Supplementary data file 1). These data were consistent with known alveolar macrophage behaviors, as these cells are characterized as hyporesponsive to proinflammatory stimuli, including IFNγ (27), do not adhere to the M1-M2 paradigm, and exhibit mixed M1/M2 phenotypes (24,25). By contrast, IFNγ stimulation alone elicited a modest but statistically significant increase in *Nos2* transcript levels (8.9-fold) in J774 cells, which increased to 170-fold following *C. neoformans* infection (Fig. 6B), and this was not accompanied by *Arg1* expression.

**Figure 6:**
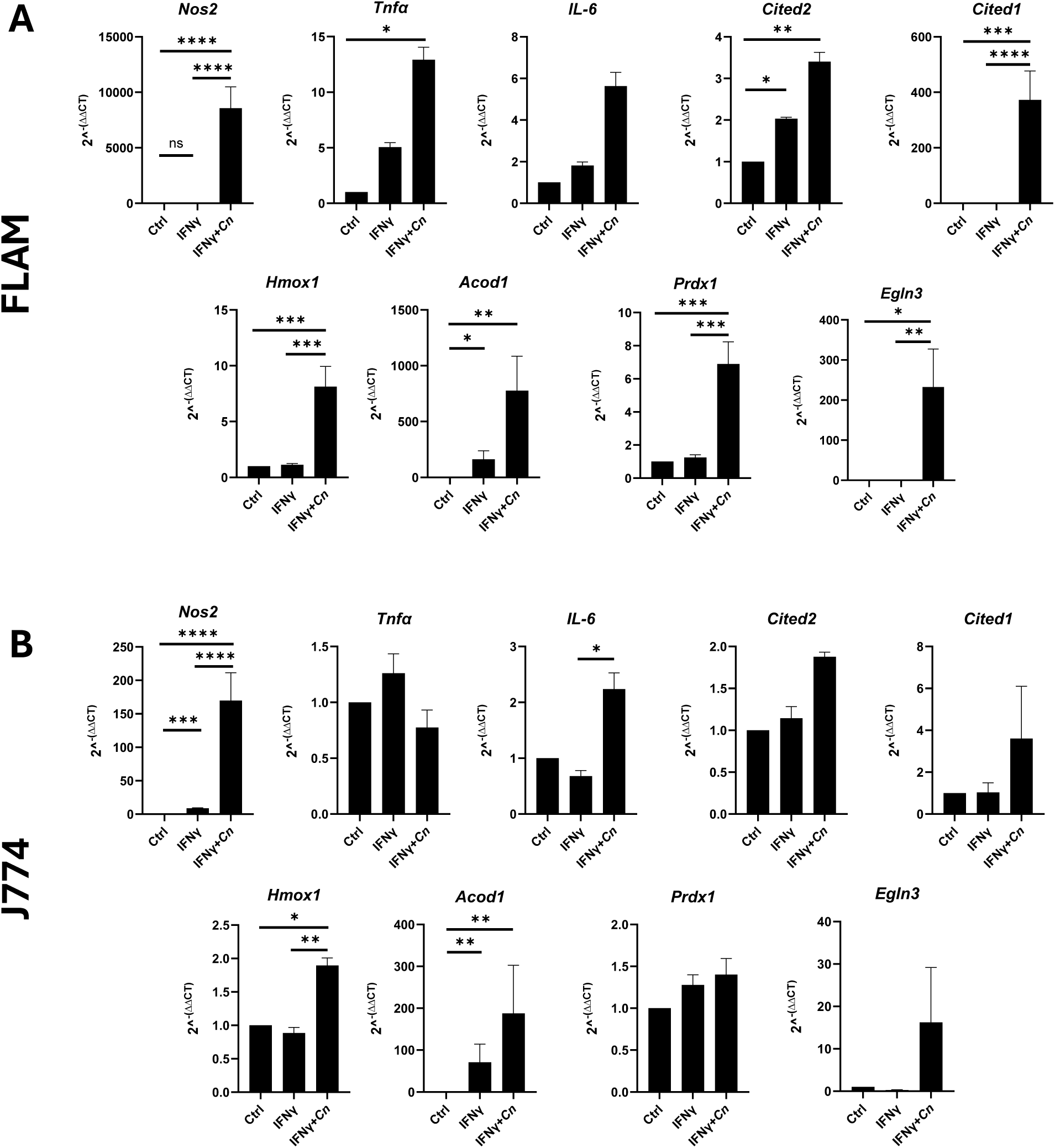
C. neoformans infection increases the expression of M1 markers and stress-responsive genes in FLAMs and J774 cells. (A) FLAM and (B) J774 cells were left untreated (Ctrl) or incubated with IFNγ (200 units/mL) for 24 h and then mock-infected (IFNγ) or infected with opsonized H99S *C. neoformans* (IFNγ*Cn*). RNA was harvested at 24 h post-infection, and the expression of the indicated transcripts was measured by qRT-PCR. Results are expressed as the mean of at least three biological repeats. Error is represented as the S.E.M. Statistical significance was assessed using a one-way ANOVA followed by a Tukey’s multiple comparison test and is indicated as follows: * *p*<0.05; \*\**p*<0.01; \*\*\**p*<0.001; ****p<0.0001.

The impact of IFNγ and *C. neoformans* infection on the steady state RNA level of genes encoding proinflammatory cytokines also differed between the two macrophage types. In our hands, neither IFNγ nor *C. neoformans* infection affected *Tnfa* levels in J774 cells, and *C. neoformans* infection resulted in only a very modest ∼2-fold increase in *IL-6* transcript levels over IFNγ stimulation alone (Fig. 6B). Whereas infection of FLAMs stimulated a robust 12.9-fold increase in *Tnfa* levels, and while *IL-6* levels were also increased 5.6-fold over unstimulated cells, this was not statistically significant (Fig. 6A).

As *C. neoformans* infection is known to induce host cell stress (8), we also measured the transcript levels of several DEGs identified in our transcriptome screen that are known to participate in stress response. Here, we found that *Hmox1* and *Acod1* transcript levels were increased in both FLAMs and J774 cells following infection with *C. neoformans*, although the fold change was far higher in FLAMs. *Hmox-1*, which encodes the heme oxygenase 1 (HO-1) enzyme, is primarily involved in the degradation of heme and the protection against oxidative stress, but may also decrease macrophage proinflammatory activity by promoting M2 repolarization (48). For *Acod1*, IFNγ stimulation alone was sufficient to promote its expression in both FLAMs and J774 cells, but this level was further increased following *C. neoformans* infection. This gene, which encodes aconitate decarboxylase 1, is known to be upregulated in macrophages following microbial infections and produces the metabolite itaconate, which may have immunomodulatory and antibacterial activities, although it is unclear whether it has antifungal properties (49–51). Additionally, from this set of stress-responsive genes, *C. neoformans* infection also stimulated increased levels of *Prdx1* transcripts, but only in FLAMs. The peroxiredoxin 1 protein encoded by this gene is involved in redox signaling and is known to enhance the expression of proinflammatory cytokines and stimulate glycolytic activity in macrophages following exposure to PAMPs (52).

The CBP/p300-interacting transactivator with the glutamic acid/aspartic acid-rich carboxy-terminal domain (CITED) family of transcriptional co-regulators has been shown to modulate the gene expression response to proinflammatory stimuli in macrophages. More specifically, CITED2 operates as a common co-repressor of STAT, IRF, NF-κB, and HIF1 by occupying the same binding site on CBP/p300 utilized by these transcription factors to recruit this histone acetyltransferase to gene cis-regulatory sites (53,54). In this way, CITED2 attenuates macrophage proinflammatory gene expression induced by IFNγ and various PAMPs (55–58). In contrast, CITED1 appears to function as a co-enhancer and has been shown to increase the RNA level of select interferon-stimulated genes (ISGs) in IFNγ-stimulated macrophages (59). Additionally, expression of both *Cited1* and *Cited2* has been shown to be affected by a range of pro-inflammatory stimuli, with *Cited1* transcript levels increased by either IFNγ stimuli or *C. neoformans* infection of RAW264.7 cells (9,59). Interestingly, RNA levels of both *Cited1* and *2* in J774 cells was unaffected by either IFNγ stimuli or *C. neoformans* infection. Indeed, *Cited1* transcript levels were near-undetectable in these cells by qRT-PCR. Whereas in FLAMs, *Cited2* RNA level was increased by either IFNγ stimulation or *C. neoformans* infection. While IFNγ stimulation had no effect on *Cited1* transcript levels in FLAMs, they were increased by >300-fold following *C. neoformans* infection.

Finally, as the transcriptome analysis indicated that *C. neoformans* infection resulted in increased levels of transcripts associated with the HIF1 pathway in FLAMs, expression of *Egln3*, a bona fide HIF1 target gene, was quantified in both macrophage types. While IFNγ-stimulation or *C. neoformans* infection did not have a statistically significant effect on *Egln3* transcript levels in J774 cells, these were increased >200-fold in infected FLAMs compared to unstimulated controls. These data were consistent with the notion that *C. neoformans* infection promotes HIF1 activity in FLAMs, but not J774 cells.

### C. neoformans infection stimulates a glycolytic shift in FLAMs

Our initial GO analysis of the transcriptome data indicated that *C. neoformans* infection stimulated an increase in RNA levels for glycolytic genes in FLAMs but not J774 cells. While macrophages exposed to an M1-polarizing environment are known to undergo a glycolytic shift, analogous to the Warburg effect, this change in FLAM glycolytic gene expression was unexpected as AMs are heavily dependent on mitochondrial respiration for ATP generation. To ensure that this result was valid, it was rigorously tested using a Pearson’s correlation to compare ‘global’ glycolytic gene expression in FLAMs, AMs, and macrophage cell lines. Additionally, the expression of individual glycolytic genes was also verified by qRT-PCR.

The Pearson’s correlation which utilized a 101-gene set representing all glycolytic genes associated with the GO terms “glycolytic process,” “canonical glycolysis,” and “regulation of glycolytic process” indicated that glycolytic gene expression in unstimulated FLAMs was more similar to primary AMs and AMs grown *ex vivo* for two months than J774 or RAW264.7 macrophage cell lines (Fig. 7A). While glycolytic gene expression in IFNγ-stimulated FLAMs (FlamM1) exhibited a slightly higher correlation with all AM transcriptome datasets (AMHost, AMX2, and AM_Immgen) than unstimulated FLAMs (Flam M0), the opposite was true for *C. neoformans*-infected FLAMs (FlamM1Cn). Here, the correlation value indicated that glycolytic gene expression in infected FLAMs (FlamM1Cn) was more similar to that in J774 (J774M0) and RAW264.7 (RAWM0) cells. Additionally, neither IFNγ stimulation (J774M1, RAWM1) nor *C. neoformans* infection (J774M1Cn, RAWM1Cn) resulted in large changes in the correlation values for J774 and RAW264.7 samples with other transcriptome datasets.

**Figure 7:**
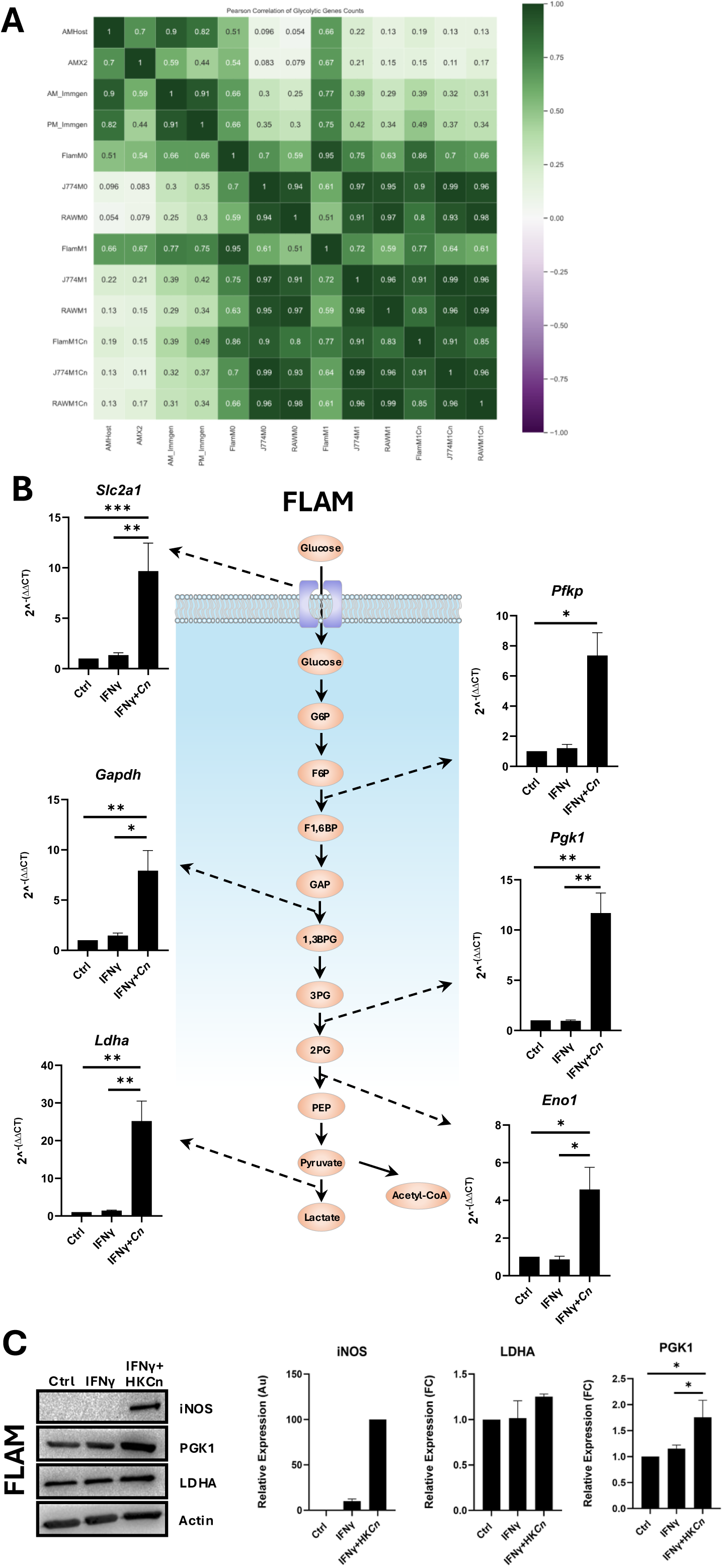
C. neoformans infection increases the expression of glycolytic genes in FLAMs. (A) The expression of 101 known glycolytic genes in the transcriptome datasets described in 1A, together with transcriptome datasets from IFNγ-stimulated (M1 suffix) and IFNγ-stimulated +*C. neoformans* infected (M1Cn suffix) FLAM, J774, and RAW264.7 macrophages, were compared using Pearson’s correlation (101). A value of 1 (green) indicates a perfect correlation, whereas negative values (purple) indicate anti-correlation. (B) qRT-PCR validation of select DEGs identified in the RNAseq screen described in Figure 5 that are associated with the *Canonical Glycolysis* GO term or play a role in the regulation of glycolysis. Results are expressed as the mean of at least three biological repeats. Error is represented as the S.E.M. Statistical significance is indicated as follows: \**p*<0.05; \*\**p*<0.01; ****p<0.0001 (one-way ANOVA followed by a Tukey’s multiple comparison test). The diagram provides a simplified representation of the glycolysis pathway with arrows representing enzyme-catalyzed reactions. Dashed lines link these with qRT-PCR data for the transcript encoding that enzyme or transporter. (C) FLAMs were stimulated with IFNγ (200 units/mL) for 24 h and mock-infected (IFNγ) or infected with heat-killed, opsonized H99S (IFNγ*Cn*) for 2 h before the remaining extracellular yeast was removed. Cells were harvested at 24 h post-infection, and iNOS, PDK1, and LDHA protein levels were measured by western blotting. Actin was utilized as a loading control. Protein levels were quantified by densitometry based on three discrete biological repeats.

To explore the effect of *C. neoformans* infection on the FLAM glycolytic pathway at the single-gene level, RNA levels of five of the eight DEGs associated with the ‘canonical analysis’ BP GO term that were shown to be upregulated in our transcriptome analysis, together with *Slc2a1*, which encodes the GLUT1 glucose transporter, were measured in untreated, IFNγ-primed, and *C. neoformans*-infected FLAMs by qRT-PCR. While IFNγ stimulation alone had no effect on any of the transcripts tested, *Slc2a1*, *Pfkp*, *Gapdh*, *Pgk1*, *Eno1*, and *Ldha* RNAs were all increased in FLAMs between 4– and 25-fold over untreated controls following *C. neoformans* infection (Fig. 7B). Consistent with the transcriptome analysis and Pearson’s correlation, *C. neoformans* infection had only minimal effects on the transcript levels of these genes in J774 cells with only *Slc2a1* and *Gapdh* showing increased transcript levels over IFNγ-stimulated, mock-infected samples (Fig. S5).

Before testing whether these changes in glycolytic gene transcript levels also resulted in increased glycolytic activity in infected FLAMs, we first tested whether heat-killed *C. neoformans* were also capable of stimulating glycolytic gene expression in these cells. Here, heat-killed *C. neoformans* was almost equally effective at stimulating increased *Slc2a1*, *Gapdh*, and *Ldha* expression (Fig. S6A). Infection with heat-killed *C. neoformans* also had similar effects to live yeast on the expression of the pro-inflammatory genes, including *Nos2*, *Tnfa*, and *IL-6* (Fig. S6A). As observed for infection with live *C. neoformans*, only *Nos2* expression was increased in J774 cells following infection with heat-killed *C. neoformans*, and the expression of all four glycolytic genes tested was not affected (Fig. S4B). Collectively, these data confirm that the effects of *C. neoformans* on glycolytic gene expression in the FLAM model of AM:fungal interactions and additionally indicate this response does not require live, metabolically active yeast. This finding also provided a methodology by which the impact of *C. neoformans* on host cell metabolic activity could be tested without concern that the metabolic activity of the yeast would interfere with measurements.

To determine whether the changes in glycolytic gene transcripts detected in FLAMs resulted in a corresponding increase in glycolytic enzymes, lactate dehydrogenase A (LDHA) proteins, phosphoglycerate kinase 1 (PGK1) protein levels were measured in FLAMs infected with heat-killed *C. neoformans* by western blot analysis. For these experiments, iNos was employed as a positive control since *Nos2* transcripts, which encode iNos, were increased >400 fold following infection with heat-killed *C. neoformans* (Fig. S6B). As expected, iNos proteins appeared abundant in protein samples from FLAMs infected with heat-killed *C. neoformans* but were undetectable in control samples (Fig. 7C). Although *C. neoformans* infection did not affect the expression of LDHA proteins, PGK1 levels were increased by ∼60% over naïve, uninfected cells (Fig. 7C).

Based on both the transcript and protein level data, we hypothesized that *C. neoformans* infection would stimulate increased glycolytic activity in FLAMs but not J774 cells. To test this, we measured proton efflux rate (PER) and oxygen consumption rate (OCR) following *C. neoformans* infection of FLAMs and J774 cells to appraise changes in glycolytic activity and mitochondrial respiration. For these experiments, HKCn was utilized instead of live *C. neoformans* as HKCn had a comparable effect on FLAM glycolytic gene expression and would not interfere with the measurement of host cell metabolic activity. Consistent with our hypothesis, at 24 h post-infection (Fig. 8A-F), mitochondrial respiration in J774 cells showed a modest decrease in activity (Fig. 8A), with no detectable impact on glycolysis (Fig. 8C), and relatively minor effects on the overall rate of ATP production in these cells (Fig. 8E). Whereas, in FLAMs, there was a clear decrease in mitochondrial respiration (Fig. 8B), which was accompanied by a substantial increase in glycolysis (65.77%; Fig. 8D), but no net increase in the overall ATP production rate (Fig. 8F).

**Figure 8:**
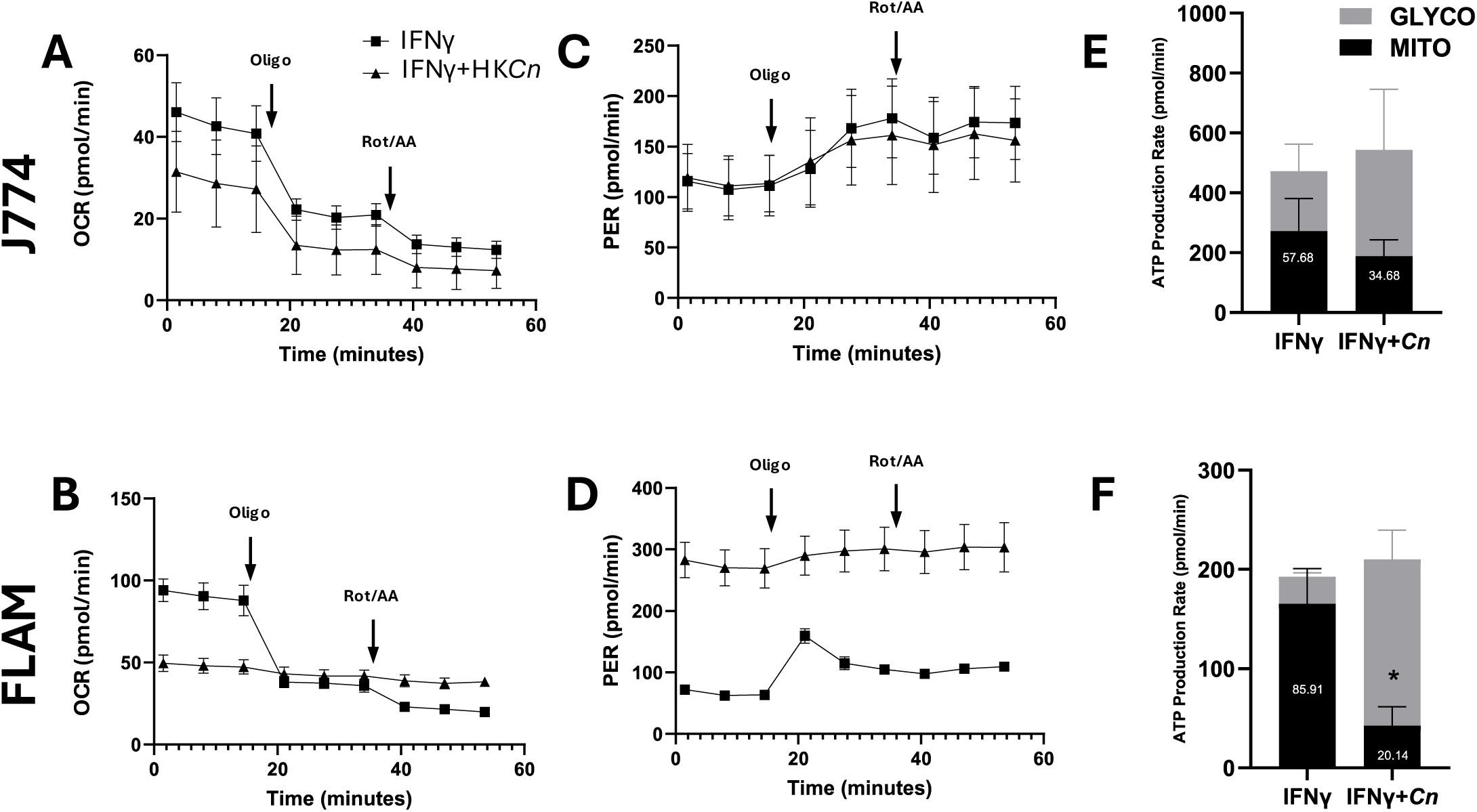
C. neoformans infection induces a glycolytic shift in FLAMs. J774 and FLAMs were M1-polarized by incubation with IFNγ (200 units/mL) for 24 h and then mock-infected (IFNγ) or infected with opsonized heat-killed H99S-eGFP *C. neoformans* (IFNγ+*Cn*) for 2 h before the remaining extracellular yeast were removed. (A+B) The oxygen consumption rate (OCR), (C+D) proton efflux rate (PER), and (E+F) ATP production rate were measured using an XF Real-Time ATP Rate Assay (Agilent). Arrows in (A-D) indicate the timing of oligomycin (Oligo; 1.5µM), or rotenone and antimycin A (Rot/AA; 0.5µM) injection. All experiments were performed as a minimum of three biological repeats. Error is represented as S.E.M. Differences in metabolic activity between the samples were appraised using an unpaired *t*-test with a Welch’s correction and are indicated as follows: \**p*<0.05

### C. neoformans-induced increase in FLAM glycolytic gene expression is HIF1-dependent

As our transcriptome analysis indicated that *C. neoformans* infection increased the levels of RNA for genes in the HIF1 pathway in FLAMs (Supplementary data file 1; Fig. 5E and S2A), we hypothesized that the expression of the glycolytic genes and associated metabolic switching observed in these cells was also HIF1-dependent. To explore this possibility, we used a gain and loss-of-function approach, by either increasing HIF1 activity using the HIF1α-stabilizing agent, dimethyloxalylglycine (DMOG), or blocking HIF1 activity using the highly specific HIF1:chromatin binding inhibitor, echinomycin. Here, we found that DMOG treatment alone was sufficient to induce an almost 200-fold increase in the RNA level of the bona fide HIF1-regulated gene, *Egln3*, and in *Nos2*, which is also known to be HIF1-regulated in a range of cell types, including those of the myeloid lineage ((60–63); Fig. 9A). Consistent with our hypothesis, DMOG treatment also increased RNA levels of the glycolytic genes, *Slc2a1*, *Pgk1*, and *Ldha*. In all cases, IFNγ stimulation did not further enhance the expression changes of these genes. Finally, echinomycin treatment substantially reduced *Egln3*, *Slc2a1*, *Pfkp*, and *Pgk1* transcript levels in heat-killed *C. neoformans-*infected FLAMs (Fig. 9B). Collectively, these data indicate that *C. neoformans* infection stimulates HIF1 activity in FLAMs, and the increases in glycolytic gene expression observed in these cells are at least partially HIF1-dependent.

**Figure 9:**
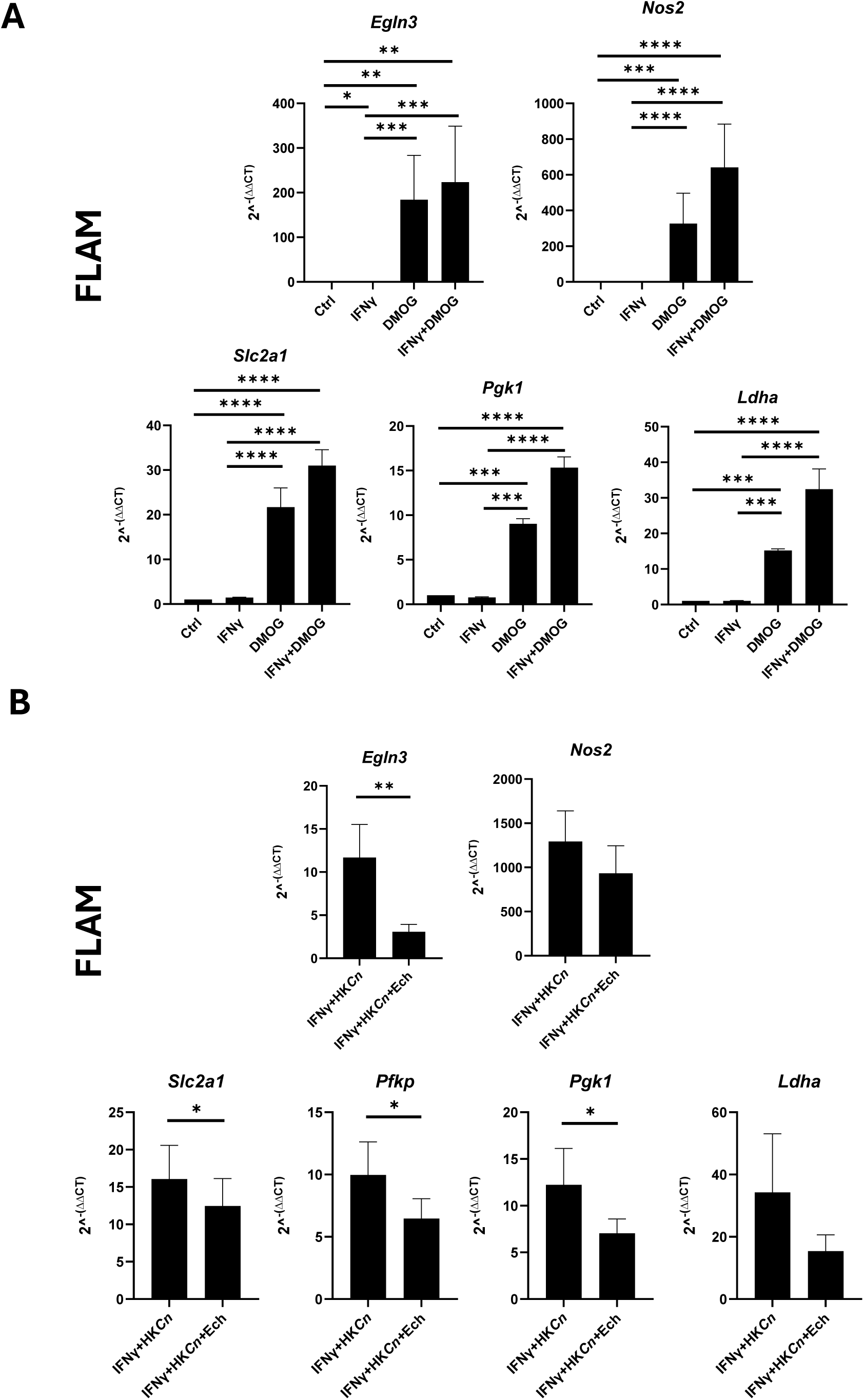
C. neoformans-induced expression of select glycolytic genes is HIF1-dependent. (A) FLAMs were treated with vehicle for 48 h (Ctrl), stimulated with IFNγ (200 units/mL) for 24 h, followed by IFNγ and vehicle for 24 h (IFNγ), treated with vehicle for 24 h, followed by dimethyloxalylglycine (DMOG; 250 μM) for 24 h (DMOG), or stimulated with IFNγ for 24 h followed by IFNγ and DMOG for 24 h (IFNγ+DMOG). The cells were harvested, and the expression level of the indicated transcripts was measured by qRT-PCR. (B) FLAMs were stimulated with IFNγ (200 units/mL) for 24 h and infected with opsonized heat-killed H99S for 2 h before the remaining extracellular yeast were removed and incubated for a further 24 h with vehicle (IFNγ+HK*Cn*) or echinomycin (2.5 nM; IFNγ+HK*Cn*+Ech). The cells were harvested, and the expression of the indicated transcripts was measured by qRT-PCR. All experiments were performed as a minimum of three biological repeats. Error is represented as the S.E.M. Statistical significance was assessed using (A) a one-way ANOVA followed by a Tukey’s multiple comparison test, or (B) a paired *t*-test. Statistical significance is indicated as follows: N.S.; not significant; \**p*<0.05; \*\**p*<0.01; *** *p*<0.001; ****p<0.0001.

## Discussion

While it has long been understood that macrophage populations within the lung play a highly significant role in determining the course and outcome of *C. neoformans* infection (3,64), difficulties associated with maintaining primary AMs in culture have complicated efforts to examine *C. neoformans*:AM interactions in detail. As a potential solution, we investigated whether FLAMs could be used as an AM surrogate for this purpose. Broken into four discrete phases, this study was designed to (i) determine whether the transcriptome profile of FLAMs closely matched that of AMs, (ii) evaluate the responses of these cells to IFNγ stimulation, (iii) examine the transcriptional response of FLAMs to intracellular *C. neoformans* infection, and finally (iv) provide mechanistic insights into any detected transcriptomic and phenotypic changes. For all phases, data developed in FLAMs were compared to analogous datasets produced in other macrophage models, including J774, which is commonly used to study the innate immune response to *C. neoformans* infection (8,11,43,65–68), enabling us to detect differences in behavior that are attributable to FLAMs, and by extension, AMs.

For the first phase of this study, we expanded on prior validation of FLAMs included in Thomas *et al* (29), the first publication describing this model, showing that the steady-state transcriptome profile of FLAMs is more similar to primary AMs than PMs. Here, using our own FLAM transcriptome data sets, we validate this result and show that gene expression in FLAMs is distinct from the macrophage-like cell lines, J774 and RAW264.7 (Fig. 1). The similarities between FLAMs and AMs extend beyond steady-state gene expression. Our metabolic profiling assays indicated that FLAMs, like AMs (20), are heavily dependent on mitochondrial respiration, generating <15% of their total ATP requirements from glycolysis, whereas J774 cells derive ∼40% of ATP from glycolysis (Fig. 2A). Collectively, these data strengthen the case that FLAMs are an appropriate AM model.

For the second phase, we examined the gene expression response of FLAMs to IFNγ stimulation, which enabled us to explore several aspects of macrophage biology that are relevant to AM:*C. neoformans* interactions. Firstly, a robust IFNγ response is crucial for pulmonary protection from *C. neoformans* (69), and is known to enhance fungal clearance in rodent cryptococcosis models, at least in part through the enhancement of macrophage antifungal activity (32,33). Secondly, our prior studies indicate that intracellular *C. neoformans* infection compromises the classical activation of host macrophages by attenuating M1-associated gene expression (9). Thirdly, examination of the AM IFNγ response at the proteome level by other groups indicates that these cells are hyporesponsive to IFNγ (27).

While our transcriptome profiling experiments unambiguously show that both FLAMs and J774 cells are IFNγ-responsive (Fig. 3), with increased steady-state RNA levels of *Stat1*, *Irf1*, and other ISGs seen in these cells following IFNγ stimulation (Supplementary data file 1), several key differences were evident. Although the FLAM IFNγ response was arguably similar in magnitude to that observed in J774 macrophages, based on the number and fold change of DEGs, the identity of these DEGs showed broad differences. The overlap between upregulated DEGs was modest, with <50% of FLAM DEGs common to J774 cells (Fig. 3E). Notably, only J774 cells showed increased levels of the M1 marker *Nos2*. The failure of IFNγ stimulation to increase *Nos2* expression in FLAMs could be due to high basal levels of the nuclear receptor, peroxisome proliferator-activated receptor-γ (PPARγ), a transcriptional regulator required for the proper maturation of tissue-resident AMs (19), which appeared highly expressed in our FLAM transcriptome data (Supplementary data file 1). This transcription factor has been shown to control IFNγ-stimulated *Nos2* transcription in AMs (70), acting as a brake on the inflammatory response to prevent excessive lung inflammation. Given that prior studies suggest AMs exhibit a muted response to IFNγ, these data are unsurprising (27).

The degree of overlap observed between down-regulated DEGs in IFNγ-stimulated FLAMs and J774 cells was even smaller than that observed for the up-regulated DEGs (Fig. 3F). Here, only 6.5% of FLAM DEGs for the Ctrl:IFNγ pairwise comparison were common to the equivalent J774 DEG set and were largely associated with different biological processes. Specifically, GO analysis showed enrichment of genes associated with the cell cycle in FLAMs, but not J774 cells, with these DEGs including *Ccna2*, *Ccnb1, Ccnd1*, and *Cdk1*. Consistent with prior reports showing that IFNγ stimulation induces macrophage cell cycle arrest (71), these data indicate that IFNγ also likely disrupts FLAM cell cycle progression and proliferation.

As a necessary precursor to the third phase, which constitutes the core of this project, we tested the ability of FLAMs to ingest *C. neoformans* and found that these cells phagocytose yeast with similar efficiency to J774 cells. Additionally, time-lapse microscopy of *C. neoformans*-infected cells indicated that the yeast exhibit a slower intracellular replication rate in host FLAMs compared to J774s (Fig. 4G), with 0.38 doublings per hour occurring in J774 cells versus 0.24 in FLAMs (*p*<0.05; Fig. 4H). To address this in a more rigorous manner, we supplemented our manual analysis with a custom mathematical model which compensates for the inherent difficulty of reliably detecting small *C. neoformans* buds by confocal microscopy (Fig. 4I) and assessing population-level growth patterns in the presence of significant individual-cell variability. Data from this analysis does not support host-cell-specific differences in the intracellular proliferation of *C. neoformans.* Distributions of individual cell growth parameters (inter-budding and maturation times) were very similar for the two host cell types. And, while it proved difficult to reliably estimate a population-level maturation time, population-level estimates for the two cell types were similar overall: Inter-budding times for FLAM and J774 cells were within 10 minutes of each other, and at the stable age– and stage-distribution, doubling times for FLAM and J774 cells were 7 and 8.6 hours, respectively. This contrasts with the doubling time calculation based on the growth of the populations over time, which indicated that *C. neoformans* proliferates more slowly in FLAM cells. This difference, however, does not imply a contradiction because doubling times in stage-structured populations depend on the distribution of individuals among life stages, where, in the context of this research, those stage are the immature and mature (actively budding) cell stages. Unlike the initial doubling time analysis, the model-based assessment of growth is based on an analysis of the comprehensive pattern of growth in individual cells, and the model-estimated population doubling time reflects the population’s intrinsic potential for growth, which need not match the doubling time experienced by any sample of the population over a finite window of time. In addition, due to differences in method, the model-based growth analysis was also carried out over a longer window of time than the population doubling time analysis. We also conducted a model-based analysis of nonlytic exocytosis (NLE) events in the two host cell types. Unlike intracellular proliferation, differences in NLE rates were large and significant (likelihood ratios < 0.01), with FLAM cells experiencing NLE events at approximately four times the rate of J774 cells.

Given that our data indicated that the efficiency of *C. neoformans* ingestion and the intracellular growth rates were well-matched between these two macrophage types, we compared the gene expression response of *C. neoformans-*infected FLAMs and J774 cells. In contrast to our prior work, which showed that *C. neoformans* infection partially reversed IFNγ-induced gene expression changes in RAW264.7 macrophages (9), we observed a robust proinflammatory response in FLAMs, with these cells exhibiting increased steady-state RNA of *Nos2*, *Tnf*, and various ISGs (Fig. 5+6). Furthermore, this response was stronger and broader than that seen in J774 cells, with 819 upregulated DEGs detected in FLAMs versus only 302 in J774s (Fig. 5E), and these DEGs typically exhibiting larger fold changes (Fig. 5A+B). While the reasons for this difference are unclear, our RNA-sequencing data show that *Ppar*γ transcript levels decreased 4.24-fold in FLAMs following *C. neoformans* infection. Given the important role played by PPARγ in the control of various AM activities, including inflammation and *Nos2* expression (72), it is plausible that decreased levels of this transcriptional regulator are at least partially responsible for these changes. The expression of *iNos* and other proinflammatory genes seen in *C. neoformans*-infected FLAMs is consistent with this notion (Fig. 6A).

This broader transcriptome response observed in FLAMs was both quantitatively and qualitatively different from that observed in J774 cells, with our GO analysis revealing that distinct biological processes were impacted in the two macrophage types. Perhaps most notably, DEGs associated with the BP term “canonical glycolysis” were upregulated in FLAMs, but not J774 cells (Fig. 5E), and included *Slc2a1*, which encodes the GLUT1 glucose transporter, as well as *Pfkp*, *Gapdh*, *Pgk1*, *Ldha*, and *Eno1*, which encode glycolytic enzymes (Fig. 7B). RNA increases did not necessarily result in increased protein levels of all these enzymes. In fact, our data showed that LDHA levels were not affected, but PGK1 protein levels were increased in FLAMs following infection with heat-killed *C. neoformans* (Fig. 7C). As this pleiotropic enzyme can facilitate metabolic reprogramming of cells by increasing glycolytic flux and inhibiting mitochondrial oxidative phosphorylation (73–75), we hypothesized that *C. neoformans*-induced expression of PGK1 (and other glycolytic regulators) would result in a glycolytic shift in infected FLAMs. To test this hypothesis, we measured changes in OCR and PER in FLAMs and J774 cells following *C. neoformans* infection. To avoid the potential confounding factor of the yeast metabolic activity, heat-killed *C. neoformans* were employed in these assays. As expected, host FLAMs exhibited a substantial increase in the rate of ATP production attributable to glycolysis, with a concurrent decrease in ATP output from the mitochondria of similar magnitude, leaving the overall ATP production rate of these cells unaffected. By contrast, neither glycolytic nor mitochondrial activity was affected in infected J774 cells.

Since various glycolytic genes, including *Pgk1,* in macrophages are sensitive to HIF1α overexpression (76), and *C. neoformans*-infected FLAMs showed increased expression of genes associated with the cellular response to hypoxia (Fig. 5E), a process coordinated by HIF1, we hypothesized that HIF1 was at least partially responsible for the increased glycolytic gene expression detected in these cells. This was explored in the final stage of this study. Here, we were able to show that stabilization of HIF1 proteins, using the pharmacological agent DMOG, was sufficient to enhance glycolytic gene expression, as well as the HIF1-regulated gene *Egln3*, and the M1-marker, *Nos2*. Perhaps more importantly, echinomycin, an inhibitor of HIF1-DNA binding (77), partially reversed *C. neoformans*-induced expression of *Slc2a1*, *Pfkp*, and *Pgk1*, supporting the notion that *C. neoformans*-induced HIF1 activity in host FLAMs is a contributor to the metabolic shift occurring in these cells. As the primary purpose of this study was to test the viability of FLAMs as an *in vitro* model to investigate AM-*C. neoformans* interactions, the consequences of this glycolytic shift are beyond the scope of this work. This result is, however, surprising and demands further exploration in future studies. It is, however, known that during their development, AMs downregulate glycolysis shortly after migrating to the alveoli, becoming relatively metabolically inflexible and highly dependent on mitochondrial respiration. Indeed, the high-oxygen, low-glucose environment provided by the lung is well suited for this mode of ATP generation, and even other macrophage types transferred to the lung environment eventually take on the metabolic characteristics of AMs (78,79). The utilization of mitochondrial respiration may be required for AMs to efficiently carry out effector functions, such as the catabolism of lipid-rich surfactant and performing efferocytosis, to maintain lung tissue homeostasis.

While it is well understood that metabolic shifts favoring glycolysis are a core feature of the response to microbial ligands for macrophages of the monocytic lineage, glycolysis is considered largely dispensable for AM proinflammatory activity, as not only do AMs not undergo a metabolic shift following exposure to LPS, but inhibition of glycolysis also has no effect on the expression of pro-inflammatory cytokines (20). Our initial metabolic profiling data of FLAMs is consistent with this result in the sense that these cells do not increase glycolytic activity when maintained in high glucose media or following IFNγ stimulation (Fig. 2A+C). In both experimental conditions, mitochondrial respiration still accounted for greater than 85% of total ATP without significantly increasing overall energy production, although LPS stimulation did elicit increased PER in these cells (Fig. 2C). However, recent evidence suggests that AMs have a degree of metabolic flexibility and are capable of a metabolic shift under specific environmental circumstances. For example, Woods *et al.* recently demonstrated that AMs undergo glycolytic reprogramming in a model of acute respiratory distress syndrome, which is a hypoxic state, and this effect was HIF1-dependent. Furthermore, the response could be mimicked by non-hypoxic stabilization of HIF1α using the HIF prolyl hydrolase inhibitor, FG-4592 (80). Additionally, Rosa *et al.* showed a Warburg–like shift in lung tissue from Wistar rats infected with *Cryptococcus gatti,* although it was not specifically shown that AMs contributed to this response. Additionally, proteomic analysis of MRC-5 cells (human lung fibroblasts) infected with *C. gatti* revealed an increase in gene expression of several glycolytic enzymes and decreased expression of several TCA cycle proteins (81).

In myeloid-derived macrophages, it is well understood that metabolic shifts favoring glycolysis are a core feature of macrophage activation, or M1 polarization, and pro-inflammatory gene expression following exposure to microbial ligands (47,76,82). However, the impact of this metabolic shift on AMs is less clear as these cells do not conform to the standard M1/M2 paradigm (24,26). Although in this study, we show that increased glycolytic activity in FLAMs is accompanied by a profound increase in the expression of the murine M1 marker, iNOS, at both the transcript and protein level (Fig. 6A and 7C). Given the important role played by iNOS in the production of nitric oxide (NO) and reactive nitrogen species, expression of this enzyme likely enhances the anticryptococcal activity of these cells.

While our study does not provide a mechanism for *C. neoformans*-induced HIF1 activity in FLAMs, several possibilities appear plausible. As high NO levels are known to inhibit the function of prolyl hydroxylases through S-nitrosylation, expression of iNOS in *C. neoformans-*infected FLAMs may stabilize HIF1α during normoxic conditions (83,84). Additionally, *C. neoformans* has been shown to activate NF-κB through both toll-like and C-lectin receptor signaling, and as the *Hif1a* promoter contains κB sites, *C. neoformans* infection may stimulate HIF1 activity in an NF-κB-dependent manner (85).

Our future studies will investigate how HIF1 is regulated in AMs during fungal infection, including the exploration of the two possibilities described above, and the role played by HIF1 in the regulation of AM antimicrobial activities. We will also address several other questions raised by this study. Firstly, it is not entirely clear whether HIF1-dependent expression of core glycolysis enzymes is solely responsible for the metabolic changes observed in FLAMs. We also show that *C. neoformans* infection stimulates expression of the HIF1-regulated gene, *Acod1* (Fig. 6A), which encodes aconitate dehydrogenase I. This enzyme catalyzes the production of itaconate, a metabolite that exerts complex effects on macrophage immunometabolism, impacting both glycolysis and mitochondrial activity, while also affecting the proinflammatory and antimicrobial activities of the cell (86). Secondly, our study shows that *C. neoformans* infection stimulates *Cited1* expression in FLAMs (Fig. 6A), which has also been observed in RAW264.7 cells (9). The transcriptional co-regulator encoded by this gene enhances IFNγ-stimulated gene expression in macrophages (59), and bears a C-terminal conserved region 2 (CR2) domain that is common to all CITED family members. Given that this feature enables CITED2 to function as a repressor of HIF1-dependent pro-inflammatory gene expression in macrophages (87), likely by blocking HIF1-CBP/p300 interactions (53,88), it seems possible that expression of CITED1 may function as a feedback mechanism to control HIF1 activity in FLAMs.

In closing, this study features the first use of FLAMs as an *in vitro* model to investigate AM-*C. neoformans* interactions, introducing a new and versatile tool to the field, which may overcome some of the challenges associated with primary AMs. The data presented here, which suggest a role for HIF1 signaling and enhanced glycolytic activity in the response to intracellular *C. neoformans* infection in AMs, will help to focus future *in vivo* studies and may provide potential new targets for therapeutic intervention.

## Materials & Methods

*Reagents* – Unless otherwise specified, all reagents, cytokines, and consumables were obtained from Thermo Fisher Scientific, Waltham, MA.

*Mammalian Cell Culture –* J774 macrophage-like cells were obtained from ATCC (Manassas, VA) and were cultured in DMEM supplemented with 10% fetal bovine serum (FBS), 25 mM HEPES, 1% nonessential amino acids, and 1% penicillin and streptomycin. FLAMs were produced by the Olive Lab (Michigan State University), as described in Thomas *et al* 2022 (29), and were maintained in culture for no longer than 40 days in RPMI 1640 supplemented with 10% FBS, 30 ng/mL GM-CSF, and 15 ng/mL TGFβ. All cell lines were maintained at 37 °C in a humidified 5% CO_2_ atmosphere. For all western blotting, qRT-PCR, and RNA sequencing experiments, cells were seeded in 6-well plates at a density of 4×10^5^ cells/well in 2 mL of medium and grown to ∼80% confluency prior to treatment with the indicated reagents. Prior to *C. neoformans* infection, macrophages were stimulated by incubation with 200 U/mL recombinant murine IFNγ (Biolegend, San Diego, CA) for 24 hours.

*Yeast Cell Culture and Infection* – The serotype A H99S strain of *C. neoformans* or H99S stably expressing EGFP were cultured in yeast peptone dextrose (YPD) broth with shaking at 37°C for 36 h, washed 3 times in PBS, pelleted by centrifugation at 750 × g, and counted. The cells were opsonized in 20% goat serum (Sigma Aldrich, St. Louis, MO) containing 20 μg 18B7 antibodies targeting the *C. neoformans* capsular polysaccharide glucuronoxylomannan (GXM), per 1.0 × 10^7^ cells in a 1 mL volume. For experiments requiring heat-killed (HK) *C. neoformans*, the yeast was heated to 65°C for 45 minutes prior to opsonization. Unless otherwise specified, macrophages were infected at an MOI of 3:1 (yeast:macrophage) by incubating the macrophages with opsonized yeast for 2 h before removing undigested yeast by washing with PBS before adding fresh growth medium and the indicated cytokines to the cells.

*Immunoblotting –* For immunoblotting experiments, cells were lysed in RIPA buffer containing protease and phosphatase inhibitors (Sigma). The protein concentration of the samples was determined by BCA assay and normalized by dilution with the appropriate lysis buffer. All samples were boiled in 1× Laemmli buffer and resolved by SDS-PAGE and visualized by western blotting.

*Antibodies –* The primary antibodies used for western blotting experiments were as follows: β-actin (A2066, Sigma), iNOS (#13120, Cell Signaling), PGK1 (#68540, Cell Signaling), and LDHA (#2012, Cell Signaling). Primary antibody binding was detected using mouse anti-rabbit IgG-horseradish peroxidase (HRP; sc-2357, Santa Cruz Biotechnology) or anti-mouse m-IgGkappa binding protein (BP)-HRP (sc-516102, Santa Cruz Biotechnology), as appropriate. Membranes were incubated with enhanced chemiluminescent (ECL) reagents and bands were visualized using a ChemiDoc MP Imaging System with Image Lab Software (Bio-Rad, Hercules, CA).

*RNA extraction and RNA sequencing* – Total RNA was extracted using RNeasy Mini Kits (Qiagen) with genomic DNA removed from samples using RNase-Free DNase kits (Qiagen) for on-column DNA digestion. Clean RNA was resuspended in 10 μL diethyl pyrocarbonate (DEPC)-treated water and shipped to Novogene (Sacramento, CA). Here, RNA quality was appraised using an Agilent 2100 Bioanalyzer and samples with an acceptable RNA integrity number were used for cDNA library production and RNA sequencing using the HiSeq 2500 system to produce 150 bp transcriptome paired-end reads.

*Analysis of RNA sequencing data* – Quality checks of fastq RNAseq data files were provided by Novogene. Based on the data quality, no file trimming was necessary. Reads were aligned to the mouse genome (build GRCm39, accession GCA_000001635.9) and a gene level read count was obtained using STAR aligner (version 2.5.3a;(89)). Scaffolding was provided by the mouse reference genome annotation (version 39.90, (90)) in CyVerse Discovery Environment (91). Read count files were joined using multi-join (92), imported into R (version 4.3.2; R Studio version 2023.12 (Ocean Storm)), and samples were clustered based on their whole genome gene expression profile using EdgeR (version 4.0.0; (93)), as described in (94). To evaluate samples for inclusion or exclusion, the data were displayed as a multi-dimensional scale plot. As all samples per condition clustered with their replicate pool (n=3), they were all used in subsequent analyses. To make pairwise comparisons for differentially expressed genes (DEGs) among replicate groups, CuffDiff2 (version 2.2.1; (95)) was used with the same mouse genome annotation and genome files. From the pairwise comparisons, DEGs with fold change ≥ 2.0 and q ≤ 0.05 were considered biologically relevant and statistically significant. Gene ontology (GO) analysis was performed in Database for Annotation, Visualization, and Integrated Discovery (DAVID) 2021 bioinformatics resource tool (version Dec. 2021; (96,97)). GO terms were ranked and plotted as –log(p-value).

Gene set enrichment analysis (GSEA) was performed using GSEA software (version 4.2.3; (98,99)). Read counts derived from STAR output (89) were imported into R and the SARTools R package, DESeq2 version (version 1.7.4; (100)) was used to produce normalized read counts per experiment per sample. Per GSEA recommendations for RNA sequencing experiments (98), genes with no counts in any sample were removed, along with low-expressing genes (mean or geometric mean <10 reads across all samples).

*Pearson Correlation* – These analyses were performed for two groups of genes, one representing an AM transcriptome signature, and another for glycolytic genes. The AM transcriptome signature genes were obtained from Gautier *et al* (2012) figures 2b (genes specifically upregulated in alveolar macrophages), 2c (transcription factors specifically upregulated in alveolar macrophages), and supplemental figure 2 (genes specifically downregulated in alveolar macrophages) (101). Gene names were converted to Ensembl Gene IDs using genome tools at the Mouse Genome Informatics website (https://www.informatics.jax.org/; (102,103)) resulting in a 155 gene set. The glycolytic genes were obtained from GO terms “glycolytic process” (GO:0006096), “canonical glycolysis” (GO:0061621), and “regulation of glycolytic process” (GO:0006110), resulting in a 101-gene set. The STAR-SARTools pipeline was used to generate a normalized gene count file for all samples to be compared and a jupyter notebook (version 7.0.8) was coded to perform the following steps for each gene group: First, genes with no counts across all samples were removed and 1 was added to all samples in order to calculate a geometric mean for each row/gene. Any row with a geometric mean <10 was dropped. The AM signature gene set or glycolytic gene set was merged with the gene expression data table to produce a table of AM signature or glycolytic gene expression across the samples. A single count was added to all values, and the geometric mean of replicates was calculated. The geometric mean of counts per gene was used to calculate a Pearson’s correlation matrix for each gene group and visualized as color correlation plots with a scale of –1 (purple) to 1 (green). Additionally, genes encoding general macrophage markers, AM-specific genes, and PM-specific genes were selected from the literature (29,101,104), and differences between the samples in the expression of these genes were calculated as the log2 of the geometric mean of each replicate group and visualized as a heatmap. Python libraries used were Seaborn (version 0.13.0), Pandas (version 2.1.4), NumPy (version 1.26.4), matplotLib (3.8.0), SciPy (version 1.12.0).

*RT-PCR* – For quantitative RT-PCR (qRT-PCR) experiments, RNA (isolated as described above) was reverse transcribed to produce cDNA libraries using Maxima H Minus Reverse Transcriptase in the presence of dNTPs, Oligo dT primers, and RiboLock RNase inhibitor (ThermoScientific). cDNAs of interest were amplified using appropriate primer pairs and PerfeCTa SYBR Green FastMix (Quantabio) in a CFX Opus 96 Real-Time PCR Instrument (Bio-Rad). Normalization was performed using two reference genes, *Actb and Cyc1* (*Actb* F-CACTGTCGAGTCGCGTCC, R-TCATCCATGGCGAACTGGTG, *Cyc1* F-CTAACCCTGAGGCTGCAAGA, R-GCCAGTGAGCAGGGAAAATA). Where possible, primers were designed such that at least one primer for each pair spanned an exon-exon boundary and produced a product size between 70 and 150 bp. The sequences (5’ to 3’) of primer pairs used were as follows: *Acod1:* F-AGTTTTCTGGCCTCGACCTG, R-AGAGGGAGGGTGGAATCTCT, *Cited1:* F-CTGCCACCGATTTATCGGACTT, R-CTCCTGGTTGGCATCCTCCTT, *Cited2:* F-GCAAAGACGGAAGGACTGGA, R-CGTAGTGTATGTGCTCGCCC, *Egln3:* F-AAGGAGCGGTCCAAGGCAAT, R-ATACAGCGGCCATCACCATT, Eno1: F-TGTACACCGCAAAAGGTCTCT, R-CCCATGAAGCGGGTCTTATCA, Gapdh: F-CCCTTAAGAGGGATGCTGCC, R-TACGGCCAAATCCGTTCACA, Hif1a: F-GACAGAGCCGGCGTTTAGG, R-CGACGTTCAGAACTCATCCTATTTT, Hmox1: F-ACCTTCCCGAACATCGACAG, R-CAGCTCCTCAAACAGCTCAATG, Il-6: F-CTCTGCAAGAGACTTCCATCCA, R-GACAGGTCTGTTGGGAGTGG, Ldha: F-AGTAAGTCCTCAGGCGGCTA, R-GGACTTTGAATCTTTTGAGACCTTG, Nos2: F-GGTGAAGGGACTGAGCTGTT, R-ACGTTCTCCGTTCTCTTGCAG, Pdrx1: F-TGTCCCACGGAGATCATTGC, R-GGGTGTGTTAATCCATGCCAG, Pfkp: F-ATTGGAGGATTTGAGGCCTACC, R-AGGAACCATAACCATGGGGAC, Pgk1: F-CCACAGAAGGCTGGTGGATT, R-GTCTGCAACTTTAGCGCCTC, Slc2a1: F-GATCCCAGCAGCAAGAAGGT, R-TAGCCGAACTGCAGTGATCC, and Tnfa: F-TAGCCCACGTCGTAGCAAAC, R-GCAGCCTTGTCCCTTGAAGA.

*Fluorescence Microscopy –* FLAMs or J774 cells were plated at a density of 4.0×10^5^/dish with 2 mL growth medium in 35 mm glass-bottom dishes (Cellvis, USA) and incubated with 200 units/mL IFNγ for 24 h prior to infection with H99S-GFP opsonized with complement and 18B7. The cells were incubated with the yeast for 2 h at an MOI of 0.5:1 before extracellular yeast were removed by washing with growth medium. Fresh medium was added containing calcofluor white (CFW; 7.5 μg/mL) and propidium iodide (PI; 7.5 μg/mL) to label extracellular yeast and dead cells, respectively, and the infected macrophages were imaged immediately using a Zeiss LSM700 confocal laser scanning microscope equipped with a Plan-Apochromat 63x magnification/1.40 numerical aperture oil immersion DIC M27 objective lens and controlled using Zen software (Zeiss). On track 1, EGFP fluorescence was excited using a 488 nm laser. On track 2, CFW and PI fluorescence were imaged concurrently using a 405 nm laser to excite CFW fluorescence and a 550 nm laser to excite PI fluorescence. A 544 nm beamsplitter was used to send CRW fluorescence to PMT1, which was detected through a 555 nm short-pass filter, and PI fluorescence was sent to PMT2 and detected through a 560 nm long-pass filter. Images were recorded at 30-minute intervals for a minimum of 11 hours across a minimum of seven different positions per sample as z-stacks. To accommodate differences in cell thickness, J774 images were collected as 5 z-slices at 2 μm intervals across an 8 μm range, and FLAM images were collected as 7 z-slices at 2 μm intervals across a 12 μm range. Image files were viewed in Fiji (105), and the number of intracellular yeast in each host macrophage was recorded at each time point. To normalize the data, the number of intracellular yeast in a host cell was divided by the initial number in that same cell at t=0 h. Host cells were culled from the analysis if: 1) they did not remain entirely visible within the field, 2) contained less than one but more than two intracellular EGFP-positive *C. neoformans* at t=0 h, 3) lysed or divided, or the intracellular *C. neoformans* were lost by NLE or were transferred to another host cell within the first 3 h of the timecourse. To quantify NLE, images were analyzed at each time point, and a count was made for each cell that met the inclusion criteria for rate analysis. If a cell showed a reduction in intracellular *C. neoformans* of at least one yeast that was sustained for a minimum of two consecutive images without fungal or macrophage death, the cell was designated as having undergone a NLE event.

*Metabolic Profiling* – To measure baseline ATP production rate, FLAMs and J774 cells were plated at a density of 12.5×10^4^ per well in Seahorse XF cell culture plates, and measurements were taken 24 h later. To measure changes in metabolic flux, cells were plated at a lower density to account for extra days in culture due to the experimental design. In this case, FLAMs and J774s cells were plated at 1.0×10^4^ or 5×10^3^ per well, respectively. Cells were treated and/or infected as described with heat-killed *C. neoformans*, and OCR and PER rates were measured using an XFp Real-Time ATP Rate Assay Kit and Seahorse XF HS Mini Analyzer (Agilent Technologies, USA). Total ATP values were normalized to cell counts to account for well-to-well variation in cell density. To measure real-time metabolic shifts associated with macrophage activation, FLAMs or J774 cells were plated at a density of 1.5×10^4^ per well in Seahorse XF cell culture plates. At 24 hrs post-plating, PER was measured by following a modified version of the Agilent Technologies Application Note, “Real Time Discrimination of Inflammatory Macrophage Activation Using Agilent Seahorse XF Technology”. In brief, the plates were loaded into the Seahorse XF HS Mini Analyzer, and PER was measured at 6-minute intervals over the entire duration of the experiment with IFNγ (200 units/mL), LPS (100 ng/mL; *Salmonella enterica* serotype typhimurium; Sigma Aldrich), or IFNγ and LPS injected at 18 minutes to stimulate macrophage activation. At 120 minutes post-activation, the glycolysis inhibitor, 2-deoxyglucose (DG; 50mM), was injected, and a further four measurements were recorded.

*Statistical analysis* – All experiments were performed as three discrete biological repeats unless otherwise stated. Statistical analyses were performed in GraphPad Prism 7 (GraphPad, USA) using the tests indicated in the figure legends.

*Modeling Intracellular Population Growth –* A mathematical model of population growth via asymmetric budding was developed for comparison of intracellular proliferation in J774 and FLAM cells. The model includes two dependent variables, *B*(*t,r*) and *M*(*t,r,a*), which record the number of daughter and mother cells, respectively, and two growth parameters, *T_b_* and *T_m_*, which represent the time between budding events in a mother cell and the time between emergence and first budding in a daughter cell, respectively. Mother cells are associated with three independent variables: *t*, time, *r*, reproductive age, and *a*, age relative to entry into the mother stage. Reproductive age and absolute age are tracked separately for mother cells since the former determines the timing of budding events and the latter determines cell size. Since daughter cells have yet to bud, the reproductive and absolute age of daughter cells are identical. The evolution of the model variables through time is governed by the following system of partial difference equations and boundary conditions.

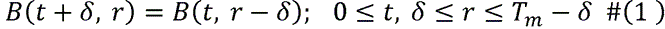

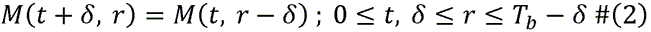

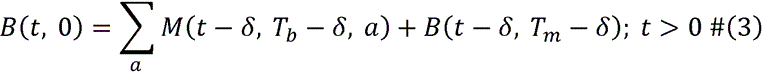

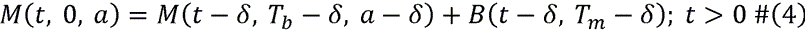

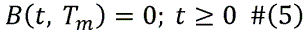

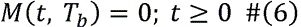

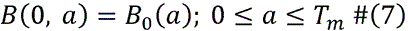

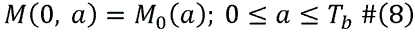

*Modeling Error –* The deterministic model of cell growth is complemented by a probabilistic model of error to allow for maximum likelihood parameter estimation. We consider three types of error. First, small cells can be caught between z-slices. Second, cells that are aligned vertically can be confounded. Finally, there is a potential for miscount due to human or microscope errors. The model of error is informed by details of the microscope settings, spherical geometry of yeast cells, and empirical counting rules. Since this model derivation is technical, details are presented in the appendix.

*Derivation of the stable age– and stage-distributions* – While the doubling time for the population varies with initial conditions and through time, the model nevertheless possess a stable age– and stage-distribution which features a constant doubling time. This stable doubling time provides a convenient measure of the population’s intrinsic potential for growth.

The stable age– and stage-distributions are derived as follows: Substituting *B*(*t,r*) = *f*(*t*)*a*(*r*) into equation (1) we find that

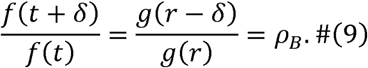

This yields 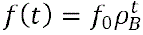 and 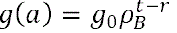, from which it follows that

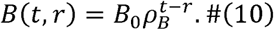

Similarly,

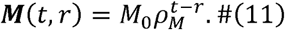

Simultaneously enforcing (10)-(11) and (3)-(4) yields *B*_0_ = *M*_0_, *p_B_* = *p_M_*, and

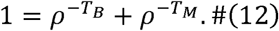

Equation (12) determines *p*, from which the stable doubling time is computed:

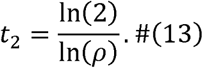

The stable stage distributions can also be found. The stable fraction of immature daughter cells is

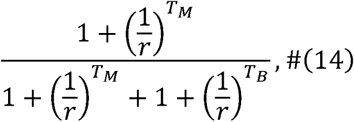

*Model Validation and Parameterization –* The descriptive ability of the model was verified by visual inspection of model fits to individual cell counts (see supplementary mathematical Figure 11 for samples) and by consideration of normalized negative log likelihood values of the fits (see Mathematical Appendix Fig. 12 and 13). In all but a handful of J774 cells, negative log likelihood values for individual cells were in the range of [0,1], which corresponds to a geometric mean for the probability of a count in the range of [0.36, 1]. Using the probabilistic model of error, we perform maximum-likelihood parameter estimation (MATLAB, fminsearch) to parameterize the growth process in individual macrophages. Details of our numerical method are available in the appendix, and the code is available at the GitHub repository.

*Estimating Cell Growth and Size-T*o characterize the growth rate of small yeast, several (*n*= 13) small yeast cells with good visibility were identified. Zen Lite Software was used to measure the radius of the largest cross section of each cell as an estimate of its radius. Each cell was followed through time to obtain a time series of radii with at least three observation times. Preliminary visual inspection of the resulting time series of radii led us to select a linear model for the growth of a cell’s radius. The linear growth rate, λ, for each cell was estimated by minimizing the sum of the squared errors (see Mathematical Appendix Fig. 10). The arithmetic mean of the growth rates for individual cells was used as the radial growth rate for the population. Using this procedure, growth rates of λ = 0.012 *µm mim*^-1^ and λ = 0.016 *µm mim*^-1^ were estimated for J774 and FLAM cells, respectively, with standard deviations of 0.005 *µm mim*^-1^ and 0.008 *µm mim*^-1^ in J774 and FLAM cells, respectively.

A representative size for a mature mother cell (*R*_2_) was obtained by estimating the radii of yeast that were actively budding. Since this analysis focuses on large cells, two successive cross sections of each cell were measured to obtain a better estimate the cell’s radius, as described in Davis, 2025 (106). Using this procedure, the radius of a mature mother cell was estimated at the 2nd and 3rd observed budding events. We thus collected two radii estimates from 10 mature yeast cells, and then averaged the 20 radii, to estimate an average radius of a mother cell. The average radius of a mother cell was 3.2 μm for J774 and 3.4 μm for FLAMs, with a standard deviation of 0.12 μm and 0.17 μm in J774 and FLAM cells, respectively.

*Modeling NLE Events –* Host cell types exhibit differences in the timing and fractions of cells experiences nonlytic exocytosis (NLE) and other events (OTHER). To measure the significance of these differences we constructed an event time model where event times are determined by multi-step, competing Poisson processes. In this model, cell progress from one stage of the process to the next, and from select stages into OTHER and NLE fates at constant rates. Let *x_i_*(*t*) represent the probability that the cell is in state *i* at time *t*, *N*(*t*) represent the probability the cell experienced an NLE event before time *t*, and *0*(*t*) represent the probability the cell experienced an OTHER event before time *t*, all conditional on the initial state of the cell.

We initially considered models where NLE and OTHER events occur from any stage but found a model where NLE events only occur in the final stage provides competitive fits while having fewer parameters. The model is as follows:

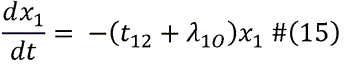

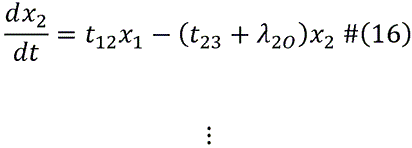

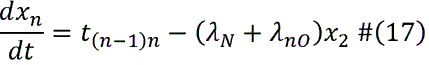

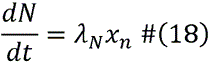

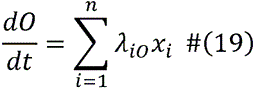

This model is exactly solvable as a system of coupled, linear, differential equations with constant coefficients. Nonetheless, the system was solved numerically in MATLAB (ode45.m).

The case where NLE and OTHER events follow exponential distributions is a special case of the above model where *n* = 1.

We used the Akiake information criterion

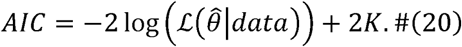

where *K* is the number of parameters (107) (MathWorks, 2025) to compare the ability of models with 1-4 stages to fit events times from both cell lines. To assess the significance of differences in NLE events between cell lines, the relative likelihood of the best-fit models for each host cell type were computed as

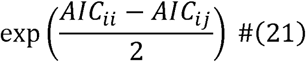

Where *AIC_ii_* denotes the minimal value of the AIC achieved by the best-fit model for host type *i* and *AIC_ij_* denotes the minimal value of the AIC achieved by the best-fit model for host type *i* with the constraint that rates of NLE events match those of the best-fit model for host type *j*, that is, where optimization is performed over parameters governing exit to other events.

The descriptive ability of the model for J774 and FLAM event-time data is illustrated in Mathematical Appendix Fig. 14-17.

## Availability of data and materials

The original RNA sequencing files have been deposited in NCBI GEO under the BioProject ID PRJNA1284885. All other data generated or analyzed during this study are included in this published article. All non-commercial plasmid constructs used in this study are available on request from David E. Nelson.

## Supporting information

Math modeling appendix

RNAsequencing data

## Acknowledgments

We thank the Middle Tennessee State University Molecular Biosciences PhD program and Undergraduate Research Center for funding and support.

## Author contributions

DEN conceived, designed, and coordinated this study, wrote and edited the manuscript, and obtained funding. DAW contributed to the writing and editing of the manuscript, prepared all samples used for RNAseq-based transcriptome profiling (unless otherwise indicated), and performed and analyzed all experiments except those attributed to other co-authors. JBG performed and analyzed the experiments shown in Figs. 7C and S7. RLST assisted with designing the RNAseq assays, performed the bioinformatic analysis of the data used for Figs. 1, 3, 5, 7A, and S2-4, and obtained funding. RNL, AED, and MEC constructed the mathematical model to simulate intracellular *C. neoformans* replication and produced the data featured as Fig. 4I. EEM assisted with the design and coordination of the study and obtained funding. CHK assisted with the experiments shown in Fig. 7C, ENC assisted with the experiments shown in Fig. 4, AME assisted with the bioinformatic analysis of the data used for Figs. 1A, 3, and 4, and performed and analyzed the experiments shown in Fig. S5, and CAH assisted with the experiments shown in Fig. 9. All authors reviewed the results and approved the final version of the manuscript.

## Consent for publication

Not applicable.

## Ethics approval and consent to participate

Not applicable.

## Figure Legends

**Supplemental Figure S1: ifNγ stimulation alone does not affect the balance of glycolysis to mitochondrial respiration in FLAMs and J774 macrophage-like cells.** J774 and FLAM cells were left untreated (Ctrl) or stimulated with IFNγ (200 units/mL) for 24 h. ATP production was measured using an XF Real-Time ATP Rate Assay (Agilent). Differences in metabolic activity between the samples were appraised using a Welch’s *t*-test and found to be not significant.

**Supplemental Figure S2: Transcriptional responses of FLAMs and J774 macrophage-like cells to intracellular *C. neoformans* infection.** (A) FLAM and (B) J774 cells were left untreated (Ctrl) or M1-polarized by incubation with IFNγ (200 units/mL) for 24 h and then mock-infected (IFNγ) or infected with opsonized H99S-eGFP (IFNγ*Cn*) and then harvested for RNA sequencing analysis at 24 h post-infection. Common and differentially expressed genes from Ctrl:IFNγ and Ctrl: IFNγ*Cn* pairwise comparisons are represented as Venn diagrams. Gene ontology (GO) analysis was performed for each gene set in DAVID and the top seven biological process (BP) terms are displayed ranked by –log(*p*-value). The number of genes associated with each term is reported, and all GO terms are significant (False discovery rate <0.5%) except those below the dashed blue line where this is present. GO terms of special interest are marked in red.

**Supplemental Figure S3: Transcriptional response of FLAMs to intracellular *C. neoformans* infection.** (A+B) FLAM cells were left untreated (Ctrl) or stimulated with IFNγ (200 units/mL) for 24 h and then mock-infected (IFNγ) or infected with opsonized H99S-eGFP *C. neoformans* (IFNγ*Cn*) and then harvested for RNA sequencing analysis at 24 h post-infection. Common and differentially expressed genes from Ctrl:IFNγ and Ctrl:IFNγ*Cn* pairwise comparisons are represented as Venn diagrams for (A) upregulated and (B) downregulated genes. Gene ontology (GO) analysis was performed for each gene set in DAVID and the top seven biological process (BP) terms are displayed ranked by –log(*p*-value). The number of genes associated with each term is reported, and all GO terms are significant (False discovery rate <0.5%) except those below the dashed blue line where this is present. GO terms of special interest are marked in red.

**Supplemental Figure S4: Transcriptional response of J774 cells to intracellular *C. neoformans* infection.** (A+B) J774 cells were left untreated (Ctrl) or stimulated with IFNγ (200 units/mL) for 24 h and then mock-infected (IFNγ) or infected with opsonized H99S-eGFP *C. neoformans* (IFNγ*Cn*) and then harvested for RNA sequencing analysis at 24 h post-infection. Common and differentially expressed genes from Ctrl:IFNγ and Ctrl:IFNγ*Cn* pairwise comparisons are represented as Venn diagrams for (A) upregulated and (B) downregulated genes. Gene ontology (GO) analysis was performed for each gene set in DAVID, and the top seven biological process (BP) terms are displayed ranked by –log(*p*-value). The number of genes associated with each term is reported, and all GO terms are significant (False discovery rate <0.5%) except those below the dashed blue line, where this is present. GO terms of special interest are marked in red.

**Supplemental Figure S5: ifNγ stimulation and *C. neoformans* infection have a minimal effect on glycolytic gene expression in J774 cells.** J774 cells were left untreated (Ctrl) or stimulated with IFNγ (200 units/mL) for 24 h and then mock-infected (IFNγ) or infected with opsonized H99S (IFNγ+*Cn*). The expression of transcripts encoding glycolytic enzymes and regulators was quantified by qRT-PCR. Results are expressed as the mean of at least three biological repeats. Error is represented as the S.E.M. Statistical significance assessed using a one-way ANOVA followed by a Tukey’s multiple comparison test and is indicated as follows: \**p*<0.05; \*\**p*<0.01; ****p<0.0001.

**Supplemental Figure S6: Heat-killed *C. neoformans* infection stimulates the expression of proinflammatory and glycolytic genes in FLAMs.** (A) FLAM and (B) J774 cells were left untreated (Ctrl) or stimulated with IFNγ (200 units/mL) for 24 h and then mock-infected (IFNγ) or infected with heat-killed opsonized H99S-eGFP *C. neoformans* (IFNγ+HK*Cn*). Expression of the indicated transcripts was quantified by qRT-PCR. Results are expressed as the mean of at least three biological repeats. Error is represented as the S.E.M. Statistical significance assessed using a one-way ANOVA followed by a Tukey’s multiple comparison test and is indicated as follows: \**p*<0.05; \*\**p*<0.01.

**Supplemental Figure S7: DMOG treatment stimulates PGK1 expression in FLAMs.** FLAMs were treated with vehicle for 48 h (Ctrl), stimulated with IFNγ (200 units/mL) for 24 h followed by IFNγ and vehicle for 24 h (IFNγ), treated with vehicle for 24 h followed by dimethyloxalylglycine (DMOG; 250 μM) for 24 h (DMOG), or stimulated with IFNγ for 24 h followed by IFNγ and DMOG for 24 h (IFNγ+DMOG). The cells were harvested and, PGK1 and LDHA protein levels were measured by western blotting. Actin was utilized as a loading control. Western blot images shown are a representative example from three discrete biological repeats. Results are expressed as the mean of at least three biological repeats. Error is represented as the S.E.M. Statistical significance assessed using a one-way ANOVA followed by a Tukey’s multiple comparison test and is indicated as follows: \**p*<0.05; \*\**p*<0.01.

**Mathematical Appendix Figure 1: Illustration of Derivation of Conditions for Type I Countability.** The depth of the focal z slice (black line) into the mother cell (*h*_2_) is expressed as a function of the depth of the slice into the daughter cell (*h*_l_) using similar triangles.

**Mathematical Appendix Figure 2: Illustration of Range of Simultaneous Visibility of Mother and Daughter in Type I Countability.** Here the depth of the *z* slice, *h*, is measured relative to the top of the daughter cell. The range of visible depths for the mother is *L*_2_ < *h* < *U*_2_, while that is the daughter is *L*_l_ < *h* < *U*_l_,. The overlap of the regions is seen in gray. z slices within the gray region yield simultaneous visible cross sections of both cells enabling the daughter cell to be counted.

**Mathematical Appendix Figure 3: Illustration of Type 2 Countability.** Panels illustrate possible successive cross sections of a mother daughter pair. As we move through successive *z* slices, the mother cell drops out of view as the daughter comes into view. The concavity of the cell radius as a function of the depth of the *z* slice let’s us distinguish daughter from mother in this case. This figure is for illustrative purposes only and is not drawn to scale.

**Mathematical Appendix Figure 4: Illustration of Countability Regions in Cases A-C**. Panels depict countability cases A-C from left to right. The daughter cell’s potential to be counted changes as she grows. In cases A and B, dark pink denotes regions where the small daughter is obscured by the mother. In case C, pale blue and pale green denote regions where the large daughter is always counted. This figure is for illustrative purposes only and is not drawn to scale.

**Mathematical Appendix Figure 5: Illustration of numerical method, part 1.** The model is solved in the light gray regions *R*_1_ and *R*_2_ where the reproductive age, *r*, of cells is greater than time, *t*, by following the initial data only the characteristic lines. Sample characteristic lines are illustrated in purple. This procedure yields solution values along the boundaries *r* = *T_m_* for daughter cells (orange) and *r* = *T_b_* for mother cells (various colors) which are used to initialize the boundary *r* = 0 for both mother and daughter cells (various colors).

**Mathematical Appendix Figure 6: Illustration of numerical method, part 2.** To prevent double counting, cell counts on the boundaries *r* = *T_m_* and *r* = *T_b_* are zeroed out (red). The model is solved in the light blue regions *R*_3_ and *R*_4_ where r< t by following the newly budded cells along the characteristic lines. Sample characteristic lines are illustrated in purple. This procedure yields solution values along the entire upper boundary *t* = *T_b_* (dashed black line) which are used to initialize the next iteration of the algorithm.

**Mathematical Appendix Figure 7: Convergence Behavior with Increasing Initial Guesses.** The likelihood values of numerical solutions to the parameter optimization routine for increasing numbers of initial guesses are compared using the convergence measure for the routine (see equation (19), where the tolerance for convergence is 10^-3^. The height of the bars gives the value of the convergence measure between solutions, where positive values indicate improvements in likelihood. The number of initial parameter guesses for the solution is represented as a sum *x* + *y*, where x denotes the number of choices for the pair (*T_b_*, *T_m_*) and y denotes the number of choices for the initial age of a cell, *a*_0_. Hence, each cell is fit using a total of *xy* initial guesses for the parameters. A small number of cells (n=5) exhibit low variability in the solution likelihood as the number of initial guesses increases (convergence measure <10^-2^).

**Mathematical Appendix Figure 8: Effect of Number of Initial Guesses on Likelihood Ratios.** The likelihood values of numerical solutions to the parameter optimization routine for increasing numbers of initial guesses are compared using the likelihood ratios. Values greater than one indicate improvement in likelihood. The number of initial parameter guesses for the solution is represented as a sum *x* + *y*, where x denotes the number of choices for the pair (*T_b_*, *T_m_*) and y denotes the number of choices for the initial age of a cell, *a*_0_. Hence, each cell is fit using a total of *xy* initial guesses for the parameters. A small number of cells exhibit low variability in the solution likelihood (∼1%) as the number of initial guesses varies.

**Mathematical Appendix Figure 9: Parameter Convergence Under Increasing Initial Guesses.** The parameter values for numerical solutions to the parameter optimization routine for increasing numbers of initial guesses are compared. Visible red and blue dots indicate variability in the numerical solution’s parameters. The number of initial parameter guesses for the solution is represented as a sum *x* + *y*, where x denotes the number of choices for the pair (*T_b_*, *T_m_*) and y denotes the number of choices for the initial age of a cell, *a*_0_. Hence, each cell is fit using a total of *xy* initial guesses for the parameters.

**Mathematical Appendix Figure 10: Estimating the Growth Rate of Daughter Cells.** Illustration of the method of estimating the linear growth of the daughter cell radius.

**Mathematical Appendix Figure 11: Sample Fits of the Mathematical Model to Individual J774 Cells.** Model population size (blue stars) are shown along with empirical counts (red circles). We see counts often underestimate the model population size, due in part to the limited visibility of small daughter cells. Right panel: Cell 10 demonstrates how the empirical count sometimes falls, presumably due to limited visibility of small daughters.

**Mathematical Appendix Figure 12: Plots of Normalized Negative Log Likelihood Values for Numerical Fits of the Mathematical Model to Individual J774 and FLAM cells.** Individual cell counts that include fewer than 2 complete inter-budding times, that is, fail to achieve a count of at least four cells, are excluded from the individual-cell analysis since they contain very limited information for estimating inter-budding and maturation times. Counts are truncated once they achieve a count of six to minimize miscounts due to cell crowding. Cells are arranged by the length of their count data in descending order.

**Mathematical Appendix Figure 13: Scatter Plots of Individual-Cell Normalized Negative Log Likelihood Values for Numerical Fits of the Mathematical Model to a Population.** The descriptive ability of population-level parameter estimates for individual cell counts is quantified by the normalized negative log likelihood of the population parameters for each individual cell count. Population-level parameter estimation was carried out by minimizing the negative log likelihood of individual observations, under the assumption that the population was homogeneous. In the case of J774 cells, outlier cells 2 and 3 were excluded from the population-level analysis. Cells are arranged by the length of their count data in descending order. Counts are truncated once they achieve a count of six to minimize miscounts due to cell crowding. Cells are arranged by the length of their count data in descending order.

**Mathematical Appendix Figure 14: Comparison of Model and Empirical NLE Time Probability Density and Cumulative Density Functions in J774 Cells**.

**Mathematical Appendix Figure 15: Comparison of Model and Empirical OTHER Time Probability Density and Cumulative Density Functions in J774 Cells**.

**Mathematical Appendix Figure 16: Comparison of Model and Empirical NLE Time Probability Density and Cumulative Density Functions in FLAM Cells**.

**Mathematical Appendix Figure 17: Comparison of Model and Empirical OTHER Time Probability Density and Cumulative Density Functions in FLAM Cells**.

## References

1. Rajasingham, R., Govender, N. P., Jordan, A., Loyse, A., Shroufi, A., Denning, D. W., Meya, D. B., Chiller, T. M., and Boulware, D. R. (2022) The global burden of HIV-associated cryptococcal infection in adults in 2020: a modelling analysis. Lancet Infect Dis 22, 1748–1755

2. Mitchell, T. G., and Perfect, J. R. (1995) Cryptococcosis in the era of AIDS--100 years after the discovery of Cryptococcus neoformans. Clin Microbiol Rev 8, 515–548

3. Osterholzer, J. J., Milam, J. E., Chen, G. H., Toews, G. B., Huffnagle, G. B., and Olszewski, M. A. (2009) Role of dendritic cells and alveolar macrophages in regulating early host defense against pulmonary infection with Cryptococcus neoformans. Infection and immunity 77, 3749–3758

4. Zafar, H., Altamirano, S., Ballou, E. R., and Nielsen, K. (2019) A titanic drug resistance threat in Cryptococcus neoformans. Curr Opin Microbiol 52, 158–164

5. Melhem, M. S. C., Leite Junior, D. P., Takahashi, J. P. F., Macioni, M. B., Oliveira, L., de Araujo, L. S., Fava, W. S., Bonfietti, L. X., Paniago, A. M. M., Venturini, J., and Espinel-Ingroff, A. (2024) Antifungal Resistance in Cryptococcal Infections. Pathogens 13

6. Iyer, K. R., Revie, N. M., Fu, C., Robbins, N., and Cowen, L. E. (2021) Treatment strategies for cryptococcal infection: challenges, advances and future outlook. Nat Rev Microbiol 19, 454–466

7. Fisher, M. C., Hawkins, N. J., Sanglard, D., and Gurr, S. J. (2018) Worldwide emergence of resistance to antifungal drugs challenges human health and food security. Science 360, 739–742

8. Coelho, C., Souza, A. C., Derengowski, L. D., de Leon-Rodriguez, C., Wang, B., Leon-Rivera, R., Bocca, A. L., Goncalves, T., and Casadevall, A. (2015) Macrophage Mitochondrial and Stress Response to Ingestion of Cryptococcus neoformans. Journal of immunology

9. Subramani, A., Griggs, P., Frantzen, N., Mendez, J., Tucker, J., Murriel, J., Sircy, L. M., Millican, G. E., McClelland, E. E., Seipelt-Thiemann, R. L., and Nelson, D. E. (2020) Intracellular Cryptococcus neoformans disrupts the transcriptome profile of M1– and M2-polarized host macrophages. PloS one 15, e0233818

10. Hayes, J. B., Sircy, L. M., Heusinkveld, L. E., Ding, W., Leander, R. N., McClelland, E. E., and Nelson, D. E. (2016) Modulation of Macrophage Inflammatory Nuclear Factor kappaB (NF-kappaB) Signaling by Intracellular Cryptococcus neoformans. The Journal of biological chemistry 291, 15614–15627

11. Tucker, S. C., and Casadevall, A. (2002) Replication of Cryptococcus neoformans in macrophages is accompanied by phagosomal permeabilization and accumulation of vesicles containing polysaccharide in the cytoplasm. Proceedings of the National Academy of Sciences of the United States of America 99, 3165–3170

12. Zhang, L., Zhang, K., Li, H., Coelho, C., de Souza Goncalves, D., Fu, M. S., Li, X., Nakayasu, E. S., Kim, Y. M., Liao, W., Pan, W., and Casadevall, A. (2021) Cryptococcus neoformans-Infected Macrophages Release Proinflammatory Extracellular Vesicles: Insight into Their Components by Multi-omics. mBio 12

13. Kalem, M. C., Humby, M. S., Wohlfert, E. A., Jacobs, A., and Panepinto, J. C. (2021) Cryptococcus neoformans Coinfection Dampens the TNF-alpha Response in HIV-1-Infected Human THP-1 Macrophages. mSphere 6

14. Ginhoux, F., and Guilliams, M. (2016) Tissue-Resident Macrophage Ontogeny and Homeostasis. Immunity 44, 439–449

15. Ginhoux, F., Schultze, J. L., Murray, P. J., Ochando, J., and Biswas, S. K. (2016) New insights into the multidimensional concept of macrophage ontogeny, activation and function. Nature immunology 17, 34–40

16. Evren, E., Ringqvist, E., and Willinger, T. (2020) Origin and ontogeny of lung macrophages: from mice to humans. Immunology 160, 126–138

17. Guilliams, M., De Kleer, I., Henri, S., Post, S., Vanhoutte, L., De Prijck, S., Deswarte, K., Malissen, B., Hammad, H., and Lambrecht, B. N. (2013) Alveolar macrophages develop from fetal monocytes that differentiate into long-lived cells in the first week of life via GM-CSF. The Journal of experimental medicine 210, 1977–1992

18. Yu, X., Buttgereit, A., Lelios, I., Utz, S. G., Cansever, D., Becher, B., and Greter, M. (2017) The Cytokine TGF-beta Promotes the Development and Homeostasis of Alveolar Macrophages. Immunity 47, 903–912 e904

19. Schneider, C., Nobs, S. P., Kurrer, M., Rehrauer, H., Thiele, C., and Kopf, M. (2014) Induction of the nuclear receptor PPAR-gamma by the cytokine GM-CSF is critical for the differentiation of fetal monocytes into alveolar macrophages. Nature immunology 15, 1026–1037

20. Woods, P. S., Kimmig, L. M., Meliton, A. Y., Sun, K. A., Tian, Y., O’Leary, E. M., Gokalp, G. A., Hamanaka, R. B., and Mutlu, G. M. (2020) Tissue-Resident Alveolar Macrophages Do Not Rely on Glycolysis for LPS-induced Inflammation. Am J Respir Cell Mol Biol 62, 243–255

21. Pereverzeva, L., van Linge, C. C. A., Schuurman, A. R., Klarenbeek, A. M., Ramirez Moral, I., Otto, N. A., Peters-Sengers, H., Butler, J. M., Schomakers, B. V., van Weeghel, M., Houtkooper, R. H., Wiersinga, W. J., Bonta, P. I., Annema, J. T., de Vos, A. F., and van der Poll, T. (2022) Human alveolar macrophages do not rely on glucose metabolism upon activation by lipopolysaccharide. Biochim Biophys Acta Mol Basis Dis 1868, 166488

22. Shibata, Y., Berclaz, P. Y., Chroneos, Z. C., Yoshida, M., Whitsett, J. A., and Trapnell, B. C. (2001) GM-CSF regulates alveolar macrophage differentiation and innate immunity in the lung through PU.1. Immunity 15, 557–567

23. Trapnell, B. C., and Whitsett, J. A. (2002) Gm-CSF regulates pulmonary surfactant homeostasis and alveolar macrophage-mediated innate host defense. Annu Rev Physiol 64, 775–802

24. Bain, C. C., and MacDonald, A. S. (2022) The impact of the lung environment on macrophage development, activation and function: diversity in the face of adversity. Mucosal Immunol 15, 223–234

25. Vogel, D. Y., Vereyken, E. J., Glim, J. E., Heijnen, P. D., Moeton, M., van der Valk, P., Amor, S., Teunissen, C. E., van Horssen, J., and Dijkstra, C. D. (2013) Macrophages in inflammatory multiple sclerosis lesions have an intermediate activation status. J Neuroinflammation 10, 35

26. Osterholzer, J. J., Chen, G. H., Olszewski, M. A., Zhang, Y. M., Curtis, J. L., Huffnagle, G. B., and Toews, G. B. (2011) Chemokine receptor 2-mediated accumulation of fungicidal exudate macrophages in mice that clear cryptococcal lung infection. The American journal of pathology 178, 198–211

27. Thiel, B. A., Lundberg, K. C., Schlatzer, D., Jarvela, J., Li, Q., Shaw, R., Reba, S. M., Fletcher, S., Beckloff, S. E., Chance, M. R., Boom, W. H., Silver, R. F., and Bebek, G. (2024) Human alveolar macrophages display marked hypo-responsiveness to IFN-gamma in both proteomic and gene expression analysis. PloS one 19, e0295312

28. Busch, C. J., Favret, J., Geirsdottir, L., Molawi, K., and Sieweke, M. H. (2019) Isolation and Long-term Cultivation of Mouse Alveolar Macrophages. Bio Protoc 9

29. Thomas, S. T., Wierenga, K. A., Pestka, J. J., and Olive, A. J. (2022) Fetal Liver-Derived Alveolar-like Macrophages: A Self-Replicating Ex Vivo Model of Alveolar Macrophages for Functional Genetic Studies. Immunohorizons 6, 156–169

30. Heng, T. S., Painter, M. W., and Immunological Genome Project, C. (2008) The Immunological Genome Project: networks of gene expression in immune cells. Nature immunology 9, 1091–1094

31. Subramanian, S., Busch, C. J., Molawi, K., Geirsdottir, L., Maurizio, J., Vargas Aguilar, S., Belahbib, H., Gimenez, G., Yuda, R. A. A., Burkon, M., Favret, J., Gholamhosseinian Najjar, S., de Laval, B., Kandalla, P. K., Sarrazin, S., Alexopoulou, L., and Sieweke, M. H. (2022) Long-term culture-expanded alveolar macrophages restore their full epigenetic identity after transfer in vivo. Nature immunology 23, 458–468

32. Hardison, S. E., Herrera, G., Young, M. L., Hole, C. R., Wozniak, K. L., and Wormley, F. L., Jr. (2012) Protective immunity against pulmonary cryptococcosis is associated with STAT1-mediated classical macrophage activation. Journal of immunology 189, 4060–4068

33. Hardison, S. E., Ravi, S., Wozniak, K. L., Young, M. L., Olszewski, M. A., and Wormley, F. L., Jr. (2010) Pulmonary infection with an interferon-gamma-producing Cryptococcus neoformans strain results in classical macrophage activation and protection. The American journal of pathology 176, 774–785

34. Leopold Wager, C. M., Hole, C. R., Wozniak, K. L., Olszewski, M. A., Mueller, M., and Wormley, F. L., Jr. (2015) STAT1 signaling within macrophages is required for antifungal activity against Cryptococcus neoformans. Infection and immunity 83, 4513–4527

35. Wong, L. H., Sim, H., Chatterjee-Kishore, M., Hatzinisiriou, I., Devenish, R. J., Stark, G., and Ralph, S. J. (2002) Isolation and characterization of a human STAT1 gene regulatory element. Inducibility by interferon (IFN) types I and II and role of IFN regulatory factor-1. The Journal of biological chemistry 277, 19408–19417

36. Kershaw, N. J., Murphy, J. M., Liau, N. P., Varghese, L. N., Laktyushin, A., Whitlock, E. L., Lucet, I. S., Nicola, N. A., and Babon, J. J. (2013) SOCS3 binds specific receptor-JAK complexes to control cytokine signaling by direct kinase inhibition. Nat Struct Mol Biol 20, 469–476

37. Feldmesser, M., Kress, Y., Novikoff, P., and Casadevall, A. (2000) Cryptococcus neoformans is a facultative intracellular pathogen in murine pulmonary infection. Infection and immunity 68, 4225–4237

38. Coelho, C., Bocca, A. L., and Casadevall, A. (2014) The intracellular life of Cryptococcus neoformans. Annual review of pathology 9, 219–238

39. Kechichian, T. B., Shea, J., and Del Poeta, M. (2007) Depletion of alveolar macrophages decreases the dissemination of a glucosylceramide-deficient mutant of Cryptococcus neoformans in immunodeficient mice. Infection and immunity 75, 4792–4798

40. Santiago-Tirado, F. H., and Doering, T. L. (2017) False friends: Phagocytes as Trojan horses in microbial brain infections. PLoS pathogens 13, e1006680

41. Santiago-Tirado, F. H., Onken, M. D., Cooper, J. A., Klein, R. S., and Doering, T. L. (2017) Trojan Horse Transit Contributes to Blood-Brain Barrier Crossing of a Eukaryotic Pathogen. mBio 8

42. Alvarez, M., and Casadevall, A. (2006) Phagosome extrusion and host-cell survival after Cryptococcus neoformans phagocytosis by macrophages. Current biology: CB 16, 2161–2165

43. Ma, H., Croudace, J. E., Lammas, D. A., and May, R. C. (2006) Expulsion of live pathogenic yeast by macrophages. Current biology: CB 16, 2156–2160

44. Alvarez, M., and Casadevall, A. (2007) Cell-to-cell spread and massive vacuole formation after Cryptococcus neoformans infection of murine macrophages. BMC Immunol 8, 16

45. Ma, H., Croudace, J. E., Lammas, D. A., and May, R. C. (2007) Direct cell-to-cell spread of a pathogenic yeast. BMC Immunol 8, 15

46. Dragotakes, Q., Fu, M. S., and Casadevall, A. (2019) Dragotcytosis: Elucidation of the Mechanism for Cryptococcus neoformans Macrophage-to-Macrophage Transfer. Journal of immunology 202, 2661–2670

47. Corcoran, S. E., and O’Neill, L. A. (2016) HIF1alpha and metabolic reprogramming in inflammation. J Clin Invest 126, 3699–3707

48. Naito, Y., Takagi, T., and Higashimura, Y. (2014) Heme oxygenase-1 and anti-inflammatory M2 macrophages. Arch Biochem Biophys 564, 83–88

49. Michelucci, A., Cordes, T., Ghelfi, J., Pailot, A., Reiling, N., Goldmann, O., Binz, T., Wegner, A., Tallam, A., Rausell, A., Buttini, M., Linster, C. L., Medina, E., Balling, R., and Hiller, K. (2013) Immune-responsive gene 1 protein links metabolism to immunity by catalyzing itaconic acid production. Proceedings of the National Academy of Sciences of the United States of America 110, 7820–7825

50. Shi, S., Blumenthal, A., Hickey, C. M., Gandotra, S., Levy, D., and Ehrt, S. (2005) Expression of many immunologically important genes in Mycobacterium tuberculosis-infected macrophages is independent of both TLR2 and TLR4 but dependent on IFN-alphabeta receptor and STAT1. Journal of immunology 175, 3318–3328

51. Naujoks, J., Tabeling, C., Dill, B. D., Hoffmann, C., Brown, A. S., Kunze, M., Kempa, S., Peter, A., Mollenkopf, H. J., Dorhoi, A., Kershaw, O., Gruber, A. D., Sander, L. E., Witzenrath, M., Herold, S., Nerlich, A., Hocke, A. C., van Driel, I., Suttorp, N., Bedoui, S., Hilbi, H., Trost, M., and Opitz, B. (2016) IFNs Modify the Proteome of Legionella-Containing Vacuoles and Restrict Infection Via IRG1-Derived Itaconic Acid. PLoS pathogens 12, e1005408

52. Chen, H. R., Sun, Y., Mittler, G., Rumpf, T., Shvedunova, M., Grosschedl, R., and Akhtar, A. (2024) MOF-mediated PRDX1 acetylation regulates inflammatory macrophage activation. Cell Rep 43, 114682

53. Chu, W. T., Chu, X., and Wang, J. (2020) Investigations of the underlying mechanisms of HIF-1alpha and CITED2 binding to TAZ1. Proceedings of the National Academy of Sciences of the United States of America 117, 5595–5603

54. Freedman, S. J., Sun, Z. Y., Kung, A. L., France, D. S., Wagner, G., and Eck, M. J. (2003) Structural basis for negative regulation of hypoxia-inducible factor-1alpha by CITED2. Nat Struct Biol 10, 504–512

55. Lou, X., Sun, S., Chen, W., Zhou, Y., Huang, Y., Liu, X., Shan, Y., and Wang, C. (2011) Negative feedback regulation of NF-kappaB action by CITED2 in the nucleus. Journal of immunology 186, 539–548

56. Pong Ng, H., Kim, G. D., Ricky Chan, E., Dunwoodie, S. L., and Mahabeleshwar, G. H. (2020) CITED2 limits pathogenic inflammatory gene programs in myeloid cells. FASEB J 34, 12100–12113

57. Zafar, A., Ng, H. P., Chan, E. R., Dunwoodie, S. L., and Mahabeleshwar, G. H. (2023) Myeloid-CITED2 Deficiency Exacerbates Diet-Induced Obesity and Pro-Inflammatory Macrophage Response. Cells 12

58. Zafar, A., Pong Ng, H., Diamond-Zaluski, R., Kim, G. D., Ricky Chan, E., Dunwoodie, S. L., Smith, J. D., and Mahabeleshwar, G. H. (2021) CITED2 inhibits STAT1-IRF1 signaling and atherogenesis. FASEB J 35, e21833

59. Subramani, A., Hite, M. E. L., Garcia, S., Maxwell, J., Kondee, H., Millican, G. E., McClelland, E. E., Seipelt-Thiemann, R. L., and Nelson, D. E. (2023) Regulation of macrophage IFNgamma-stimulated gene expression by the transcriptional coregulator CITED1. Journal of cell science 136

60. Hu, R., Dai, A., and Tan, S. (2002) Hypoxia-inducible factor 1 alpha upregulates the expression of inducible nitric oxide synthase gene in pulmonary arteries of hyposic rat. Chin Med J (Engl) 115, 1833–1837

61. Jung, F., Palmer, L. A., Zhou, N., and Johns, R. A. (2000) Hypoxic regulation of inducible nitric oxide synthase via hypoxia inducible factor-1 in cardiac myocytes. Circ Res 86, 319–325

62. Melillo, G., Taylor, L. S., Brooks, A., Musso, T., Cox, G. W., and Varesio, L. (1997) Functional requirement of the hypoxia-responsive element in the activation of the inducible nitric oxide synthase promoter by the iron chelator desferrioxamine. The Journal of biological chemistry 272, 12236–12243

63. Peyssonnaux, C., Datta, V., Cramer, T., Doedens, A., Theodorakis, E. A., Gallo, R. L., Hurtado-Ziola, N., Nizet, V., and Johnson, R. S. (2005) HIF-1alpha expression regulates the bactericidal capacity of phagocytes. J Clin Invest 115, 1806–1815

64. Monga, D. P. (1981) Role of macrophages in resistance of mice to experimental cryptococcosis. Infection and immunity 32, 975–978

65. Mukherjee, S., Feldmesser, M., and Casadevall, A. (1996) J774 murine macrophage-like cell interactions with Cryptococcus neoformans in the presence and absence of opsonins. The Journal of infectious diseases 173, 1222–1231

66. Lee, H. H., Del Pozzo, J., Salamanca, S. A., Hernandez, H., and Martinez, L. R. (2019) Reduced phagocytosis and killing of Cryptococcus neoformans biofilm-derived cells by J774.16 macrophages is associated with fungal capsular production and surface modification. Fungal Genet Biol 132, 103258

67. Alanio, A., Desnos-Ollivier, M., and Dromer, F. (2011) Dynamics of Cryptococcus neoformans-macrophage interactions reveal that fungal background influences outcome during cryptococcal meningoencephalitis in humans. mBio 2

68. Lafont, E., Sturny-Leclere, A., Coelho, C., Lanternier, F., and Alanio, A. (2024) Assessing Phagocytosis of Cryptococcus neoformans Cells in Human Monocytes or the J774 Murine Macrophage Cell Line. Methods in molecular biology 2775, 157–169

69. Chen, G. H., McDonald, R. A., Wells, J. C., Huffnagle, G. B., Lukacs, N. W., and Toews, G. B. (2005) The gamma interferon receptor is required for the protective pulmonary inflammatory response to Cryptococcus neoformans. Infection and immunity 73, 1788–1796

70. Malur, A., McCoy, A. J., Arce, S., Barna, B. P., Kavuru, M. S., Malur, A. G., and Thomassen, M. J. (2009) Deletion of PPAR gamma in alveolar macrophages is associated with a Th-1 pulmonary inflammatory response. Journal of immunology 182, 5816–5822

71. Xaus, J., Cardo, M., Valledor, A. F., Soler, C., Lloberas, J., and Celada, A. (1999) Interferon gamma induces the expression of p21waf-1 and arrests macrophage cell cycle, preventing induction of apoptosis. Immunity 11, 103–113

72. Huang, S., Zhu, B., Cheon, I. S., Goplen, N. P., Jiang, L., Zhang, R., Peebles, R. S., Mack, M., Kaplan, M. H., Limper, A. H., and Sun, J. (2019) PPAR-gamma in Macrophages Limits Pulmonary Inflammation and Promotes Host Recovery following Respiratory Viral Infection. J Virol 93

73. Zhang, K., Sun, L., and Kang, Y. (2023) Regulation of phosphoglycerate kinase 1 and its critical role in cancer. Cell Commun Signal 21, 240

74. Chen, Z., He, Q., Lu, T., Wu, J., Shi, G., He, L., Zong, H., Liu, B., and Zhu, P. (2023) mcPGK1-dependent mitochondrial import of PGK1 promotes metabolic reprogramming and self-renewal of liver TICs. Nature communications 14, 1121

75. Li, X., Jiang, Y., Meisenhelder, J., Yang, W., Hawke, D. H., Zheng, Y., Xia, Y., Aldape, K., He, J., Hunter, T., Wang, L., and Lu, Z. (2016) Mitochondria-Translocated PGK1 Functions as a Protein Kinase to Coordinate Glycolysis and the TCA Cycle in Tumorigenesis. Molecular cell 61, 705–719

76. Wang, T., Liu, H., Lian, G., Zhang, S. Y., Wang, X., and Jiang, C. (2017) HIF1alpha-Induced Glycolysis Metabolism Is Essential to the Activation of Inflammatory Macrophages. Mediators Inflamm 2017, 9029327

77. Kong, D., Park, E. J., Stephen, A. G., Calvani, M., Cardellina, J. H., Monks, A., Fisher, R. J., Shoemaker, R. H., and Melillo, G. (2005) Echinomycin, a small-molecule inhibitor of hypoxia-inducible factor-1 DNA-binding activity. Cancer Res 65, 9047–9055

78. Svedberg, F. R., Brown, S. L., Krauss, M. Z., Campbell, L., Sharpe, C., Clausen, M., Howell, G. J., Clark, H., Madsen, J., Evans, C. M., Sutherland, T. E., Ivens, A. C., Thornton, D. J., Grencis, R. K., Hussell, T., Cunoosamy, D. M., Cook, P. C., and MacDonald, A. S. (2019) The lung environment controls alveolar macrophage metabolism and responsiveness in type 2 inflammation. Nature immunology 20, 571–580

79. Guth, A. M., Janssen, W. J., Bosio, C. M., Crouch, E. C., Henson, P. M., and Dow, S. W. (2009) Lung environment determines unique phenotype of alveolar macrophages. Am J Physiol Lung Cell Mol Physiol 296, L936–946

80. Woods, P. S., Kimmig, L. M., Sun, K. A., Meliton, A. Y., Shamaa, O. R., Tian, Y., Cetin-Atalay, R., Sharp, W. W., Hamanaka, R. B., and Mutlu, G. M. (2022) HIF-1alpha induces glycolytic reprograming in tissue-resident alveolar macrophages to promote cell survival during acute lung injury. Elife 11

81. Rosa, R. L., Berger, M., Santi, L., Driemeier, D., Barros Terraciano, P., Campos, A. R., Guimaraes, J. A., Vainstein, M. H., Yates, J. R., 3rd, and Beys-da-Silva, W. O. (2019) Proteomics of Rat Lungs Infected by Cryptococcus gattii Reveals a Potential Warburg-like Effect. J Proteome Res 18, 3885–3895

82. Guo, C., Islam, R., Zhang, S., and Fang, J. (2021) Metabolic reprogramming of macrophages and its involvement in inflammatory diseases. EXCLI J 20, 628–641

83. Olson, N., and van der Vliet, A. (2011) Interactions between nitric oxide and hypoxia-inducible factor signaling pathways in inflammatory disease. Nitric Oxide 25, 125–137

84. Metzen, E., Zhou, J., Jelkmann, W., Fandrey, J., and Brune, B. (2003) Nitric oxide impairs normoxic degradation of HIF-1alpha by inhibition of prolyl hydroxylases. Mol Biol Cell 14, 3470–3481

85. Rius, J., Guma, M., Schachtrup, C., Akassoglou, K., Zinkernagel, A. S., Nizet, V., Johnson, R. S., Haddad, G. G., and Karin, M. (2008) NF-kappaB links innate immunity to the hypoxic response through transcriptional regulation of HIF-1alpha. Nature 453, 807–811

86. O’Neill, L. A. J., and Artyomov, M. N. (2019) Itaconate: the poster child of metabolic reprogramming in macrophage function. Nat Rev Immunol 19, 273–281

87. Kim, G. D., Das, R., Rao, X., Zhong, J., Deiuliis, J. A., Ramirez-Bergeron, D. L., Rajagopalan, S., and Mahabeleshwar, G. H. (2018) CITED2 Restrains Proinflammatory Macrophage Activation and Response. Molecular and cellular biology 38

88. Qin, X., Chen, H., Tu, L., Ma, Y., Liu, N., Zhang, H., Li, D., Riedl, B., Bierer, D., Yin, F., and Li, Z. (2021) Potent Inhibition of HIF1alpha and p300 Interaction by a Constrained Peptide Derived from CITED2. J Med Chem 64, 13693–13703

89. Dobin, A., Davis, C. A., Schlesinger, F., Drenkow, J., Zaleski, C., Jha, S., Batut, P., Chaisson, M., and Gingeras, T. R. (2013) STAR: ultrafast universal RNA-seq aligner. Bioinformatics 29, 15–21

90. Cunningham, F., Achuthan, P., Akanni, W., Allen, J., Amode, M. R., Armean, I. M., Bennett, R., Bhai, J., Billis, K., Boddu, S., Cummins, C., Davidson, C., Dodiya, K. J., Gall, A., Giron, C. G., Gil, L., Grego, T., Haggerty, L., Haskell, E., Hourlier, T., Izuogu, O. G., Janacek, S. H., Juettemann, T., Kay, M., Laird, M. R., Lavidas, I., Liu, Z., Loveland, J. E., Marugan, J. C., Maurel, T., McMahon, A. C., Moore, B., Morales, J., Mudge, J. M., Nuhn, M., Ogeh, D., Parker, A., Parton, A., Patricio, M., Abdul Salam, A. I., Schmitt, B. M., Schuilenburg, H., Sheppard, D., Sparrow, H., Stapleton, E., Szuba, M., Taylor, K., Threadgold, G., Thormann, A., Vullo, A., Walts, B., Winterbottom, A., Zadissa, A., Chakiachvili, M., Frankish, A., Hunt, S. E., Kostadima, M., Langridge, N., Martin, F. J., Muffato, M., Perry, E., Ruffier, M., Staines, D. M., Trevanion, S. J., Aken, B. L., Yates, A. D., Zerbino, D. R., and Flicek, P. (2019) Ensembl 2019. Nucleic Acids Res 47, D745–D751

91. Merchant, N., Lyons, E., Goff, S., Vaughn, M., Ware, D., Micklos, D., and Antin, P. (2016) The iPlant Collaborative: Cyberinfrastructure for Enabling Data to Discovery for the Life Sciences. PLoS Biol 14, e1002342

92. Gruening, B. A. (2016) galaxytools. 07.2016 Ed., Zenodo

93. Robinson, M. D., McCarthy, D. J., and Smyth, G. K. (2010) edgeR: a Bioconductor package for differential expression analysis of digital gene expression data. Bioinformatics 26, 139–140

94. Loraine, A. E., Blakley, I. C., Jagadeesan, S., Harper, J., Miller, G., and Firon, N. (2015) Analysis and visualization of RNA-Seq expression data using RStudio, Bioconductor, and Integrated Genome Browser. Methods Mol Biol 1284, 481–501

95. Trapnell, C., Hendrickson, D. G., Sauvageau, M., Goff, L., Rinn, J. L., and Pachter, L. (2013) Differential analysis of gene regulation at transcript resolution with RNA-seq. Nat Biotechnol 31, 46–53

96. Huang da, W., Sherman, B. T., and Lempicki, R. A. (2009) Systematic and integrative analysis of large gene lists using DAVID bioinformatics resources. Nat Protoc 4, 44–57

97. Huang da, W., Sherman, B. T., and Lempicki, R. A. (2009) Bioinformatics enrichment tools: paths toward the comprehensive functional analysis of large gene lists. Nucleic Acids Res 37, 1–13

98. Subramanian, A., Tamayo, P., Mootha, V. K., Mukherjee, S., Ebert, B. L., Gillette, M. A., Paulovich, A., Pomeroy, S. L., Golub, T. R., Lander, E. S., and Mesirov, J. P. (2005) Gene set enrichment analysis: a knowledge-based approach for interpreting genome-wide expression profiles. Proceedings of the National Academy of Sciences of the United States of America 102, 15545–15550

99. Mootha, V. K., Lindgren, C. M., Eriksson, K. F., Subramanian, A., Sihag, S., Lehar, J., Puigserver, P., Carlsson, E., Ridderstrale, M., Laurila, E., Houstis, N., Daly, M. J., Patterson, N., Mesirov, J. P., Golub, T. R., Tamayo, P., Spiegelman, B., Lander, E. S., Hirschhorn, J. N., Altshuler, D., and Groop, L. C. (2003) PGC-1alpha-responsive genes involved in oxidative phosphorylation are coordinately downregulated in human diabetes. Nat Genet 34, 267–273

100. Varet, H., Brillet-Gueguen, L., Coppee, J. Y., and Dillies, M. A. (2016) SARTools: A DESeq2– and EdgeR-Based R Pipeline for Comprehensive Differential Analysis of RNA-Seq Data. PloS one 11, e0157022

101. Gautier, E. L., Shay, T., Miller, J., Greter, M., Jakubzick, C., Ivanov, S., Helft, J., Chow, A., Elpek, K. G., Gordonov, S., Mazloom, A. R., Ma’ayan, A., Chua, W. J., Hansen, T. H., Turley, S. J., Merad, M., Randolph, G. J., and Immunological Genome, C. (2012) Gene-expression profiles and transcriptional regulatory pathways that underlie the identity and diversity of mouse tissue macrophages. Nature immunology 13, 1118–1128

102. Baldarelli, R. M., Smith, C. L., Ringwald, M., Richardson, J. E., Bult, C. J., and Mouse Genome Informatics, G. (2024) Mouse Genome Informatics: an integrated knowledgebase system for the laboratory mouse. Genetics 227

103. Blake, J. A., Baldarelli, R., Kadin, J. A., Richardson, J. E., Smith, C. L., Bult, C. J., and Mouse Genome Database, G. (2021) Mouse Genome Database (MGD): Knowledgebase for mouse-human comparative biology. Nucleic Acids Res 49, D981–D987

104. Ankley, L. M., Conner, K. N., Vielma, T. E., Godfrey, J. J., Thapa, M., and Olive, A. J. (2024) GSK3alpha/beta Restrain IFN-gamma-Inducible Costimulatory Molecule Expression in Alveolar Macrophages, Limiting CD4+ T Cell Activation. Immunohorizons 8, 147–162

105. Schindelin, J., Arganda-Carreras, I., Frise, E., Kaynig, V., Longair, M., Pietzsch, T., Preibisch, S., Rueden, C., Saalfeld, S., Schmid, B., Tinevez, J. Y., White, D. J., Hartenstein, V., Eliceiri, K., Tomancak, P., and Cardona, A. (2012) Fiji: an open-source platform for biological-image analysis. Nat Methods 9, 676–682

106. Davis, A. E. (2025) Modeling Asymmetric Cell Division of Pathogenic Yeast within Host Macrophages for Comparison to Fluorescence Confocal Microscopy Data. Middle Tennessee State University

107. Burnham, K. P., and Anderson, D. R. (2004) Multimodel Inference. Sociological Methods & Research 33, 261 – 304

