## Supplementary material for "Intracellular *C. neoformans* infection stimulates increased glycolytic activity in fetal liver-derived alveolar-like macrophages": Math modeling appendix

### 1 Modeling the probability of an empirical count

In order to be counted, a daughter cell must produce a visible cross-section that is distinguishable from its mother cell. Our model allows for two types of countability. Type I countability occurs when both the mother and daughter produce a visible cross-section in the same z-slice, ensuring they are counted as separate cells. Type II countability occurs when as moving up or down the z-slices, the mother cell moves out of view as the daughter cell comes into view. Since the cells are spherical in shape, the concavity of the cross-sectional radius as a function of the depth of the z-slice enables us to distinguish mothers and daughters in this type of countability.

Both types of countability are influenced by the probability a daughter with radius $R_1$ produces a cross section that is large enough to be seen. We take the minimal radius of a visible cross-section, $R_L$, equal to the lateral resolution of the microscope. Letting $h$ denote the depth of the $z$-slice into the daughter and $z = 2$ the width of the z-stack, we determine the range of $h$ values for which a visible cross section is produced:

$$R_1 - \sqrt{R_1^2 - R_L^2} < h < R_1 + \sqrt{R_1^2 - R_L^2}. \tag{1}$$

The probability a daughter of radius $R_1$ produces a visible slice, $P_v(R_1)$, is then the quotient of the width of the visible range and the width of the $z$-stack:

$$P_v(R_1) = \frac{2\sqrt{R_1^2 - R_L^2}}{z}, \tag{2}$$

provided the expression on the left is less than one. The threshold radius for 100% visibility is then

$$R_v = \sqrt{\left(\frac{z}{2}\right)^2 + R_L^2}. \tag{3}$$

For $R_1 > R_v$, the daughter always produces a visible cross section.

#### 1.1 Type one countability

In this section, we quantify type I countability, that is, the probability of obtaining a visible cross section of the daughter and mother cell in a single $z$-slice. The angle between the daughter and mother cell, $\theta$, impacts type I countability. By symmetry, we restrict our attention to $0 < \theta < \frac{\pi}{2}$ and consider three cases based on the position of the $z$-slice relative to the center of the cells. Since the derivation is similar in all three cases, we restrict our presentation to the case where the center of the daughter and mother cells are below the focal $z$-slice, as illustrated in Figure 1.

Letting $h_1$ denote the distance from the top of the daughter to the focal $z$-slice and $h_2$ denote the distance from the top of the mother to the focal $z$-slice, we use similar triangles to express $h_2$ as a function of $h_1$ (see Figure (1)), and derive the visible range of $h_1$ values for the mother and daughter cells.

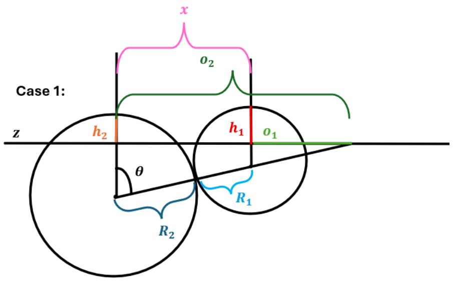


**Fig 1.** *Illustration of Derivation of Conditions for Type I Countability: The depth of the focal z slice (black line) into the mother cell ($h_2$) is expressed as a function of the depth of the slice into the daughter cell ($h_1$) using similar triangles.*

Letting $L_1 < h_1 < U_1$ denote the range of visibility for the daughter and $L_2 < h_1 < U_2$ denote the range of visibility for the mother, we find that

$$L_1 = R_1 - \sqrt{R_1^2 - R_L^2},$$
$$U_1 = R_1 + \sqrt{R_1^2 - R_L^2},$$
$$L_2 = R_1 + R_1 \cos\theta + R_2 \cos\theta - \sqrt{R_2^2 - R_L^2},$$
$$U_2 = R_1 + R_1 \cos\theta + R_2 \cos\theta + \sqrt{R_2^2 - R_L^2}.$$

Since $U_1 < U_2$, simultaneous visibility of both cells requires $L_2 < U_1$ and $L_2 < h_1 < U_1$ (see Figure 2). The condition $L_2 < U_1$ defines a critical angle, $\theta_1$, for simultaneous visibility of both cells:

$$L_2 < U_1 \text{ requires } \theta > \theta_1 \text{ where } \cos\theta_1 = \frac{\sqrt{R_2^2 - R_L^2} + \sqrt{R_1^2 - R_L^2}}{R_1 + R_2}.$$

Additionally, for $L_1 < L_2$, the range of simultaneous visibility of both cells reduces to the range of visibility for the daughter cell. The condition $L_1 < L_2$ defines a second critical angle $\theta_2$:

$$L_1 < L_2 \text{ requires } \theta < \theta_2 \text{ where } \cos\theta_2 = \frac{\sqrt{R_2^2 - R_L^2} - \sqrt{R_1^2 - R_L^2}}{R_1 + R_2}.$$

The critical angles $\theta_1$ and $\theta_2$ define three regions. In region 1, where $0 < \theta < \theta_1$, the ranges of visibility for mother and daughter are disjoint and the probability of simultaneously visualizing the mother and daughter is zero. In region 2, where $\theta_1 < \theta < \theta_2$, there exists partial overlap of the visibility ranges for the daughter and mother cell, and the range of type I countability is $L_2 < h_1 < U_1$. In region 3, where $\frac{\pi}{2} > \theta > \theta_2$, the visibility range of the daughter is contained in that of the mother so the range of type I visibility is $L_1 < h_1 < U_1$. Note that if $R_1 \geq R_v$, then $U_1 - L_1 > z$, so the daughter experiences 100% type I countability in region 3. Furthermore, if $R_1 \geq R_v$, then $U_1 - L_2 > z$, provided

$$\cos\theta \leq \frac{\sqrt{R_1^2 - R_L^2} + \sqrt{R_2^2 - R_L^2} - z}{R_1 + R_2}.$$

Hence, for $R_1 > R_v$ it is convenient to redefine the boundary $\theta_2$ by

$$\cos\theta_2 = \frac{\sqrt{R_1^2 - R_L^2} + \sqrt{R_2^2 - R_L^2} - z}{R_1 + R_2}.$$

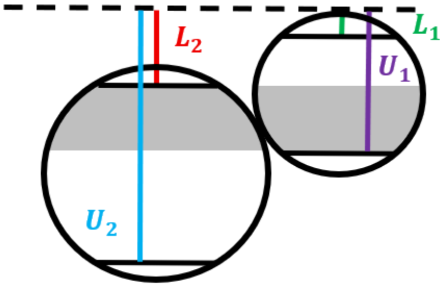


**Fig 2.** *Illustration of Range of Simultaneous Visibility of Mother and Daughter in Type I Countability: Here the depth of the $z$ slice, $h$, is measured relative to the top of the daughter cell. The range of visible depths for the mother is $L_2 < h < U_2$, while that is the daughter is $L_1 < h < U_1$. The overlap of the regions is seen in gray. $z$ slices within the gray region yield simultaneous visible cross sections of both cells enabling the daughter cell to be counted.*

With $\theta_2$ so redefined, for $R_1 \geq R_v$, a daughter experiences 100% type I countability in region 3, and less than 100% type I countability in region 2.

In summary, let

$$\cos\theta_2 = \begin{cases} \dfrac{\sqrt{R_1^2 - R_L^2} + \sqrt{R_2^2 - R_L^2} - z}{R_1 + R_2}, & R_v < R_1, \\ \dfrac{\sqrt{R_2^2 - R_L^2} - \sqrt{R_1^2 - R_L^2}}{R_1 + R_2}, & R_L < R_1 < R_v. \end{cases},$$

In case $0 < R < R_v$,

- The range of type I countability is empty in region 1 where $0 < \theta < \theta_1$.
- The range of type I countability is $L_2 < h_1 < U_1$ in region 2 where $\theta_1 < \theta < \theta_2$.
- The range of type I countability is $L_1 < h < U_1$ in region 3 where $\theta_2 < \theta < \frac{\pi}{2}$.

In case $R > R_v$,

- The range of type I countability is empty in region 1 where $0 < \theta < \theta_1$.
- The range of type I countability is $L_2 < h_1 < U_1$ in region 2 where $\theta_1 < \theta < \theta_2$.
- The width of the range of countability is greater than $z$ in region 3 where $\theta_2 < \theta < \frac{\pi}{2}$.

#### 1.2 Type two countability

In this section, we compute the probability a daughter of radius $R_1$ experiences type II countability. Recall type II countability occurs when the mother cell drops out of view as the daughter becomes visible when moving up or down the $z$-stack. With $\theta$, $h$, and $R_2$ as before, we formalize the conditions for type II countability:

1. The mother cell drops out of of view at depths less than $h$: $h < R_1 + (R_1 + R_2)\cos\theta - \sqrt{R_2^2 - R_L^2} = L_2$.
2. The cross section of the daughter at depth $h - z$ is larger than that at depth $h$: $h > l_2 := R_1 + \frac{1}{2}z$.

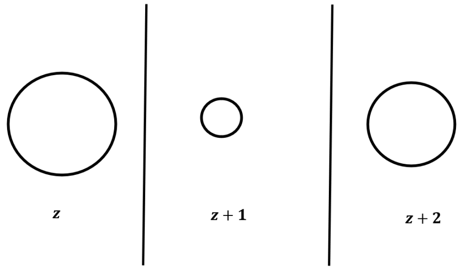


**Fig 3.** *Illustration of Type 2 Countability: Panels illustrate possible successive cross sections of a mother daughter pair. As we move through successive z slices, the mother cell drops out of view as the daughter comes into view. The concavity of the cell radius as a function of the depth of the z slice let's us distinguish daughter from mother in this case. This figure is for illustrative purposes only and is not drawn to scale.*

3. The cross section of the daughter at depth $h - z$ is visible:
   $l_3 := R_1 + z - \sqrt{R_1^2 - R_L^2} < h < u_3 := R_1 + z + \sqrt{R_1^2 - R_L^2}$.

While conditions 2 and 3 are always consistent, simultaneously satisfying conditions 1 and 2 requires

$$\cos\theta > \frac{\frac{1}{2}z + \sqrt{R_2^2 - R_L^2}}{R_1 + R_2}. \tag{4}$$

Similarly, simultaneously satisfying conditions 1 and 3 requires

$$\cos(\theta) > \frac{z + \sqrt{R_2^2 - R_L^2} - \sqrt{R_1^2 - R_L^2}}{R_1 + R_2}. \tag{5}$$

Conditions (4) and (5) place lower bounds on $R_1$ for Type II countability. The strongest bound comes from (5): Type II countability requires $R_1 > R_1^*$ where $R_1^*$ is defined by

$$R_1^* + \sqrt{(R_1^*)^2 - R_L^2} = z - R_2 + \sqrt{R_2^2 - R_L^2}.$$

Note $R_1^* < R_v$.

In addition, since $z = 2$, and $R_v^2 - R_L^2 = 1$, condition (5) is stronger for $R < R_v$ and condition (4) is stronger for $R > R_v$. Hence, we define

$$\cos\theta_{II} = \begin{cases} \dfrac{\frac{1}{2}z + \sqrt{R_2^2 - R_L^2}}{R_1 + R_2}, & R_v < R_1, \\ \dfrac{z + \sqrt{R_2^2 - R_L^2} - \sqrt{R_1^2 - R_L^2}}{R_1 + R_2}, & R_1^* < R_1 < R_v. \end{cases},$$

so that type II countability occurs for $\theta < \theta_{II}$.

We then derive the range of type II countability as a function of $R_1$. First consider the least upper bound of $h$ arising from conditions 1-2 above. Note $u_3 > L_2$ can be rearranged to

$$z > (R_1 + R_2)\cos(\theta) - \sqrt{R_1^2 - R_L^2} - \sqrt{R_2^2 - R_L^2}. \tag{6}$$

However, $f(x) := x - \sqrt{x^2 - R_L^2}$ is decreasing for $x > R_L$ and $f(R_L) = R_L < 1$. Hence, the right-hand side of (6) is less than $z = 2$, and the least upper bound of $h$ is $L_2$ in every case:

$$h < R_1 + (R_1 + R_2)\cos\theta - \sqrt{R_2^2 - R_L^2} = L_2.$$

Next we consider the greatest lower bound on $h$ arising from conditions 1-3.

- For $R_1 \geq R_v$ the greatest lower bound of $h$ comes from condition 2: $l_2 < h$.
- For $R_1 < R_v$ the greatest lower bound of $h$ comes from condition 3: $l_3 < h$.

In summary,

- For $R_1^* < R_1 < R_v$, the range of type II countability is $l_3 < h < L_2$.
- For $R_v < R_1$ the range of type II countability is

$$l_2 < h < L_2.$$

  In this case, the daughter experiences 100% type II countability for $0 < \theta < \theta_0$ where

$$\cos(\theta_0) := \frac{3 + \sqrt{R_2^2 - R_L^2}}{R_1 + R_2}.$$

##### 1.3 The probability a cell is countable

Combining the results from the previous two sections we compute the probability a daughter of radius $R_1$ and angle $\theta$ is countable as the quotient of the width of the range of countable $h$ values and the width of the $z$-stack. Fixing $R_1$ for the moment, let $p(\theta)$ be the probability a daughter of radius $R_1$ experiences type I or II countability as a function of the angle $\theta$ between the daughter and mother cell.

**Case A**: For $R_1 < R_1^* < R_v$ only type I countability occurs and the probability a daughter is countable varies with $\theta$ according to critical angles $0 < \theta_1 < \theta_2$:

- For $\theta < \theta_1$,

$$p(\theta) = 0.$$

- For $\theta_1 < \theta < \theta_2$,

$$p(\theta) = \frac{\sqrt{R_2^2 - R_L^2} + \sqrt{R_1^2 - R_L^2} - (R_1 + R_2)\cos\theta}{z}.$$

- For $\theta > \theta_2$,

$$p(\theta) = \frac{2\sqrt{R_1^2 - R_L^2}}{z}.$$

**Case B**: For $R_1^* < R_1 < R_v$ both type I and II countability occurs and the probability a daughter is countable varies with $\theta$ according to critical angles $0 < \theta_{II} < \theta_1 < \theta_2 \leq \frac{\pi}{2}$:

- For $0 \leq \theta \leq \theta_{II}$

$$p(\theta) = \frac{(R_1 + R_2)\cos(\theta) - \sqrt{R_2^2 - R_L^2} + \sqrt{R_1^2 - R_L^2} - 2}{z} < 1.$$

- For $\theta_{II} < \theta \leq \theta_1$

$$p(v|\theta) = 0.$$

- For $\theta_1 < \theta \leq \theta_2$,

$$p(\theta) = \frac{\sqrt{R_2^2 - R_L^2} + \sqrt{R_1^2 - R_L^2} - (R_1 + R_2)\cos(\theta)}{z} < 1,$$

  and for $\theta_2 < \theta \leq \frac{\pi}{2}$,

$$p(\theta) = \frac{2\sqrt{R_1^2 - R_L^2}}{z} < 1.$$

**Case C**: For $R_1 \geq R_v$ and $\sqrt{R_1^2 - R_L^2} \leq 3$ both type I and II countability occur. The probability a daughter is countable varies with $\theta$ according to the critical angles $0 < \theta_0 < \theta_1 \leq \theta_{II} < \theta_2 \leq \frac{\pi}{2}$:

- For $0 \leq \theta < \theta_0$,
$$p(\theta) = 1.$$
- For $\theta_0 \leq \theta < \theta_1$,
$$p(\theta) = \frac{(R_1 + R_2)\cos\theta - \sqrt{R_2^2 - R_L^2} - 1}{z} < 1.$$
- For $\theta_1 < \theta < \theta_{II}$, the type II countability range is
$$R_1 + 1 < h < R_1 + (R_1 + R_2)\cos\theta - \sqrt{R_2^2 - R_L^2}$$
and the type I countability range is
$$R_1 + (R_1 + R_2)\cos\theta - \sqrt{R_2^2 - R_L^2} < h < R_1 + \sqrt{R_1^2 - R_L^2},$$
giving us
$$p(\theta) = \frac{\sqrt{R_1^2 - R_L^2} - 1}{z} < 1.$$
For $\theta_{II} < \theta < \theta_2$
$$p(\theta) = \frac{\sqrt{R_2^2 - R_L^2} + \sqrt{R_1^2 - R_L^2} - (R_1 + R_2)\cos\theta}{z} < 1$$
For $\theta_2 \leq \theta < \frac{\pi}{2}$
$$p(\theta) = 1.$$

**Case D**: From the previous, we see that when $\sqrt{R_1^2 - R_L^2} > 3$, $\theta_2 < \theta_0$. The probability a daughter cell is countable is independent of $\theta$ and

$$p(\theta) = 1.$$

Assuming the angle between the daughter and mother cell is uniformly distributed, the probability a daughter of radius $R_1$ achieves type I or type II countability is

$$C(R_1) = \frac{2}{\pi}\int_0^{\frac{2}{\pi}} p(\theta)\, d\theta.$$

#### 1.4 The probability of a recorded count

In this section we describe how the probability of a count is determined from the probability an individual cell is countable and a model of error in the counting process. Since cell size increases monotonically with time, each cell age corresponds to a unique cell radius. Hence, the age index of a cell will serve as a proxy for its radius, and the probability a cell of radius $R_1$ is countable, $C(R_1)$, is replaced with the probability a cell of age index $j$ is countable, $C(j)$. Fixing an observation time, we first compute the probability $x$ cells of age class $j$ are countable given $N(j)$ cells are in this class. This probability mass function follows a binomial distribution.

$$P_j(x) = \binom{N(j)}{x}(C(j))^x(1 - C_{(j)})^{N(j)-x}$$

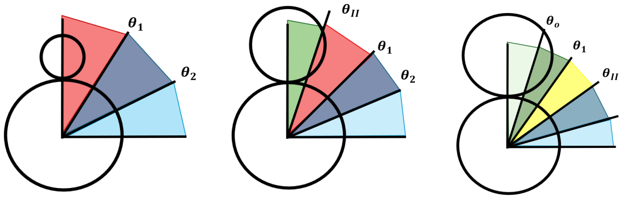


**Fig 4.** *Illustration of Countability Regions in Cases A-C. Panels depict countability cases A-C from left to right. The daughter cell's potential to be counted changes as she grows. In cases A and B, dark pink denotes regions where the small daughter is obscured by the mother. In case C, pale blue and pale green denote regions where the large daughter is always counted. This figure is for illustrative purposes only and is not drawn to scale.*

Next we compute the probability $j$ cells from the population are countable as the convolution of the probability mass functions for the number of counts from each age class. The convolution is computed sequentially in MATLAB, and initialized as the probability mass function for countability of the founding cell, $P_f$:

$$P(x) = (P_f * P_0 * P_1 * \dots P_J)(x),$$

where

$$P_f(x) = \binom{1}{x}(P_v(R_f))^x(1 - P_v(R_f))^{1-x},$$

$J$ denotes the maximal age class, and $(P_i * P_j)(x) = \sum_{y=0}^{x} P_i(x-y)P_j(y)$. Note, in contrast to daughter cells within the population, the founding cell's probability of being countable is equal to its probability of producing a visible cross-section.

Due to error, the actual count may not match the number of countable cells. We consider two types of counting error.

First, an observer might fail to notice a countable cell. We suppose each cell has a fixed probability $p_m$ of being missed. The probability of counting $x$ cells is then

$$PC(x) = \sum_{w=0}^{N} \binom{w}{x}(1 - p_m)^x(p_m)^{w-x}P(w), \tag{7}$$

where $N$ is the size of the population. The terms in equation 7 give the probability $x$ cells are counted and $w$ cells are countable, where it is understood that $\binom{w}{x} = 0$ for $w < x$.

Second, we allow for the possibility of artifacts which could lead to overestimation of the population size. The probability of observing an artifact is taken to be independent of the number of cells present, and very small, $r = 10^{-7}$. Taking artifacts into account, the probability of recording a count of $x$ cells is

$$PR(x) = PC(x)\frac{1-2r}{1-r} + PC(x-1)r + PC(x-2)r^2 + \cdots + PC(0)r^x. \tag{8}$$

In addition to representing two potential types of error in the recorded counts, the error models aid in convergence of the numerical routine by providing a means of comparing unlikely parameter choices. For example, if we neglect counting error, a parameter choice which includes one artifact and a parameter choice which includes two artifacts are equally likely, since both are impossible. Modeling unlikely events, provides information to help the solver recover from unlikely parameter choices without impacting comparisons between highly likely parameterizations.

### 2 Numerical methods for assessment of intracellular growth

#### 2.1 Overview

The model of population growth via asymmetric budding was fit to the count data by maximizing the likelihood of the parameters for the data: In brief, given a guess for the model parameters, $T_b$, $T_m$, and $a(0)$, where $a(0)$ denotes the initial age of the founding cell, the population model was solved using a time and age discretization of $\delta = 1$ min. See section (2.1.1) for details on the numerical implementation of the population growth model. The results were used to compute the probability of the empirical count at each observation time according to equation (8). Since host macrophages and intracellular yeast were highly mobile and the time between observations was relatively long (30 min), counts at distinct observation times were treated as independent conditional on the model parameters. Hence, the probability of the count data for a single cell or population was computed as the product of the probabilities at each observation time.

Rather than maximizing the probability of a count directly, we minimized its negative logarithm, i.e., the negative log-likelihood, so the best-fit parameters could be estimated using MATLAB fminsearch.m. For additional details on convergence see section 2.1.2.

When fitting individual cells, an initial round of optimization was followed by a comparison of best-fit parameters between cells using likelihood ratios: $\frac{\mathcal{L}(p_j|d_i)}{\mathcal{L}(p_i|d_i)}$, where $\mathcal{L}$ denotes the likelihood function, $p_i$ denotes the best-fit parameters for model $i$, and $d_i$ denotes the data for cell $i$. In so doing, $\mathcal{L}(p_j|d_i)$ was computed by optimizing the initial age of the founding cell $a_0$.

When possible, the results of model comparison between individual cells within a population were utilized to bootstrap improvements in the fit to each cell: Letting $N$ denote the number of cells in the host sample, for $i = 1, \dots, N$, in case a better fit to $d_i$ was found among the parameters $P = \{p_1, \dots, p_N\}$, the population growth model was refit to $d_i$ using the best-fit model in $P$ as an initial guess. This process was repeated until no further improvements in the likelihood of the fits could be made.

Some parameters and initial guesses used in the numerical simulations are listed below:

- $R_L = 0.22$ $\mu$m: radius of the smallest observable radius
- $z = 2$ $\mu$m: width of the z-stack
- $\lambda$: growth rate of a small daughter cell. $\lambda = 0.012$ $\mu$m min$^{-1}$ for J774 and $\lambda = 0.016$ $\mu$m min$^{-1}$ for FLAMs. See Materials and Methods.
- $a_0$: The initial age of a the founding cell. $a_0$ is determined by a scalar $m_0 \in [0, 1]$ according to $a_0 = m_0 a_M$, where $a_M$ stands for the maximal initial age (see section 2.1.1).
- When fitting individual cells, we used 4 random initial guesses for $m_0$.
- When fitting a population, we used used the following initial guesses for $m_0$:

$$m_0 = 0.9375, 0.4375, 0.1875, 0.0625$$

- $R_v = \frac{\sqrt{z^2 + R_L^2}}{2}$: the size of a cell that produces a visible cross section 100% of the time. $R_v = 1.0239$ $\mu$m for J774s and $FLAM$ cells.

- $R_c = \sqrt{9 + R_L^2}$: the radius at which a cell produces a countable image with probability one. $R_c = 3.0081$ $\mu$m for J774 cells and FLAMs.
- $a_v = \frac{R_v - R_L}{\lambda}$: the age corresponding to $R_v$. $a_v = 66.9923$ min for J774s and $a_v = 50.2446$ min for FLAMs.
- $a_c$: the age at which a cell produces a countable image with probability one. $a_c = 232.338$ min for J774s and $a_c = 174.2535$ min for FLAMs.
- $R^*$: the radius of a cell that is too small to experience Type II countability. $R_2$ is determined as the solution of the following equation $R^* + \sqrt{(R^*)^2 - R_L^2} = z - R_2 + \sqrt{R_2^2 - R_L^2}$, where $R_2$ is the radius of a typical mature cell (see Materials and Methods). $R^* = 1.0084$ $\mu$m for J774s and $R^* = 1.0088$ $\mu$m for FLAMs.
- Parameter bounds: $0 < T_b < T_m$, $T_m + T_b \leq 1600$, $a_c < T_m + T_b$
- Initial guesses for inter-budding and maturation times. $[T_b, T_m] = [180 \cdot \frac{4}{5}, 180]; [480 \cdot \frac{4}{5}, 480]; [600 \cdot \frac{4}{5}, 600]; [180 \cdot \frac{1}{3}, 180]; [480 \cdot \frac{1}{3}, 480]; [600 \cdot \frac{1}{3}, 600]; [180 \cdot \frac{1}{2}, 180]; [480 \cdot \frac{1}{2}, 480]; [600 \cdot \frac{1}{2}, 600]$:
- $max_{count} = 5$: count at which a times series of counts is truncated to minimize the impact of crowding.
- $min_{count} = 4$: when fitting individual cells, only include data for cells that achieve a count of at least 4 intracellular yeast.
- $R_2$: the average size of the mature intracellular yeast. $R_2 = 3.2$ $\mu$m for J774 and $R_2 = 3.6$ $\mu$m for FLAMs. See section Materials and Methods.

##### 2.1.1 Numerical Simulation of the Population Model

Recall the population growth model includes two dependent variables, $B(t, r)$ and $M(t, r, a)$, which represent the number immature daughter cells and mature mother cells, respectively, of reproductive age $r$ and, in the case of mothers, absolute age $a$, at time $t$. The absolute age of a mature mother cell is measured relative to its time in the mature mother stage class. The evolution equations:

$$
\begin{aligned}
B(t+\delta, r) &= B(t, r-\delta);\ 0 \leq t, ;\ \delta \leq r < T_m - \delta && (9)\\
M(t+\delta, r, a) &= M(t, r-\delta, a-\delta);\ 0 \leq t;\ \delta \leq r < T_m - \delta\ \ \delta \leq a < T_b - \delta && (10)
\end{aligned}
$$

are solved exactly in MATLAB, using a time discretization of $\delta = 1$ min.

The numerical algorithm tracks the number of mother and immature daughter cells of each age through time by following cells along the characteristic lines. This process is illustrated in Figures 5 and 6. For example, the characteristic lines for the mother cells have the form $(t(s), r(s), a(s)) = (t_0, r_0, a_0) + s(1, 1, 1)$.

The population model also includes the following boundary conditions:

$$
\begin{aligned}
B(t, 0) &= \mathbf{M}(t-\delta, T_b - \delta) + B(t-\delta, T_m - \delta);\ t > 0 && (11)\\
M(t, 0, a) &= M(t-\delta, T_b - \delta, a - \delta);\ t > 0, T_m < a && (12)\\
M(t, 0, 0) &= B(t-\delta, T_b - \delta);\ t > 0, T_m \leq a && (13)\\
B(t, T_m) &= 0;\ t \geq 0 && (14)\\
M(t, T_b, a) &= 0;\ t \geq 0 && (15)
\end{aligned}
$$

where $\mathbf{M}(t, r) = \sum_{a=0}^{\infty} M(t, r, a)$.

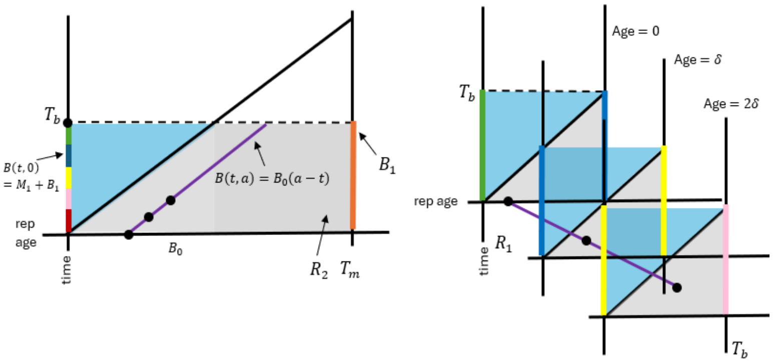


**Fig 5.** *Illustration of numerical method, part 1: The model is solved in the light gray regions $R_1$ and $R_2$ where the reproductive age, $r$, of cells is greater than time, $t$, by following the initial data only the characteristic lines. Sample characteristic lines are illustrated in purple. This procedure yields solution values along the boundaries $r = T_m$ for daughter cells (orange) and $r = T_b$ for mother cells (various colors) which are used to initialize the boundary $r = 0$ for both mother and daughter cells (various colors).*

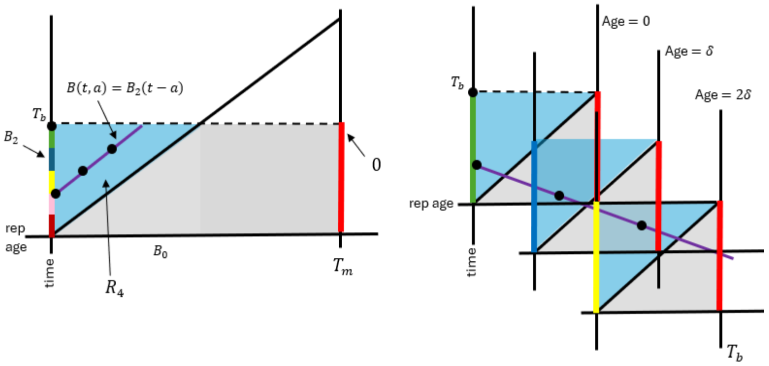


**Fig 6.** *Illustration of numerical method, part 2: To prevent double counting, cell counts on the boundaries $r = T_m$ and $r = T_b$ are zeroed out (red). The model is solved in the light blue regions $R_3$ and $R_4$ where $r < t$ by following the newly budded cells along the characteristic lines. Sample characteristic lines are illustrated in purple. This procedure yields solution values along the entire upper boundary $t = T_b$ (dashed black line) which are used to initialize the next iteration of the algorithm.*

To motivate equations 11-13, note that the final reproductive age class and initial reproductive age class are identified, as both represent the instant of budding. Hence the number of daughters and mothers of reproductive age zero at time $t$ is determined by the number of daughters and mothers of reproductive age $T_m - \delta$ and $T_b - \delta$, respectively, at time $t - \delta$. See equations 11-13. This transitional boundary condition is depicted in 5, by the color-coded boundaries. In addition, because the final and initial reproductive age class are identified, to avoid double counting, the boundaries $r = T_b$ (for mother cells) and $r = T_m$ (for daughters) are fixed to zero. The zeroing of the count for the terminal reproductive age is described in equations 14-15 and depicted in Figure 6 in red.

Our interest in the absolute age of mature mother cells stems from the fact that yeast cells grow as they age, and the size of a yeast cell impacts its probability of being counted. Since immature daughter cells have yet to bud, their reproductive age and absolute age are the same. Hence, reproductive age is sufficient to determine the size of a daughter cell. In contrast, mother cells undergo repeated cycles of budding, while their absolute age increases continuously with time. In the probabilistic counting model presented above, a cell is 100% countable, when it achieves a threshold size. The corresponding age is denoted as $a_C$. Since mothers of age greater than or equal to $a_C$ are 100% countable, there is no need to track a cell's age beyond $a > a_C$. Hence in the numerical implementation of the model, mother cells are absorbed into a final age class once achieving an age of $a_C$. These considerations give the condition for the absorbing final absolute age class in the numerical implementation of the model:

$$M(t+\delta, r, a_C) = M(t, r-\delta, a_C) + M(t, r-\delta, a_C-\delta) \tag{16}$$

This condition is depicted in the final panel of Figure 6.

Finally, we have the initial conditions:

$$B(0,r) = B_0(r);\ 0 < r < T_m \tag{17}$$

$$M(0,r,a) = M_0(r);\ 0 < r = a < T_b \tag{18}$$

Since we restrict our attention to cells with a count of one intracellular yeast at the first observation time, initializing the numerical routine only requires us to select the initial age of the founding cell, $a_0$. Since *Cryptococcus neoformans* always bud from the same location, a daughter must separate from its mother for a subsequent bud to form. For this reason we let inter-budding time, $T_b$, be the lower limit of the reproductive age for the initial cell. This choice ensures that the founding yeast cell is old enough to have separated from its mother and been taken up as a single cell by a macrophage. The upper bound for the initial reproductive age of the founder is set to $T_m + \min\{T_b, a_c\}$. If the founder's initial age is between $T_b$ and $T_m$, it is an immature daughter, and its reproductive age is equal to its absolute age. If the founder's age is greater than $T_m$, it's a mature mother cell, and hence, attached to a bud which was not counted at the initial observation. The constraint that the founder's initial age is less than $Tm + a_C$ ensures that its bud is not 100% countable. In case the founder is a mother, its initial reproductive age and that of its bud is determined by the remainder when $a_0 - Tm$ is divided by $Tb$.

An final constraint on the model parameters is $T_m > T_b$. This constraint is supported by research on other species of budding yeast [1] and visual inspection of our data.

##### 2.1.2 Error and Convergence Criteria

For evaluation of model fits and comparison of parameters between cells and populations we quantified the accuracy and error in the parameter estimates. Due to discretization, estimates of the parameters are only accurate to one minute. In addition to limits set by discretization, the convergence criteria must also be considered to determine a minimal significance level for differentiating the ability of parameters to describe data. When implementing fminseach.m (MATLAB) we used a tolerance of $Tol = 10^{-3}$. The criteria applied to the negative log likelihood for convergence is then [2]:

$$|y_{i+1} - y_i| \le Tol\,(1 + |y_i|), \tag{19}$$

where $y_i$ denotes the negative log-likelihood of parameter choice $i$. Exponentiating both sides yields a an inequality in terms of the likelihoods $P_i$:

$$e^{-Tol} P_i^{Tol} \le \frac{P_i}{P_{i+1}} \le e^{Tol} P_i^{-Tol}. \tag{20}$$

Thus, the relative error has an upper bound of $e^{Tol}P_i^{-Tol}$ and a lower bound of $e^{-Tol}P_i^{Tol}$. We see that smaller $P_i$ values result in larger relative error. In our data, the smallest likelihood observed at convergence for individual cells was $P_i = 0.0778$. Substituting this into the error bounds gives us,

$$e^{10^{-3}} \cdot (0.0778)^{10^{-3}} = 0.9984$$

and

$$e^{-10^{-3}} \cdot (0.0778)^{-10^{-3}} = 1.0016.$$

These calculations show that in the worst case scenario the relative accuracy of the likelihood values for the best-fit parameters are less than 1%. Hence, differences of 1% or more in relative likelihood are not the result of numerical error. However, there is one caveat to this conclusion, namely, that the solver is not guaranteed to converge to the global solution.

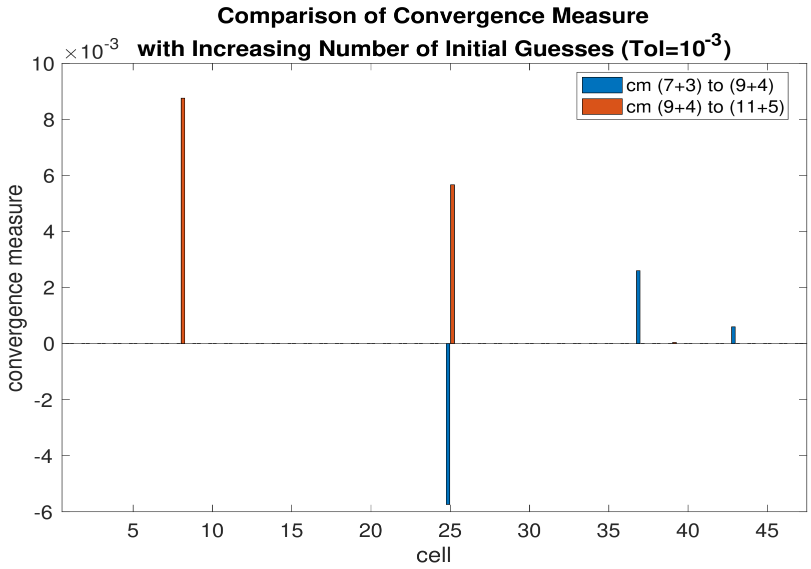


**Fig 7.** *Convergence Behavior with Increasing Initial Guesses: The likelihood values of numerical solutions to the parameter optimization routine for increasing numbers of initial guesses are compared using the convergence measure for the routine (see equation 19), where the tolerance for convergence is $10^{-3}$. The height of the bars gives the value of the convergence measure between solutions, where positive values indicate improvements in likelihood. The number of initial parameter guesses for the solution is represented as a sum $x + y$, where $x$ denotes the number of choices for the pair $(T_b, T_m)$ and $y$ denotes the number of choices for the initial age of a cell, $a_0$. Hence, each cell is fit using a total of $xy$ initial guesses for the parameters. A small number of cells ($n = 5$) exhibit low variability in the solution likelihood as the number of initial guesses increases (convergence measure $< 10^{-2}$).*

Figure 7 investigates convergence to a global optimum by comparing the convergence measure between solutions of runs with different numbers of initial guesses for the model parameters. If the convergence measure is less than the selected tolerance ($10^{-3}$), there is effectively no difference between the solutions in terms of likelihood. Looking at Figure (7) we see the convergence measure is very small when comparing solutions with different numbers of initial guesses, and beneath the convergence tolerance in all but a few cases. This provides confidence that the algorithm has converged to a solution that is very close to achieving the global optimum value.

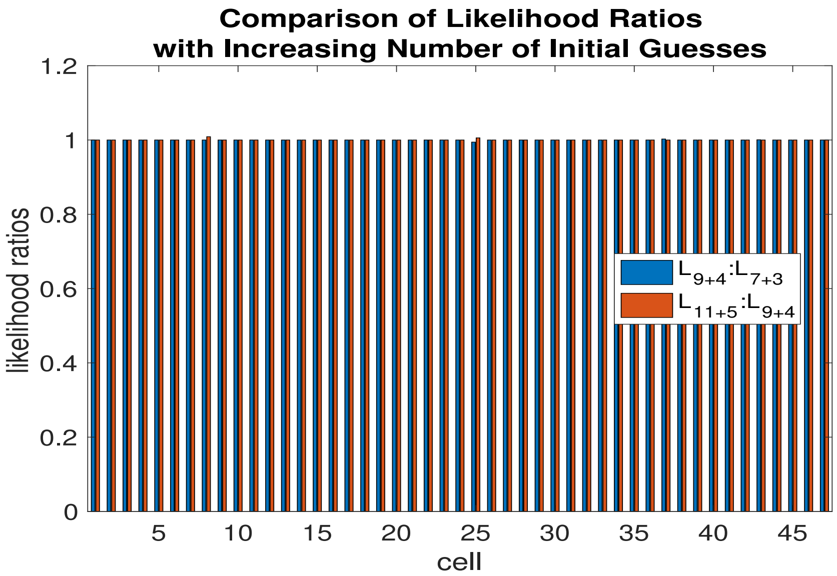


**Fig 8.** *Effect of Number of Initial Guesses on Likelihood Ratios: The likelihood values of numerical solutions to the parameter optimization routine for increasing numbers of initial guesses are compared using the likelihood ratios. Values greater than one indicate improvement in likelihood. The number of initial parameter guesses for the solution is represented as a sum $x + y$, where $x$ denotes the number of choices for the pair $(T_b, T_m)$ and $y$ denotes the number of choices for the initial age of a cell, $a_0$. Hence, each cell is fit using a total of $xy$ initial guesses for the parameters. A small number of cells exhibit low variability in the solution likelihood ($\sim 1\%$) as the number of initial guesses varies.*

Figure 8 shows the likelihood ratios between the solutions of runs with different numbers of initial guesses for the model parameters. In blue the figure displays the likelihood ratios for solutions found using nine parameters guesses and four guesses for initial age to seven parameter guesses and three guesses for initial age. In red the figure displays the likelihood ratio of solutions found using eleven parameter guesses and five guesses for initial age to nine parameter guesses and four guesses for initial age. All cells show little to minimal ($\sim 1\%$) change in the goodness of fit as we increase the number of guesses.

Figure 10 shows a scatter plot of parameter values for increasing numbers of initial guesses. As before each color represents a different set of initial guesses. The parameters remain in the same general region for all initial guesses tested. The green points, representing the largest number of initial guesses, often overlap or obscure the blue and red points, which represent smaller numbers of initial guesses. The fact that the estimated parameters are generally close together supports that the numerical simulations give a reasonably accurate estimation of the sample parameter distribution. However, we see that there is more uncertainty in the parameter estimates for individual cells than in the optimal likelihood values. This demonstrates that in some cases solutions with near identical likelihoods can have appreciable differences in their parameter values.

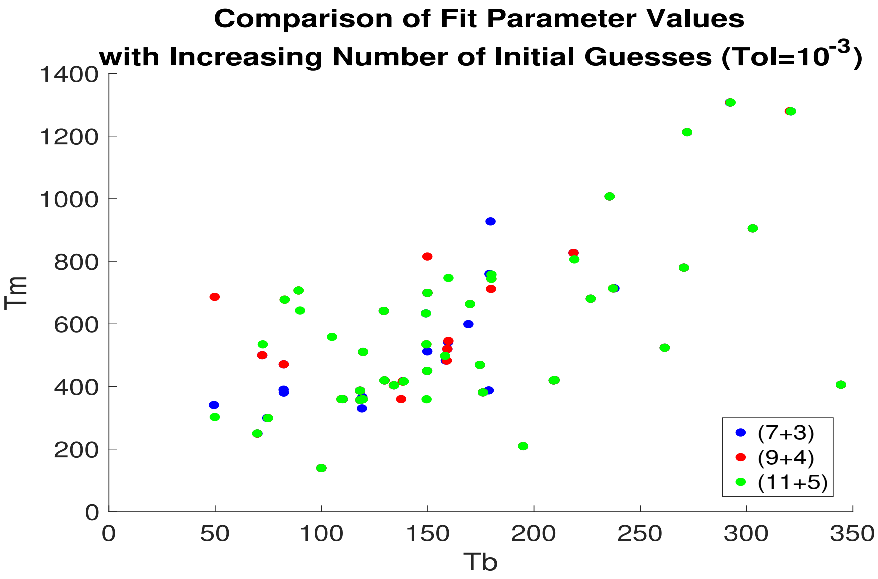


**Fig 9.** *Parameter Convergence Under Increasing Initial Guesses: The parameter values for numerical solutions to the parameter optimization routine for increasing numbers of initial guesses are compared. Visible red and blue dots indicate variability in the numerical solution's parameters. The number of initial parameter guesses for the solution is represented as a sum* $x + y$*, where* $x$ *denotes the number of choices for the pair* $(T_b, T_m)$ *and* $y$ *denotes the number of choices for the initial age of a cell,* $a_0$*. Hence, each cell is fit using a total of* $xy$ *initial guesses for the parameters.*

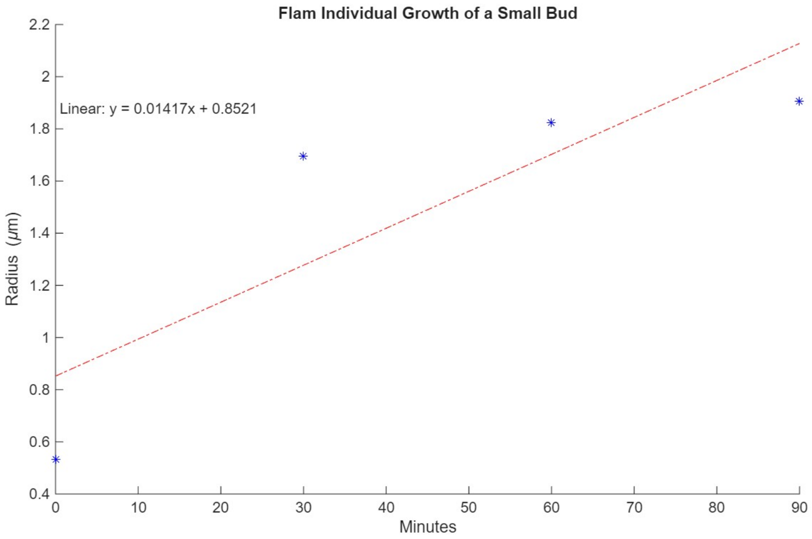


**Fig 10.** *Estimating the Growth Rate of Daughter Cells: Illustration of the method of estimating the linear growth of the daughter cell radius.*

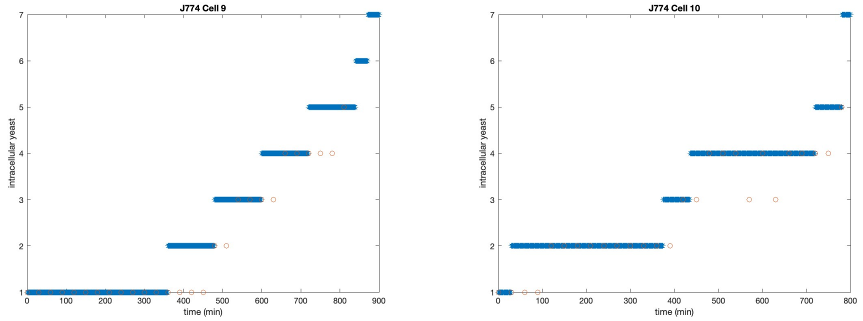


**Fig 11.** *Sample Fits of the Mathematical Model to Individual J774 Cells: Model population size (blue stars) are shown along with empirical counts (red circles). We see counts often underestimate the model population size, due in part to the limited visibility of small daughter cells. Right panel: Cell 10 demonstrates how the empirical count sometimes falls, presumably due to limited visibility of small daughters.*

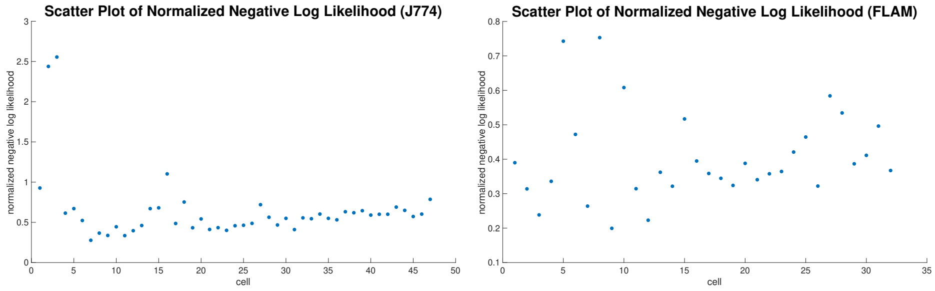


**Fig 12.** *Scatter Plots of Normalized Negative Log Likelihood Values for Numerical Fits of the Mathematical Model to Individual J774 and FLAM cells: Individual cell counts that include fewer than 2 complete inter-budding times, that is, fail to achieve a count of at least four cells, are excluded from the individual-cell analysis since they contain very limited information for estimating inter-budding and maturation times. Counts are truncated once they achieve a count of six to minimize miscounts due to cell crowding. Cells are arranged by the length of their count data in descending order.*

### 3 Supplementary Mathematical Figures

### References


1. Talia SD, Skotheim JM, Bean JM, Siggia ED, Cross FR. The effects of molecular noise and size control on variability in the budding yeast cell cycle. Nature. 2007;448(7156):947–951.

2. MathWorks. Set Optimization Options - MATLAB&Simulink; 2025. `https://www.mathworks.com/help/matlab/math/setting-options.html#bt00l89-1`.

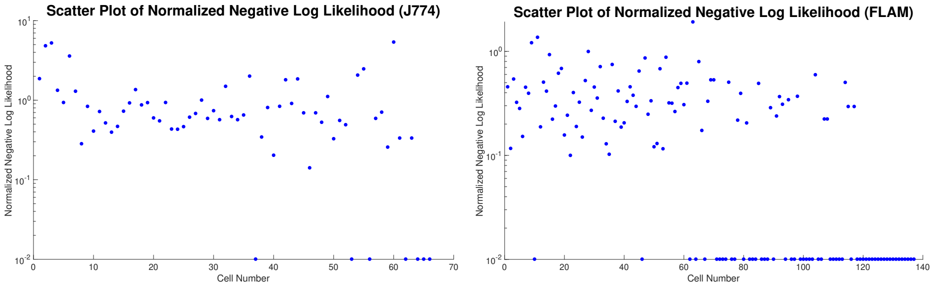


**Fig 13.** *Scatter Plots of Individual-Cell Normalized Negative Log Likelihood Values for Numerical Fits of the Mathematical Model to a Population: The descriptive ability of population-level parameter estimates for individual cell counts is quantified by the normalized negative log likelihood of the population parameters for each individual cell count. Population-level parameter estimation was carried out by minimizing the negative log likelihood of individual observations, under the assumption that the population was homogeneous. In the case of J774 cells, outlier cells 2 and 3 were excluded from the population-level analysis. Cells are arranged by the length of their count data in descending order. Counts are truncated once they achieve a count of six to minimize miscounts due to cell crowding. Cells are arranged by the length of their count data in descending order.*

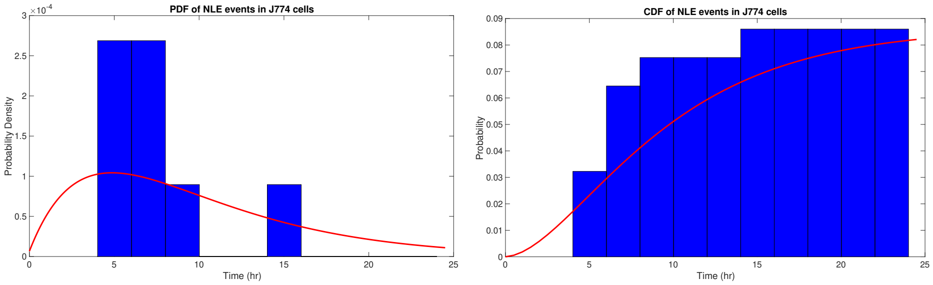


**Fig 14.** *Comparison of Model and Empirical NLE Time Probability Density and Cumulative Density Functions in J774 Cells.*

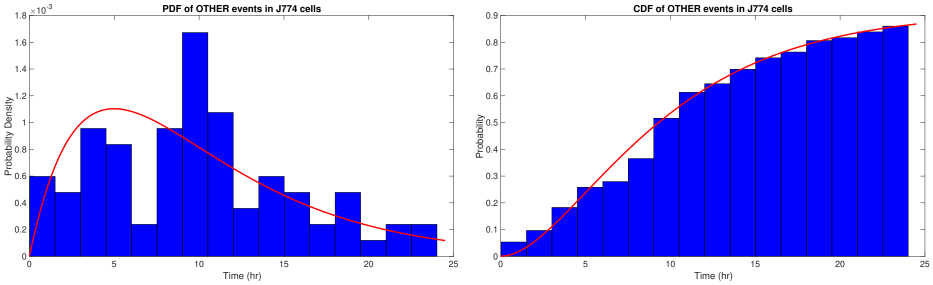


**Fig 15.** *Comparison of Model and Empirical OTHER Time Probability Density and Cumulative Density Functions in J774 Cells.*

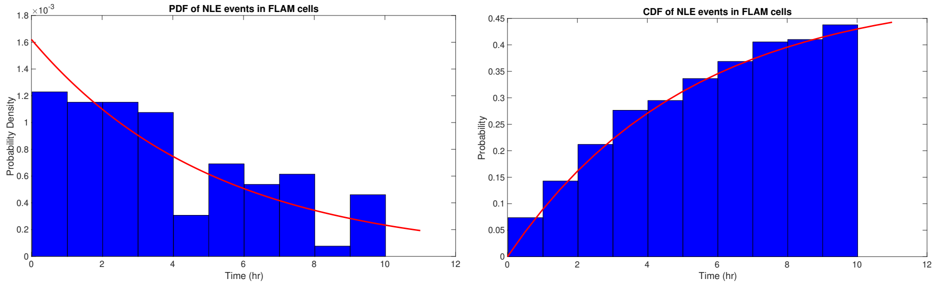


**Fig 16.** *Comparison of Model and Empirical NLE Time Probability Density and Cumulative Density Functions in FLAM Cells.*

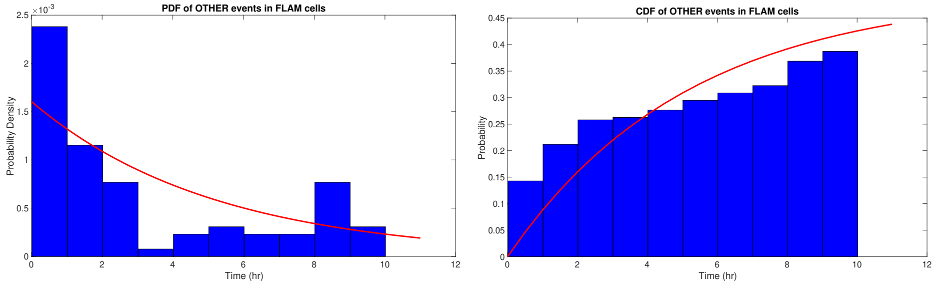


**Fig 17.** *Comparison of Model and Empirical OTHER Time Probability Density and Cumulative Density Functions in FLAM Cells.*
