## Supplementary figures and images for "Intracellular *C. neoformans* infection stimulates increased glycolytic activity in fetal liver-derived alveolar-like macrophages"

### Supplemental figures

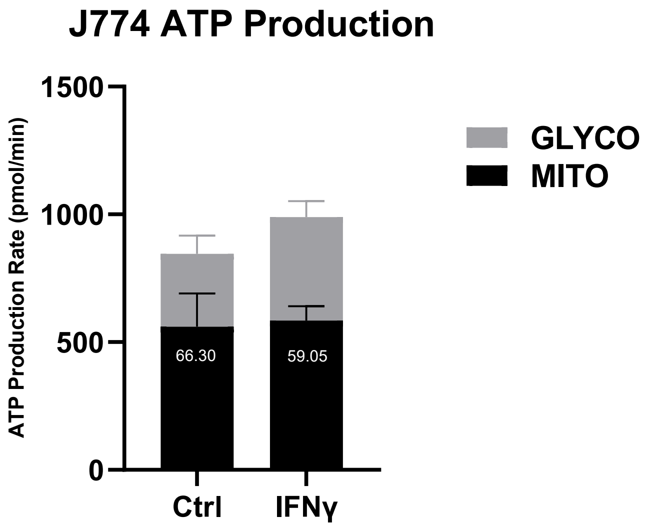

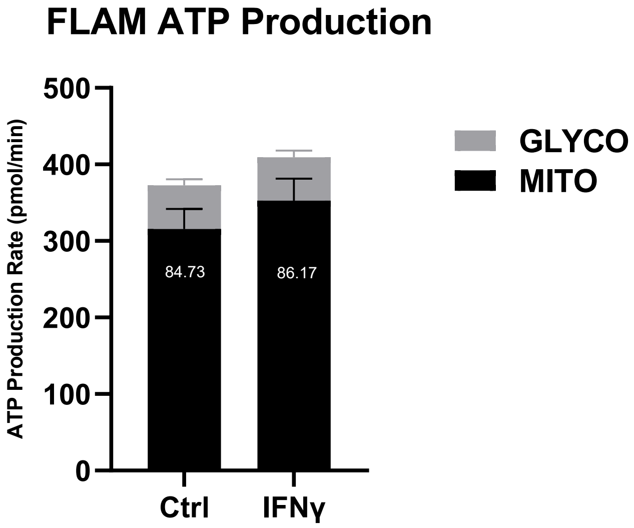

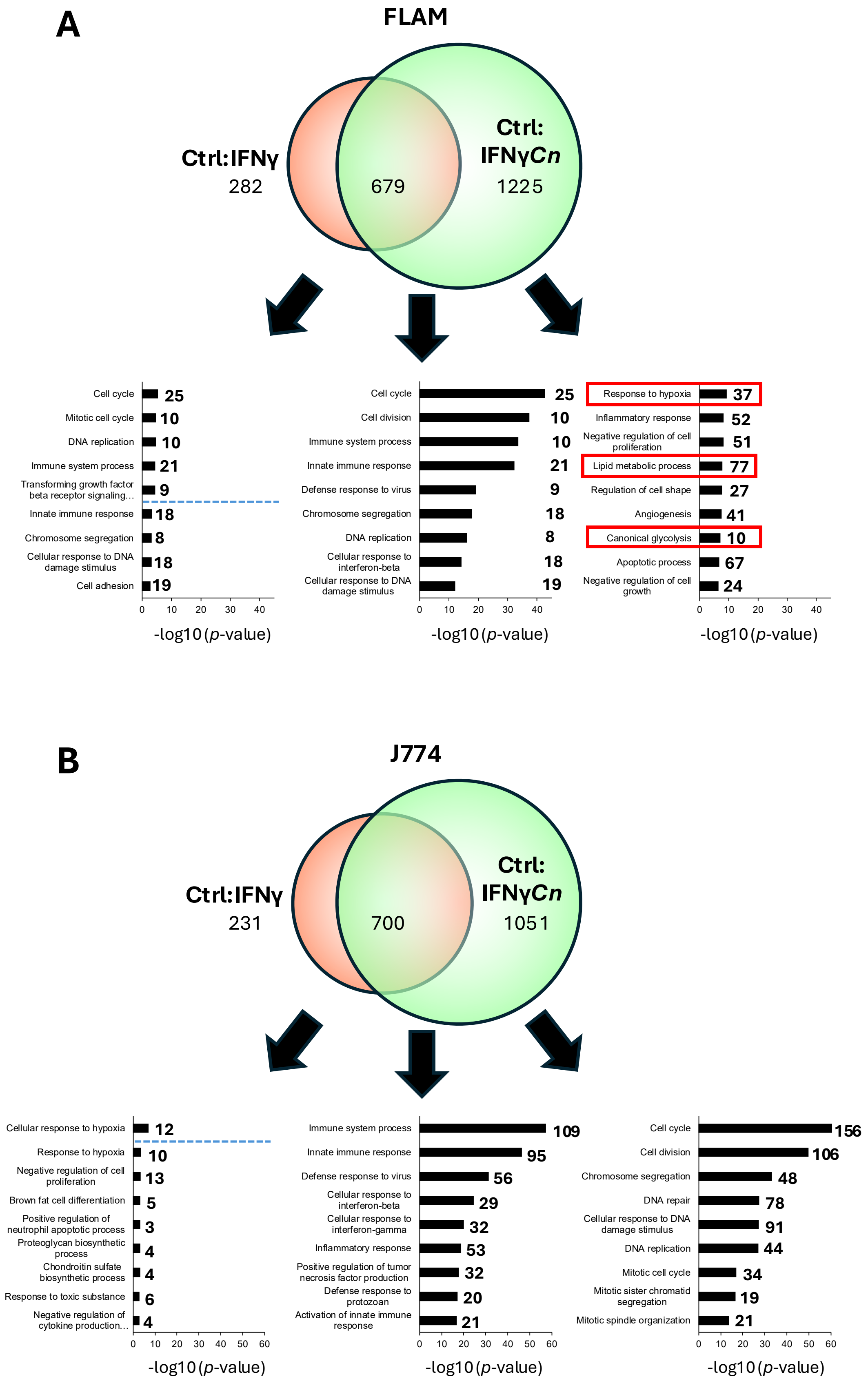

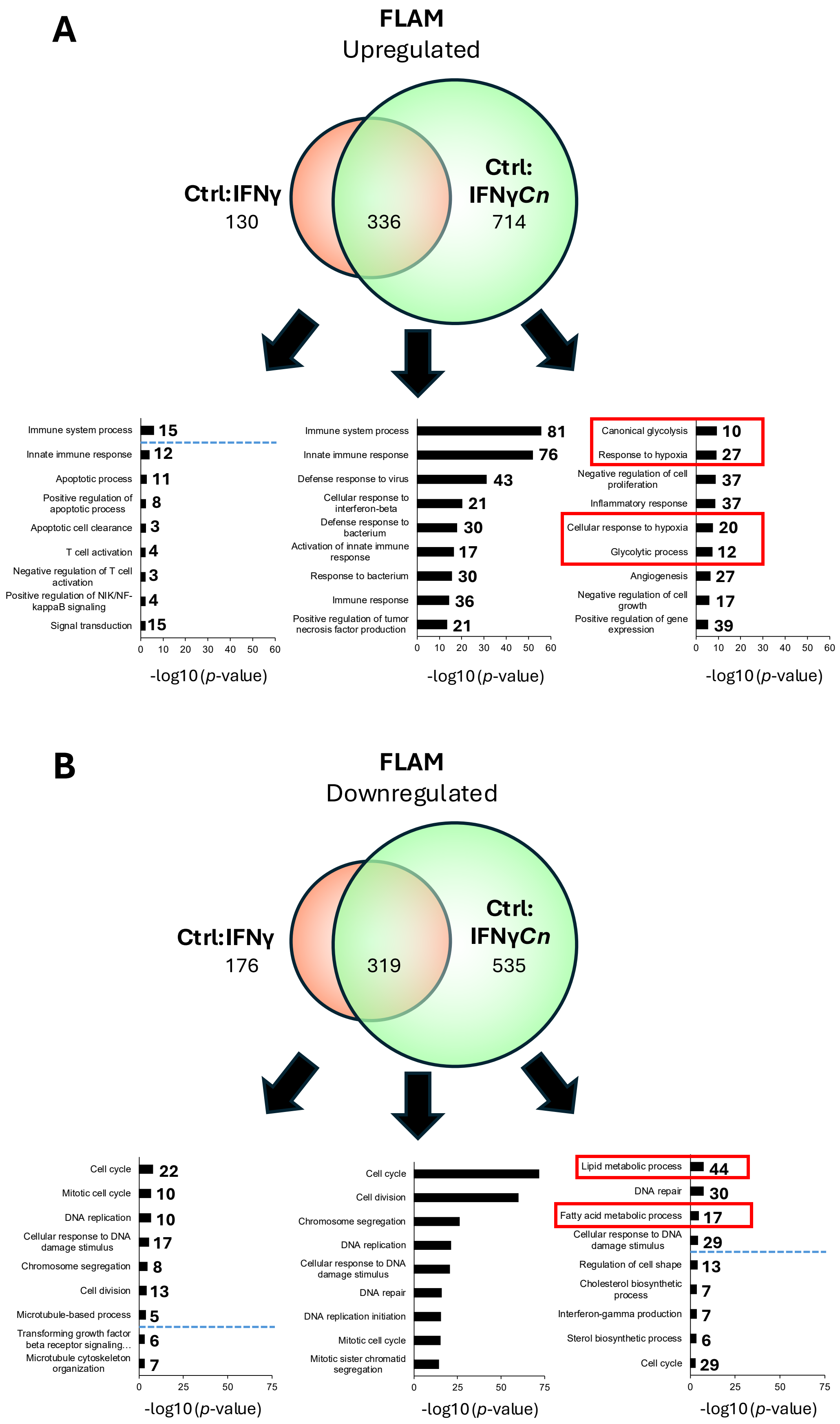

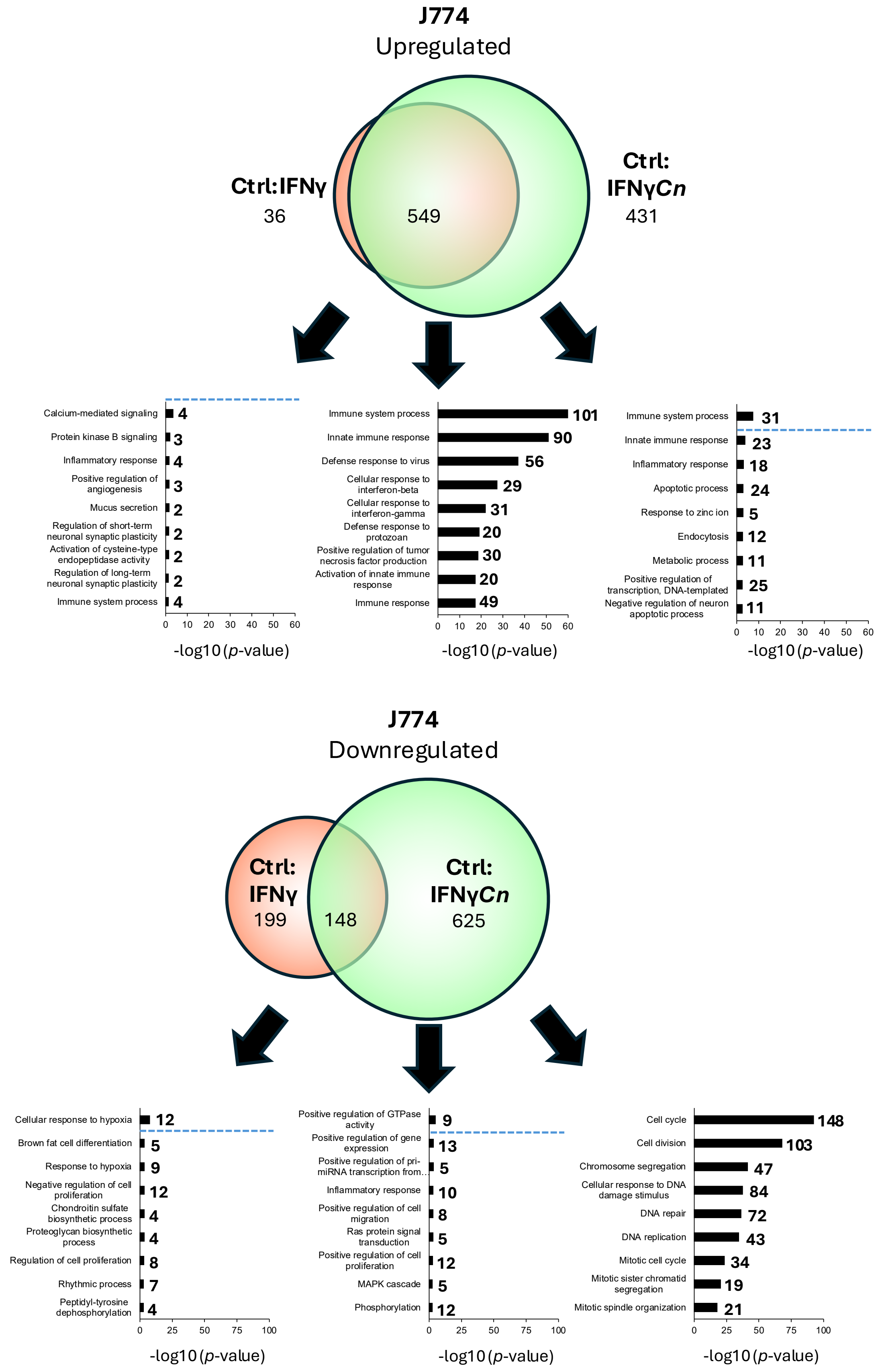

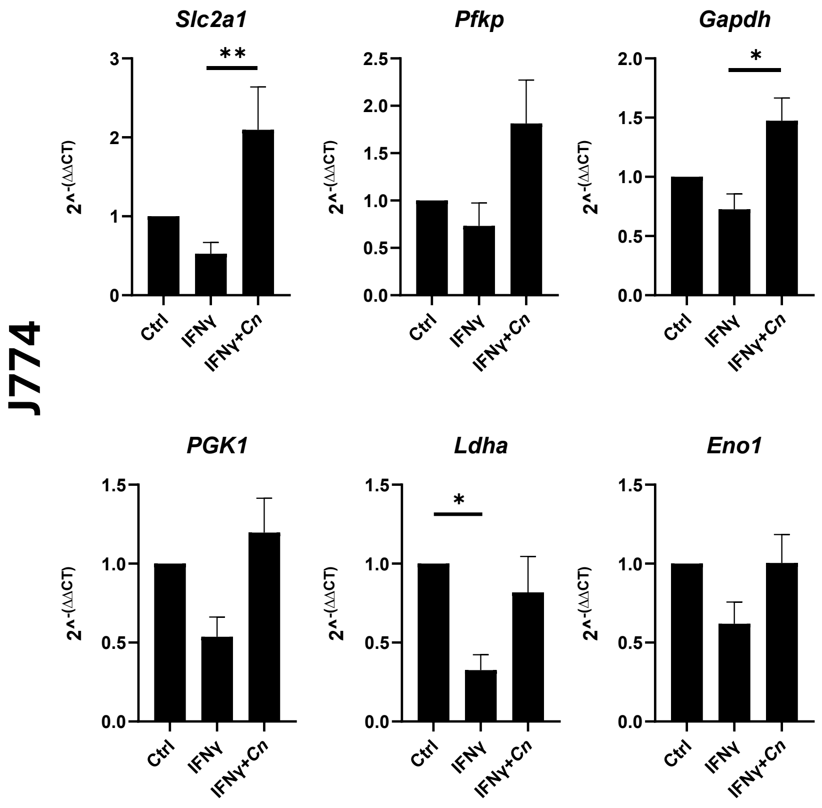

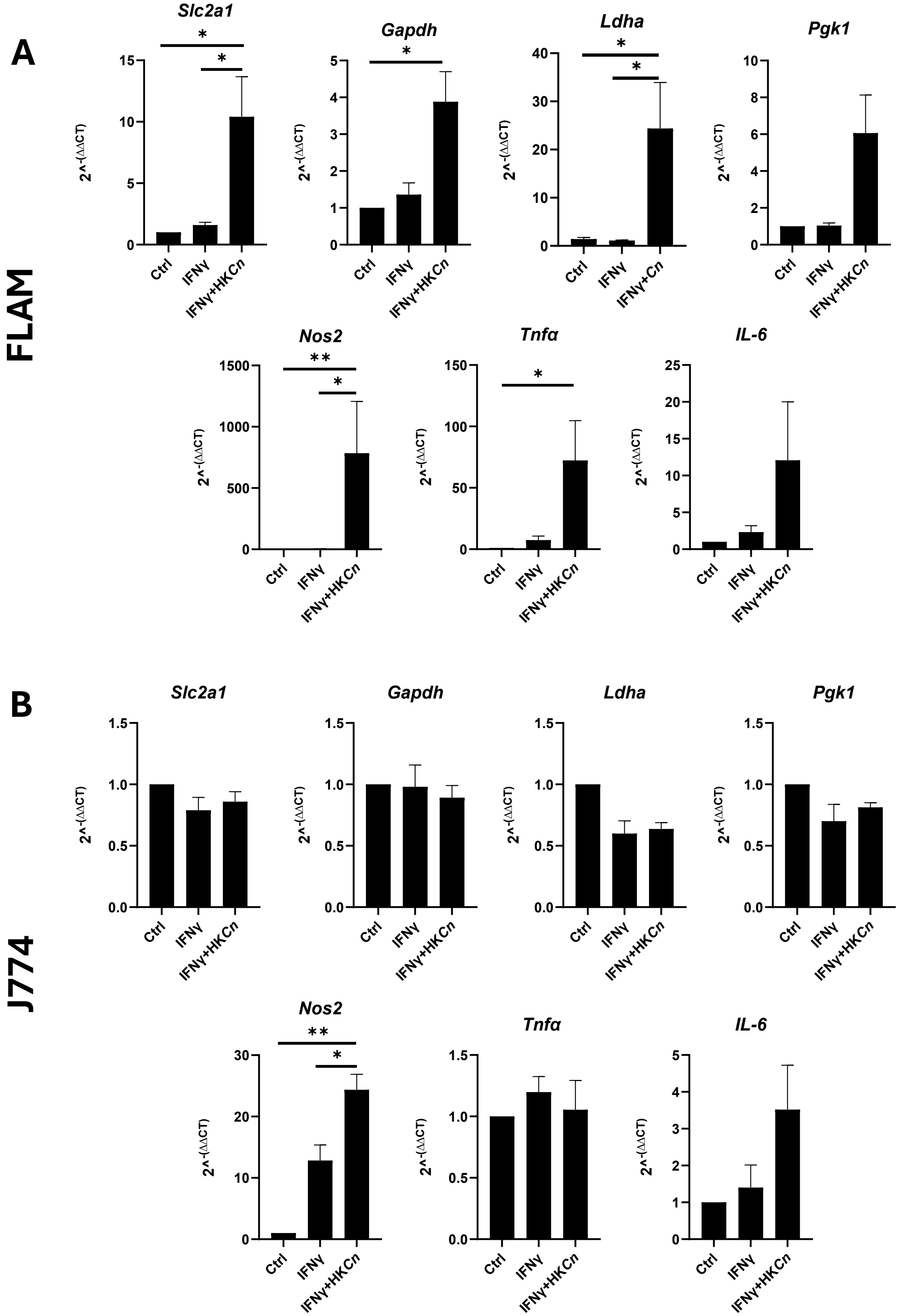

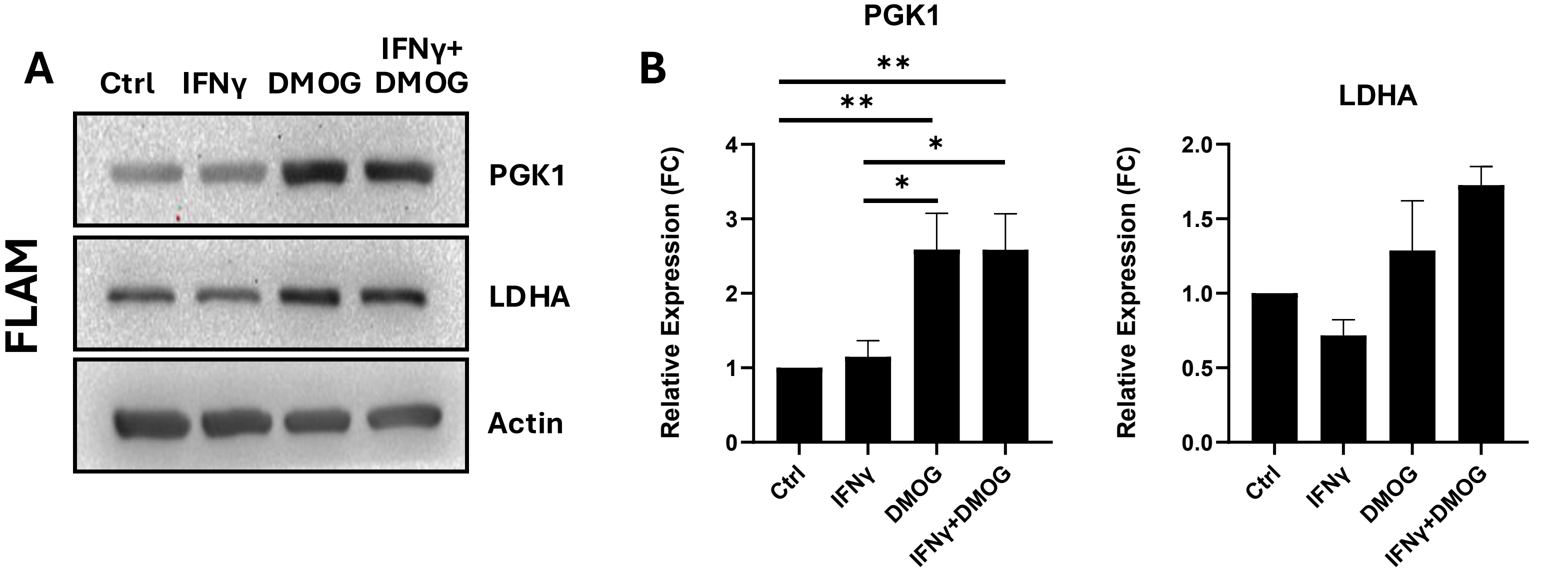
